# Behavioral susceptibility and resilience following early-life stress are associated with sex-dependent locus coeruleus noradrenergic signatures in adulthood

**DOI:** 10.1101/2025.10.11.681820

**Authors:** Dea Slavova, Valentine Greffion, Lionel Granjon, Stéphanie De-Gois, Maud Blaise, Bruno Giros, Elsa Isingrini

**Author notes:** **Corresponding author:** Elsa Isingrini, PhD, 45 rue des Saint Pères, 75006 Paris, France, 06 31 86 45 64.

## Abstract

Child adversity (CA), encompassing emotional, physical, and sexual maltreatment or abuse, affects a substantial number of children worldwide. Moreover, it is the leading predictor of psychiatric disorders such as major depressive disorder (MDD), anxiety, and suicidal behavior. Despite the robust link between CA and psychopathology, individual outcomes vary significantly, with some children demonstrating resilience. Resilience is an adaptive and dynamic process, which mitigates the long-term effects of CA, suggesting potential protective mechanisms that remain underexplored. This study investigates behavioural susceptibility and resilience following early life stress (ELS) and examined whether these outcomes are associated with sex-dependent alteration of the locus coeruleus-norepinephrine (LC-NE) system, a critical modulator of stress, cognition, and emotion. Using a maternal deprivation model combined with limited nesting and bedding, we examined sex-dependent behavioral, physiological, and neurobiological markers associated with ELS outcomes in mice. Behavioral clustering revealed distinct phenotypes: resilient, anxious, and anhedonic-like with sex-specific differences in distribution. Early markers, including body weight and ultrasonic vocalization (USV) patterns, were linked to long-term susceptibility. Neuroanatomical analyses identified sex-specific LC-NE activation patterns associated with resilient and susceptible phenotype, particularly within the caudal-dorsal LC. These findings highlight sex-dependent behavioural trajectories of ELS that are associated with distinct LC-NE activation patterns, offering insights into potential interventions targeting resilience mechanisms in children exposed to CA.

## Introduction

Child adversity (CA), encompassing emotional, physical, sexual abuse and neglect, affects around 25% of children worldwide, with neglect accounting for nearly 80% of cases^1,2^. CA is the strongest predictors of psychiatric disorders, particularly major depressive disorder, anxiety, and suicidal behavior^3–8^. In the U.S., maltreatment-related adverse childhood experiences contribute to 54% and 67% of the population attributable risk for depression and suicide attempts, respectively^9^. CA is also associated with earlier onset, greater severity, and treatment resistance in depression^10–13^. In anxiety, CA worsen outcomes, particularly by exacerbating social anxiety^14,15^. However, outcomes vary: many children adapt successfully despite adversity, a process named resilience. Resilience is an active, dynamic adaptation rather than a simple absence of pathology^16–18^.

From a neurodevelopmental perspective, CA disrupts neurobiological networks involved in threat and reward processing, emotion and cognition^19,20^. Imaging studies showed potential effects of CA on the developing brain at the structural, functional and connectivity level in the anterior cingulate cortex, prefrontal areas, and limbic regions such as the striatum, hippocampus, and amygdala^21–23^. CA also produces lasting effects on stress-related systems, notably the hypothalamic-pituitary-adrenal (HPA) axis and norepinephrine (NE) pathways^24–26^. These changes, persisting into adulthood, may sustain maladaptive coping and psychiatric risk.

While much work has focused on vulnerability mechanisms, research on protective factors predicting positive outcomes is more recent^24^. Resilience may be mediated by larger hippocampal volume, increased white matter integrity in the posterior cingulum, and enhanced prefrontal–limbic functional connectivity^25–29^. Such findings highlight potential neural substrates of resilience, however, how this network is controlled by stress-integrative structures and upstream neuronal mechanisms remains largely unknown.

The locus coeruleus (LC), the largest nucleus producing NE, is projecting throughout the entire brain^30–32^. By modulating cortical and limbic regions, the LC-NE system regulates arousal, stress reactivity, cognition, mood, and emotion^33–35^. It is among the first neurotransmitter systems to develop: in rodents, formation begins at gestational day 10–13 and continues up to three weeks postnatal^36–38^, while in humans, catecholaminergic neurons emerge by five weeks of gestation^39,40^. The LC-NE is thus crucial for brain development as it contributes to brain wiring during critical periods, whereby early life changes in NE signalling can permanently alter emotional and cognitive outcomes. For instance, altered gene expression regulating NE transmission during early development leads to long-term behavioral abnormalities in rodents^41^. Moreover, early life stress (ELS) disrupt the LC-NE system both at the anatomical and functional level. Maternal separation or adolescent stress leads to hyper-activation of the LC, elevated NE release in the paraventricular nucleus of the hypothalamus and reduced α_2_-autoreceptors in the LC^42–46^. Such changes may have lasting impact on cognitive and emotional functions throughout the lifespan, possibly sustaining psychopathological development.

In recent years, the LC-NE system has emerged as a central player in resilience, both in humans^47,48^ and animal models of chronic stress in adulthood^49–51^. Most insights come from the chronic social defeat stress (CSDS) paradigm^52^, widely used to study resilience^50,53,54^. In this model, absence of NE neurotransmission promotes susceptibility, while optogenetic stimulation of LC-NE release in the VTA reversed susceptibility to induce resilience^49^. Complementary imaging studies show LC-NE changes in post-traumatic stress disorders (PTSD), where heightened LC activity predicts greater symptom severity^55,56^. Despite these evidences, the relationship between LC-NE system activity and resilient vs susceptible outcomes following ELS remains underexplored.

Given its role in modulating key brain regions involved in both resilience networks and the impact of CA, we hypothesized that long-term behavioral outcomes following ELS would be associated with distinct patterns of LC-NE activity. The objective of this study was to develop a maternal deprivation model combined with limited nesting and bedding as a paradigm of ELS, enabling segregation of susceptible and resilient mice based on their long-term anxio-depressive phenotypes. We further investigated whether early and late adaptive mechanisms contribute to resilience. Early emotional and physiological predictors were assessed through body weight and ultrasonic vocalization (USV), while long-term outcomes were examined via corticosterone levels and LC-NE neuronal activity in adulthood.

## Methods and materials

### Animals

The Paris Cité University ethics committee (CEEA 40) approved all procedures, complying with the European Directive 2010/63/EU. C57BL/6J mice (Janvier Labs) were housed under standard laboratory conditions (22±1 °C, 60% humidity, 12h light/dark cycle) with food and water *ad libitum*.

8-week-old breeders were paired (two females/one male). Pregnancy was monitored twice weekly, and males were removed upon confirmation. Pregnant females were single-housed four days before parturition.

### Early life stress (ELS) paradigm

ELS combined maternal deprivation (MD) with limited nesting (½ of standard material) and bedding (1/10 of standard bedding). Birth was designated as P0. From P2 to P14, pups were separated from dams for 3h/day. During separation, pups were placed individually on a heating pad in a different room to prevent communication. From P14 to P21, pups were reared under standard housing conditions. Control (CTL) litters remained undisturbed under standard conditions.

Body weight was measured at P2, P5, P7, P12, and P14, and weekly thereafter. At P21, pups were group-housed by sex and condition.

### Ultrasonic vocalizations (USV) recording

Pups were recorded individually for 5 min at 9:00 a.m. in a styrofoam box, once daily for CTL and before/after MD for ELS, at P2, P5, P7 and P12^57,58^. For afternoon born litters, P2 recordings were time-adjusted^58^.

Recordings used an Avisoft UltraSoundGate 116Hb system connected a CM16 microphone (Avisoft 40011; 250 kHz, 1024 FFT, 16-bit). Gain was set to avoid overload. Analyses were performed using Hybrid Mouse (MATLAB)^59^.

### Maternal Behavior

Maternal behavior was assessed at P2, P7, and P14 before and after MD. After 30 min habituation, dams were video-recorded for 15 min. Behavior was quantified as total nest time.

### Splash Test

At P21, a 10% sucrose solution was sprayed on the dorsal coat and 5-min homecage grooming was recorded as a self-care measure^60^.

### Anxiety and depressive-like behavior in offspring

At adulthood (8–12 weeks), CTL and ELS mice underwent a battery of paradigms to increase stressfulness.

#### Three chamber test (3CH)

Sociability was assessed using a Plexiglas box (68 × 22 × 24 cm) with two side (28 × 22 × 24 cm) and one central (12 × 22 × 24 cm) chambers^61^. After 10 minutes habituation with empty wire-mesh, an unfamiliar age- and sex-matched mouse was placed under one mesh, the other left empty. Social interaction was quantified as time within a 2-cm radius around each mesh. A social interaction index was calculated: [Time_social / (Time_social + Time_empty)].

#### Novelty-suppressed feeding test (NSF)

The NSF test was conducted in a 45cm^3^ open field with bedding, after 24h food deprivation. Latency to eat a pellet on a 12.5cm white paper in the center was recorded (max 5 min), followed by 3 min homecage food intake measurement.

#### Sucrose preference test (SPT)

Mice were individually housed overnight (from 19h-9h) for two days with two water bottles. From days 3–6, one bottle contained 1% sucrose (SB) and the other water (WB), with positions alternated daily. Consumption was measured through bottle weight, and mean sucrose preference across days calculated: [SB/(SB+WB) ×100].

#### Forced swim test (FST)

Mice were placed in a glass cylinder (25 cm high, 9 cm diameter) filled with 21–23°C water. Immobility was recorded during 6 min as a measure of resignation, defined as passive floating with minimal movements to keep the head above water.

### Estrous cycle

Following behavioral tests (except SPT), vaginal smears were collected with 20µL sterile 0.9% NaCl. Samples were placed on slide, stained 1 minute with crystal violet^62^ and estrous stages visually identified.

### Corticosterone dosage

Blood (100 µL) was collected from the submandibular vein at baseline (30 min pre-FST), immediately after, and 1h30 post-FST. Samples clotted 30 min at room temperature (RT) were centrifuged (5,000 rpm, 10 min, 4°C) and supernatants stored at –20°C. Corticosterone levels were measured using the DetectX EIA Kit (Arbor Assays) and expressed as mg/µL. Percentage changes were calculated:

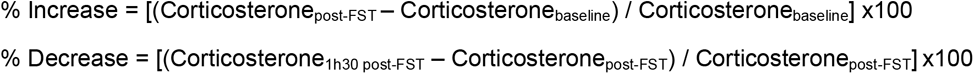

### Tissue preparation and immunohistochemistry

1h30 post-FST, mice were perfused with 0.9% NaCl followed by Antigenfix (Diapath P0014). Brains were post-fixed overnight in Antigenfix, stored in 0.1M PBS, embedded in 2% agarose and cut into 40-µm-thick coronal sections (Leica vibratome VT1000E). Free-floating sections were stored at –20°C in cryoprotective solution (30% glycerol, 30% ethylene glycol, 40% 0.1M PBS).

Sections were incubated 30 min at RT in 1% NaBH4 and transferred in blocking solution (0.1M PBS, 0.3% Triton X-100, 5% donkey serum) for 30 min at RT. Section were incubated overnight at 4°C under agitation with primary antibodies: Guinea pig anti-c-Fos (1:2500, Synaptic Systems 226004) and Mouse anti-TH (1:2000, Sigma-Aldrich MAB318) or Rabbit anti-TH (1:4000, Abcam 117112). After washes, sections were incubated for 2h at RT with the secondary antibodies: Cy3-conjugated donkey anti-guinea pig (1:500, Jackson Laboratory 706-165-148) and Alexa Fluor 488 donkey anti-mouse (1:500, Invitrogen A21202) or Alexa Fluor 488 donkey anti-rabbit (1:4000, Jackson Laboratory 711-546-152). Sections were mounted on superfrost slides, and coverslipped with Fluoromount-G™ mounting medium.

### Image acquisition and analysis

Images were acquired with a Zeiss® LSM710 confocal microscope with a ×20 objective lens. Cell counting was performed manually in ImageJ^63^. TH⁺, c-Fos⁺ and TH⁺/c-Fos⁺ cells were quantified in six sections per mouse across LC subregions. Cell densities were expressed as positive cells/µm². Proportion of activated NE cells was calculated:

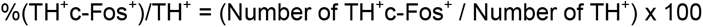

### Statistical analysis

Normality was assessed with Shapiro-Wilk test or qqplot inspection and homogeneity of variance with Levene’s test. When assumptions were met, Student’s t-test, one- or two-way ANOVA were used, followed by Tukey or Bonferroni post-hoc tests. For unequal variances, Welch (W) correction was applied when possible. If normality was violated, Mann-Whitney U (U) test was used. For repeated measures ANOVA, Mauchly’s test assessed sphericity; Greenhouse-Geisser correction was applied when violated. Mahalanobis distance and k-means clustering were computed with MATLAB and R scripts. Results are expressed as mean ± SEM (standard error of the mean) with significance set at p<0.05.

## Results

### ELS impact on maternal behavior and offspring development (Figure 1A)

Maternal behavior, evaluated by nest time (P2, P7, P14), did not differ between CTL and ELS dams (F_1.5,27.2_=2.988, p=0.08; **Figure 1B**). After weaning, corticosterone level (t_10_=0.237, p=0.817) and grooming duration (t_10_=-0.663, p=0.522) were unchanged (**Figure 1C**).

**Figure 1.**
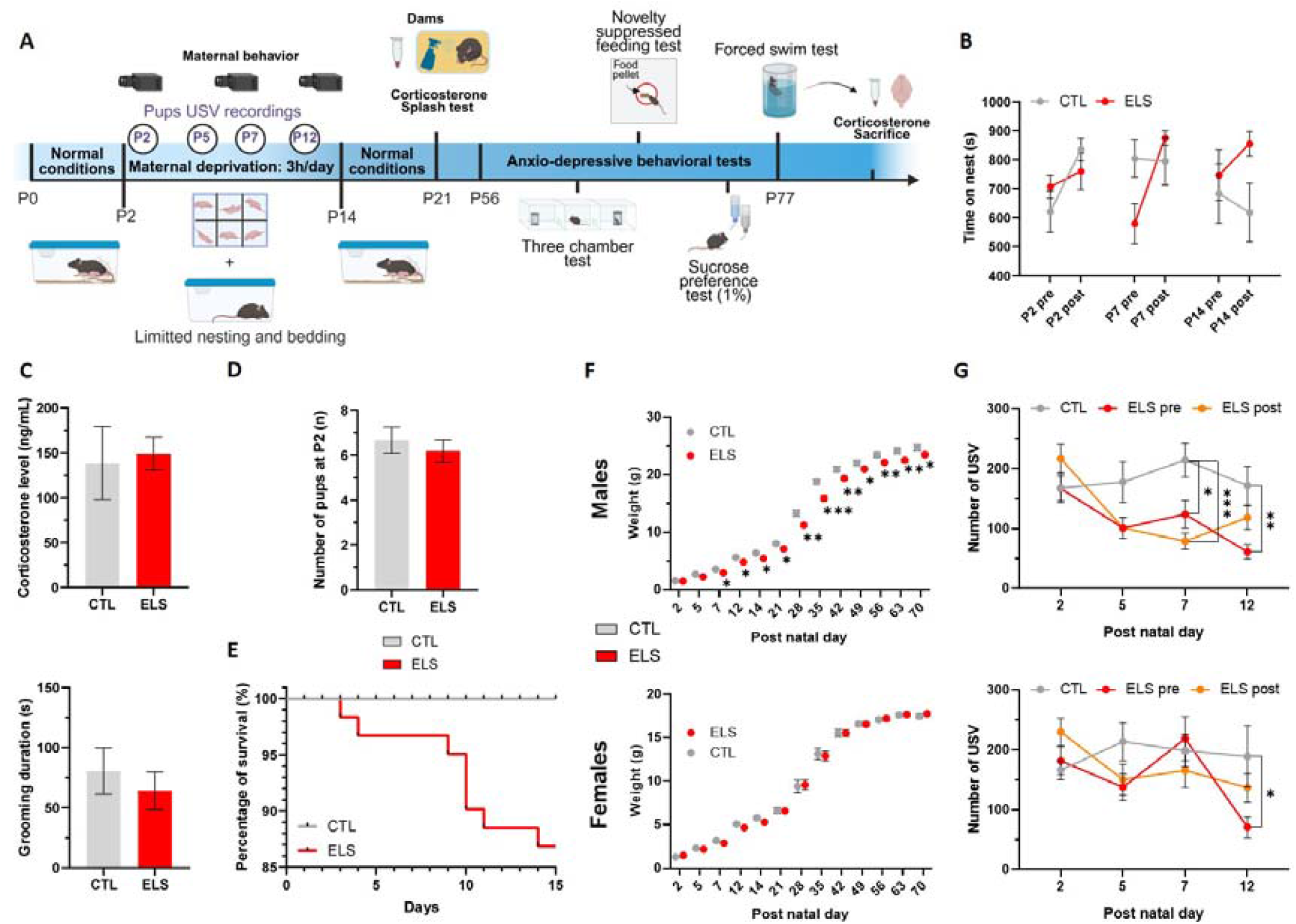
Consequences of the maternal deprivation model combined with limiting nesting/bedding on maternal behavior, pups development and adult behaviour. **A.** Timeline of the ELS paradigm applied from P2 to P14, combining maternal deprivation (3h/day) with limited nesting (1/2) and bedding (1/10) materials. Starting at P56, anxio-depressive behaviors were evaluated in male and female mice. At P77, mice were sacrificed 1h30 minutes following the FST. Created in BioRender. Sla, D. (2025). https://BioRender.com/jgluigr. **B.** Maternal behavior was measured at P2, P7 and P14 before and after the deprivation. For the time spent on the nest, no significant interaction between post-natal day, time and condition was found (F_2,72_=2.88, p=0.063). **C. (*Top)*** Corticosterone level measured at P21 in dams, 30 minutes before the splash test, show no significant difference between groups (t_13_=1.37, p=0.193). **(*Bottom*)** Results of the splash test at P21 reveal no significant differences between CTL and ELS dams in total grooming duration (t_13_=0.23, p=0.818). **D.** The number of pups at birth did not differ significantly between CTL and ELS groups (t_18_=0.637, p=0.532). **E.** ELS survival rate dropped to 87.5% compared to 100% in CTL. **F.** The body weight (g) of male **(*Top*)** and female **(*Bottom*)** mice was analyzed from P2 to P70. In males, a significant interaction between postnatal day and condition (F_3.376,118.171_=7.53, p<0.001) was found. Bonferroni post hoc tests revealed that ELS (n=25) had significantly lower weights compared to CTL (n=18) from P7 to P70 (P7: t=2.21, p = 0.034; P12: t=2.30, p=0.027; P14: t=2.44, p=0.02; P21: t=2.25, p=0.031; P28: t=3.20, p=0.003; P35: t=4.94, p<0.001; P42: t=2.91, p=0.006; P49: t=2.23, p=0.032; P56: t=3.11, p=0.004; P63: t=2.83, p=0.008; P70: t=2.22, p=0.033). In females, no interaction was detected between postnatal day and condition (F_2.1,67.2_=0.67, p=0.52); CTL: n= 19, ELS n= 21). **G.** USVs were measured in both male **(*Top*)** and female **(*Bottom*)** pups at P2, P5, P7 and P12 before and after the deprivation in ELS. In males, a significant interaction was found between condition and postnatal day (F_3,195_=3.86, p=0.001). Bonferroni post hoc tests showed that USVs were significantly reduced in ELS compared to CTL at P7 pre- and post-deprivation (Pre: t=2.89, p=0.01; post: t=4.32, p<0.001) and at P12 post-deprivation (t=3.6, p=0.002). In females, a significant interaction between postnatal day and condition was also observed (F_5.03,143.34_=2.399, p=0.04), with a significant reduction in USVs pre-deprivation at P12 (t=2.56, p=0.039). Data are presented as mean ± SEM. Two-way ANOVA was followed by Bonferroni post hoc tests. The Greenhouse-geisser correction was applied if sphericity was violated. Statistical significance is indicated as: *p < 0.05, **p < 0.01, ***p < 0.001. Abbreviations: CTL, control; ELS, early-life stress; P, post-natal day; USV, ultrasonic vocalisation; FST, Forced swim test; NSF, novelty supressed-feeding test; 3CH, 3-chamber test; SPT, sucrose preference test.

Regarding offspring, litter size at P2 was similar across groups (U=43, p=0.638), but survival dropped from 100% in CTL to 87.50% in ELS (**Figure 1D-E)**. In males, body weight reduction emerged at P7 and persisted until P70 (F_3.4,118.2_=7.53, p<0.001, P7: t=2.21, p=0.034; P12: t=2.30, p=0.027; P14: t=2.44, p=0.02; P21: t=2.25, p=0.031; P28: t=3.20, p=0.003; P35: t=4.94, p<0.001; P42: t=2.91, p=0.006; P49: t=2.23, p=0.032; P56: t=3.11, p=0.004; P63: t=2.83, p=0.008; P70: t=2.22, p=0.033). Female weight was unaffected (F_2.1,67.2_=0.67, p=0.52; **Figure 1F**). USV emissions were reduced in ELS male at P7 pre- and post-deprivation and P12 pre-deprivation (F_3,195_=3.86, p=0.001; P7pre: t=2.89, p=0.01; P7post: t=4.32, p<0.001; P12pre: t=3.6, p=0.002). In females, ELS reduced USV at P12 pre-deprivation (F_2.515,143.34_=2.399, p=0.04, P12: t=2.56, p=0.039; **Figure 1G**).

### Long-term ELS behavioral consequences: toward susceptibility and resilience

Anxio-depressive paradigms were performed from P56 to P77 (**Figure 1A**). The NSF test assessed anxiety-like behavior, the 3CH test sociability, and the sucrose preference test anhedonia. The FST, evaluated resignation, also served as an acute stressor to assess corticosterone response and LC-NE c-Fos activation. Given that raw data analysis does not account for cohort effect (**Supplementary Figure 1-2**), data were normalized using z-score, calculated within CTL groups by cohort and sex. Subsequent analyses were conducted using z-score.

No significant differences emerged between CTL and ELS groups in social (3CH M: W_41_=1.25, p=0.22; F: U_37_=187, p=0.97), anxiety-like (NSF, M: U_41_=189, p=0.38; F: U_37_=175, p=0.7), and resignation (FST, M: t_41_=0.32, p=0.75; F: t_37_=0.46, p=0.64) behaviors. However, ELS females displayed increased anhedonia, measured by reduced sucrose preference compared to CTL (W_30.387_=3.04, p=0.005). This effect was absent in male (U_41_=249, p=0.57; **Figure 2A-B)**. ELS effects on corticosterone were sex-dependent, with decreased baseline level in male (t_41_=3.3, p=0.002), and no effects in females (**Supplementary Figure 3A-B**), while LC-NE functional anatomy was unchanged in both sexes (**Supplementary Figure 4A-B**).

**Figure 2.**
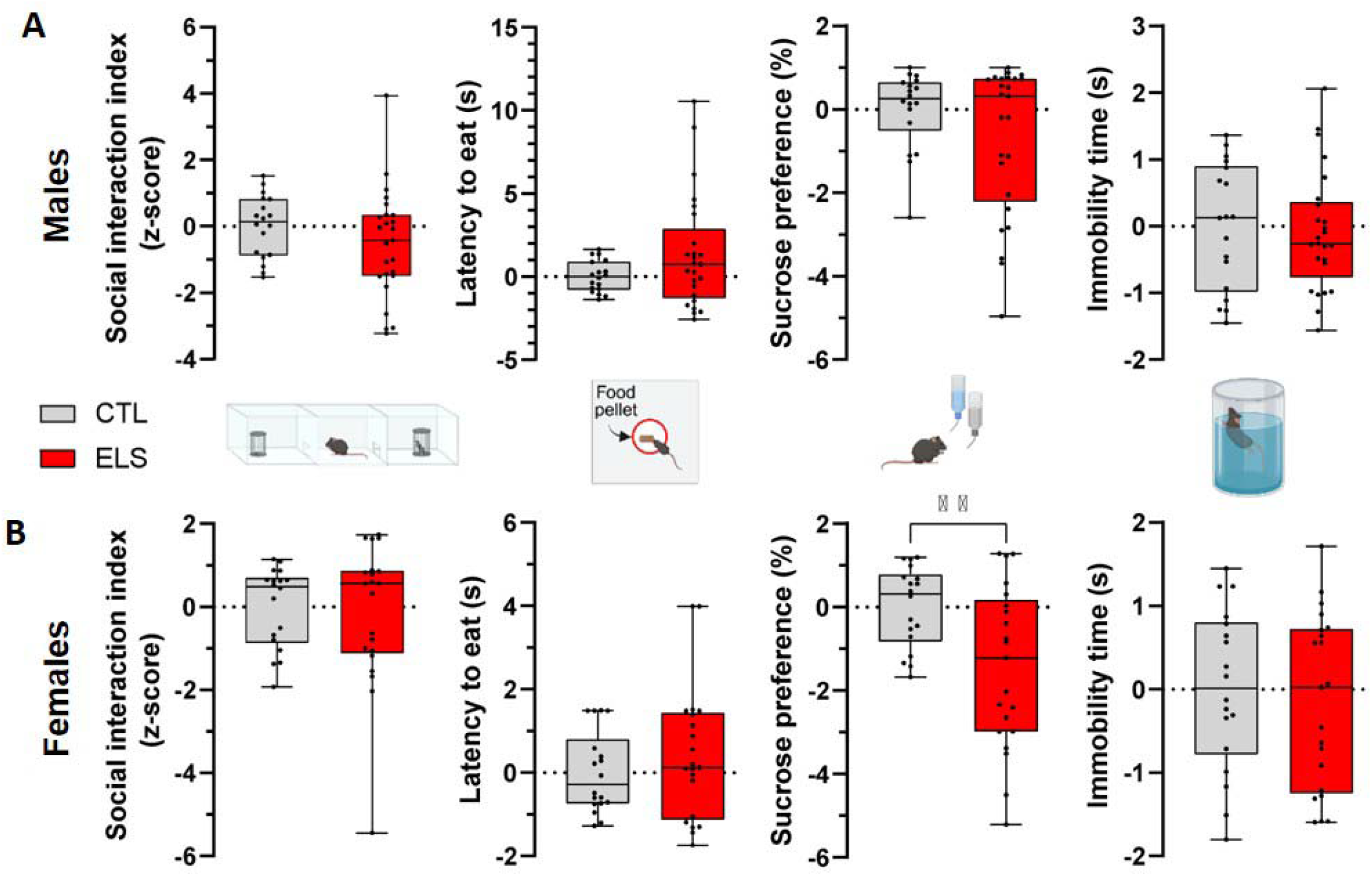
Consequences of the maternal deprivation model combined with limiting nesting/bedding on adult behaviour. **A.** In males, no significant differences between CTL and ELS groups were revealed for the social interaction index in the 3CH (t_41_=1.25, p=0.22), the latency to eat in the NSF (U_41_=189, p=0.38), the percentage of sucrose preference (SPT, U_41_=249, p=0.57) and the immobility time in the FST (t_41_=0.32, p=0.75). **B.** In females, no significant differences between CTL and ELS groups were revealed for the social interaction index in the 3CH (U_37_=187, p=0.97), the latency to eat in the NSF (U_37_=175, p=0.7) and for the time immobile in the FST (t_37_=0.46, p=0.64). However, the percentage of sucrose preference was significantly reduced in ELS group compared to CTL (W_30.387_=3.04, p=0.005). Data are presented as mean ± SEM. Statistical significance is indicated as: *p < 0.05, **p < 0.01, ***p < 0.001. Abbreviations: CTL, control; ELS, early-life stress; P, post-natal day; USV, ultrasonic vocalisation; FST, Forced swim test; NSF, novelty supressed-feeding test; 3CH, 3-chamber test; SPT, sucrose preference test.

Oestrus cycle did not influence social interaction or immobility (3CH: F_1,35_=0.51, p=0.48; FST: F_1,35_=2.05, p=0.16). However, in the NSF, ELS females in proestrus/estrus (P/E) cycle showed longer latency to eat than CTL (F_1,35_=5.72, p=0.022; P/E: t_35_=-2.51, p=0.017), an effect absent in diestrus/metestrus (D/M) (t_35_=0.69, p=0.5; **Supplementary Figure 5A**). Although cycle had no effect on corticosterone, D/M females displayed higher LC-NE activation density and proportion than P/E females, regardless of CTL or ELS status (**Supplementary figure 6A–B**).

To further distinguish susceptible and resilient individuals, we applied two analyses: Mahalanobis distance and k-means clustering, using latency to eat (NSF), social interaction index (3CH), % sucrose preference (SPT), and immobility time (FST), separately in males and females. Mahalanobis distance quantified each mouse’s deviation from the CTL distribution, with greater distance reflecting higher susceptibility. In parallel, k-means clustering assigned individuals to clusters by minimizing within-clusters variances.

### 1. The Mahalanobis distance analysis

Mahalanobis distance analysis, combined with correlation, identified bio-behavioral markers of susceptibility (**Figure 3A)**. While the Mahalanobis distance was significantly higher in ELS than CTL groups, oestrus cycle stage in females had no effect (**Supplementary Figure 5B**).

**Figure 3.**
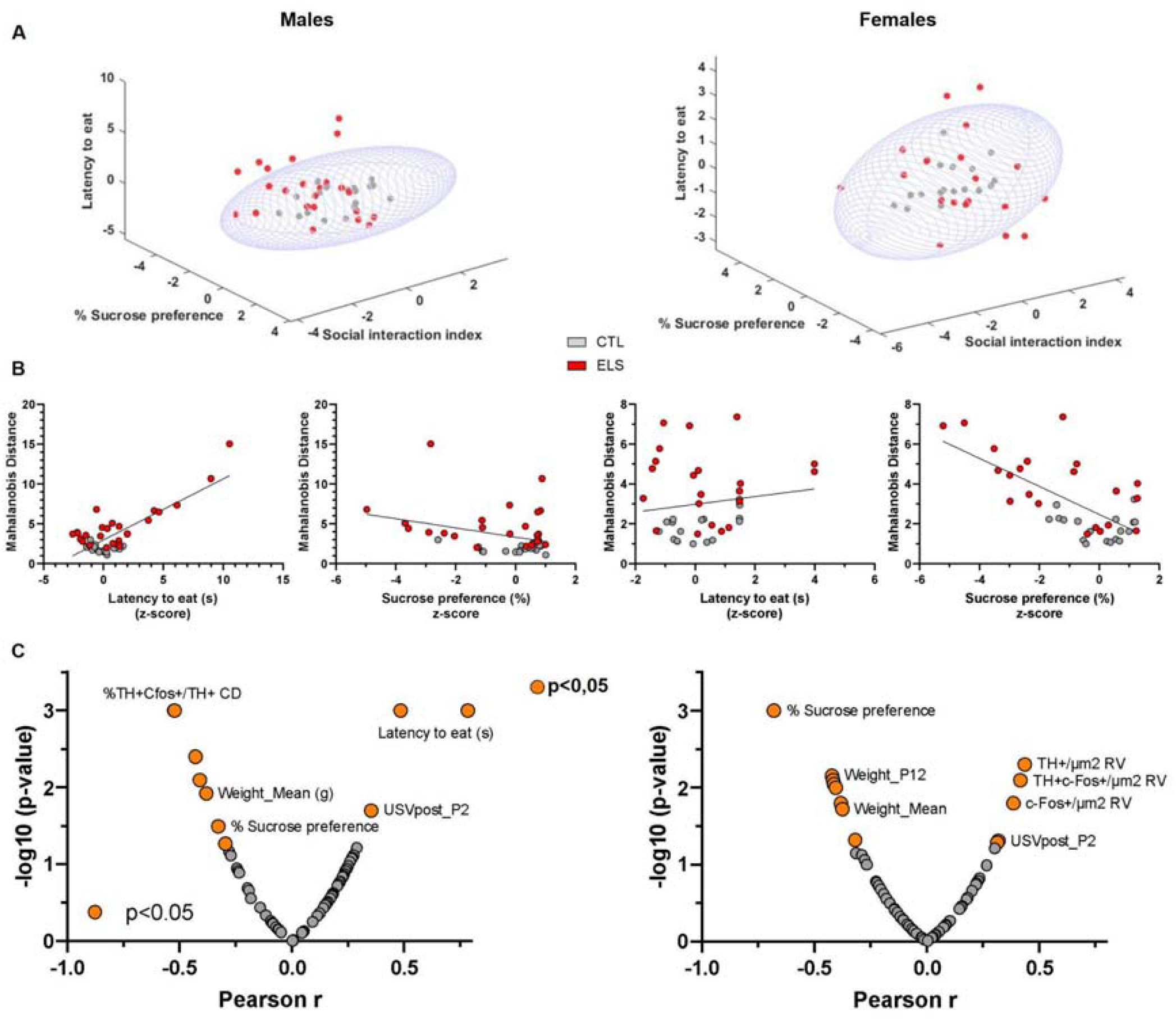
Mahalanobis distance analysis reveals long-term behavioral susceptibility to ELS. **A.** Susceptibility to ELS, quantified by the Mahalanobis distance, is shown for males (***Left***) and females (***Right***), based on four behavioral measures: social interaction index in the three-chamber test (3CH), latency to eat in the novelty-suppressed feeding test (NSF), sucrose preference (%), and immobility time in the forced swim test (FST). **B.** Pearson correlations between Mahalanobis distance and behavioral measures. In males (***Left***), Mahalanobis distance was negatively correlated with latency to eat (R = –0.78, p < 0.001) and sucrose preference (R = –0.33, p = 0.032). In females (***Right***), a significant negative correlation was found with sucrose preference (R = –0.679, p < 0.001), while no association was observed with latency to eat (R = 0.146, p = 0.376). **C.** Volcano plots illustrating –log(p) values as a function of Pearson’s r for males (***Left***) and females (***Right***). Parameters include body weight (P2–P70 and mean weight), USV (P2–P12), mean pre- and post-deprivation, corticosterone (baseline, post-FST, 1h30 post-FST, % increase, % decrease), and LC activity markers (density of TH^+^c-Fos^+^ cells and %TH^+^c-Fos^+^/TH^+^ across the rostro-caudal and dorso-ventral axes). Abbreviations: CTL, control; ELS, early-life stress; P, post-natal day; USV, ultrasonic vocalisation; FST, Forced swim test; NSF, novelty suppressed-feeding test; 3CH, 3-chamber test; SPT, sucrose preference test.

#### Behavioral correlates

In both sexes, higher Mahalanobis distance correlated with lower sucrose preference (M: R=-0.33, p=0.032; F: R=-0.679, p<0.001). In males, it was also associated with longer latency to eat in the NSF (R=-0.78, p<0.001), a link absent in females (R=0.146, p=0.376; **Figure 3B**). No correlation emerged with the social interaction index (M: R=-0.235, p=0.13; F: R=-0.205, p=0.21) or the immobility time (FST, M: R=0.053, p=0.74; F: R=-0.118, p=0.47).

#### Early-life predictors

Lower mean body weight from P2 to P14 predicted higher susceptibility in both males (R=-0.38, p=0.012) and females (R=-0.37, p=0.020). Post-deprivation USV emissions at P2 were positively correlated with susceptibility in both sexes (M: R=0.35, p=0.020; F: R=0.32, p=0.049), suggesting heightened vocalization after the first deprivation as an early predictor. Otherwise, in both sexes USV emissions were unaffected in susceptibility (**Figure 3C**).

#### Late-onset markers

Corticosterone responses to stress showed no association with Mahalanobis distance in either sex (**Figure 3C**) at baseline (M: R=0.061, p=0.70; F: R=0.27, p=0.10), post-FST (M: R=–0.25, p=0.11; F: R=–0.19, p=0.24), or 1h30 post-FST (M: R=–0.094, p=0.55; F: R=0.074, p=0.65). Similarly, neither the relative increase (M: R=–0.087, p=0.58; F: R=0.23, p=0.16) nor the subsequent decrease (M: R=–0.047, p=0.77; F: R=0.20, p=0.22) correlated with susceptibility. However, anatomical analyses revealed sex-specific LC-NE activation patterns (**Figure 3C**). In males, reduced activation of NE neurons in the caudal-dorsal LC predicted susceptibility (%TH^+^c-Fos^+^/TH^+^; R=–0.52, p<0.001), while susceptible females exhibited a significant increased activation in the rostral-ventral LC (TH^+^c-Fos^+^/µm²; R=0.41, p=0.008).

### 2. K-means clustering analysis to reveal resilience

K-means clustering of ELS mice allowed identifying three clusters in both sex: **anxious-like (ANX)**, characterized by increased NSF latency (M: F_3,12.9_=4.33, p=0.025; CTL *vs* ANX: t=–4.22, p<0.001; F: F_3,35_=6.99, p<0.001; CTL *vs* ANX: t=–4.02, p<0.001) and reduced social interaction (M: F_3,39_=7.87, p<0.001; CTL *vs* ANX: t=3.8, p=0.002; F: F_3,35_=4.43, p=0.01; CTL *vs* ANX: t=2.71, p=0.03); **anhedonic-like (ANH)**, defined by reduced sucrose preference (M: F_3,39_=19.71, p<0.001; CTL vs. ANH: t=6.98, p<0.001; F: F_3,35_=31.52, p<0.001; CTL vs. ANH: t=8.75, p<0.001); and **resilient (RES)**, showing no behavioral alterations relative to CTL (**Figure 4A-B**).

**Figure 4.**
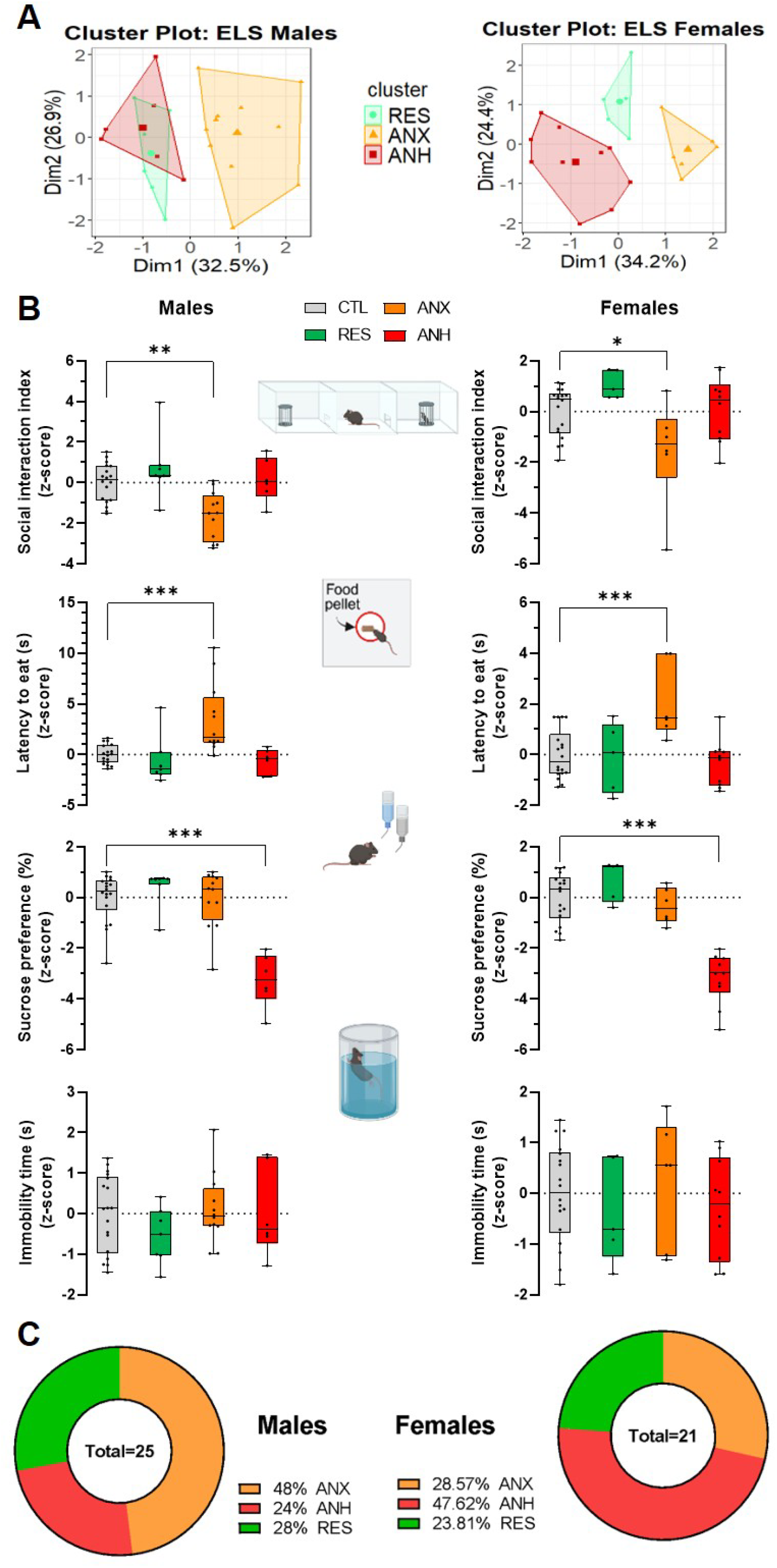
K-means clustering analysis reveals long-term behavioral susceptibility and resilience to ELS. **A.** Cluster plots from k-means analysis (k = 3) in males (***Left***) and females (***Right***), based on four behavioral measures: social interaction index in the three-chamber test (3CH), latency to eat in the novelty-suppressed feeding test (NSF), sucrose preference (%) in the sucrose preference test (SPT), and immobility time in the forced swim test (FST). Three clusters were identified in both sexes: ANX (anxiety-like, orange), ANH (anhedonic-like, red), and RES (resilient, green). **B.** Boxplots for each behavioral test comparing ANX, ANH, and RES clusters to controls (CTL), in males [(***Left***): 3CH (F_3, 39_=7.88, p<0.001; *Post hoc* CTL *vs* ANX: t=3.79, p=0.002), NSF (F_3, 12.9_=4.33, p=0.025; CTL *vs* ANX: t=–4.22, p<0.001), SPT (F_3, 39_=19.71, p<0.001; CTL *vs* ANH: t=6.98, p<0.001), FST (F_3, 39_=0.84, p=0.48)] and in females [(***Right***): 3CH (F_3, 35_=4.43, p=0.01; CTL *vs* ANX: t=2.71, p=0.03), NSF (F_3, 35_=6.99, p<0.001; CTL *vs* ANX: t=–4.02, p<0.001), SPT (F_3, 35_=31.52, p<0.001; CTL *vs* ANH: t=8.75, p<0.001), FST (F_3, 35_=0.49, p=0.69)]. **C.** Distribution of animals across RES, ANX, and ANH clusters in males (M) and females (F). Data are presented as mean ± SEM. One-way ANOVA (with Welch’s correction when Levene’s test indicated unequal variances) was followed by Bonferroni post hoc tests. Statistical significance is indicated as: *p < 0.05, **p < 0.01, ***p < 0.001. Abbreviations: CTL, control; ELS, early-life stress; ANX, anxiety-like cluster; RES, resilient cluster; ANH, anhedonic-like cluster; FST, Forced swim test; NSF, novelty supressed-feeding test; 3CH, 3-chamber test; SPT, sucrose preference test.

Although behavioral phenotypes were consistent across sexes, cluster distribution differed (**Figure 4C**). The proportion of resilient mice was nearly identical (males: 28%; females: 23.8%). However, males were more frequently assigned to the ANX cluster (48%) than to the ANH (24%), whereas females showed the reverse, with more in ANH (47.6%) than in ANX (28.6%).

In females, the distributions of the 3CH, NSF, and FST stages were similar across clusters (comparison with an expected proportion of 0.5: CTL: 3CH p=0.81, NSF p=0.48, FST p=0.48; RES: 3CH p=1, NSF p=1, FST p=1; ANX: 3CH p=0.69, NSF p=1, FST p=1; ANH: 3CH p=0.34, NSF p=0.1, FST p=0.1). Furthermore, no differences in the proportions of P/E and M/D were observed between clusters, regardless of the behavioral 3CH, NSF, or FST test (3CH: χ²=0.65, p=0.89; NSF: χ²=2.76, p=0.43; FST: χ²=4.78, p=0.21; **Supplementary figure 5C**).

#### Early-life predictors

In males, the ANX cluster displayed reduced body weight from P5 to P14 compared to CTL (**Figure 5A**; F_5.67,73.75_=5.42, p<0.001; *Post hoc*: P5: t=5.88, p<0.001; P7: t=4.90, p<0.001; P12: t=4.63, p<0.001; P14: t=4.28, p<0.001). Consistently, mean early weight was significantly lower in ANX compared to CTL (F_3,15.2_=10.69, p<0.001; t=4.61, p<0.001) and ANH (t=-4.26, p<0.001). In females, both RES and ANH clusters showed persistently reduced body weight from P5 (**Figure 5A**; F_6.3,73.8_=10.7, p<0.001; RES *vs* CTL: P5: t=2.78, p=0.035; P7: t=3.78, p=0.003; P12: t=4.21, p=0.001; P14: t=4.71, p<0.001; ANH *vs* CTL: P5: t=2.52, p=0.049; P7: t=3.12, p=0.011; P12: t=2.80, p=0.034; P14: t=2.73, p=0.04). However, only the RES cluster differed significantly from CTL in mean early weight (F_3.11,165_=10.67, p=0.001; post hoc: t=3.51, p=0.007).

**Figure 5.**
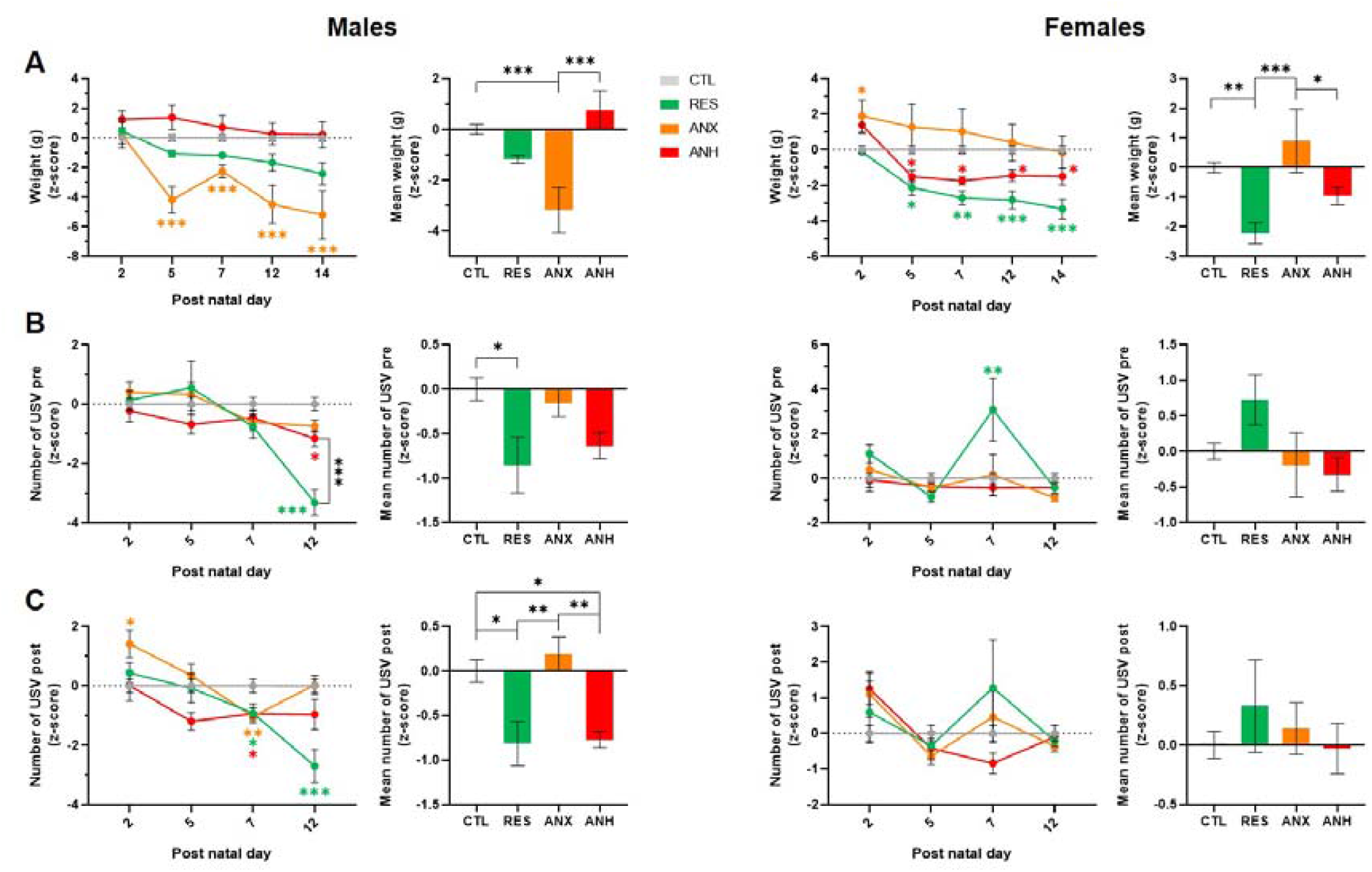
**Early predictors of susceptibility and resilience to the long-term effects of early-life stress identified by k-means clustering analysis. A. Body weight trajectories during ELS (P2–P14)**. Weight development was significantly affected in both males (F_5.7, 73.75_ = 5.42, p < 0.001) and females (F_6.3, 73.8_ = 10.7, p < 0.001). *Left:* In males, the ANX cluster displayed reduced body weight from P5 to P14 compared to CTL (P5: t = 5.88, p < 0.001; P7: t = 4.90, p < 0.001; P12: t = 4.63, p < 0.001; P14: t = 4.28, p < 0.001). Consistently, mean early weight was significantly lower in the ANX cluster compared to both CTL (F_3, 15.2_ = 10.69, p < 0.001; t = 4.61, p < 0.001) and ANH (t = –4.26, p < 0.001). *Right:* In females, both RES and ANH clusters showed reduced body weight from P5 to P14 compared to CTL (RES vs. CTL: P5: t = 2.78, p = 0.035; P7: t = 3.78, p = 0.003; P12: t = 4.21, p = 0.001; P14: t = 4.71, p < 0.001; ANH vs. CTL: P5: t = 2.52, p = 0.049; P7: t = 3.12, p = 0.011; P12: t = 2.80, p = 0.034; P14: t = 2.73, p < 0.04). For mean early weight, only the RES cluster differed significantly from CTL (F_3.11, 165_ = 10.67, p = 0.001; t = 3.51, p = 0.007). **B. Pre-deprivation USV emissions during ELS (P2–P12)**. USVs prior to daily maternal deprivation were significantly affected by ELS in males (F_6.8, 88.4_ = 5.22, p < 0.001) and females (F_7.8, 91.6_ = 3.8, p < 0.001). *Left:* In males, both ANH and RES clusters showed significantly fewer USVs at P12 compared to CTL (ANH: t = 2.79, p = 0.024; RES: t = 8.48, p < 0.001), with a stronger reduction in the RES cluster (RES vs. ANH: t = –4.42, p < 0.001). Mean pre-deprivation USVs were significantly reduced in the RES cluster compared to CTL (F_3, 15.6_ = 4.44, p = 0.019; t = 3.31, p = 0.012). *Right:* In females, the RES cluster showed higher USV emissions at P7 compared to CTL (t = –3.76, p = 0.003). **C. Post-deprivation USV emissions during ELS (P2–P12)**. *Left:* In males, while USV emissions were reduced in all groups compared to CTL at P5 (F_7.8, 101.6_ = 5.23, p < 0.001; ANX: t=3.7, p=0.004; RES: t=2.75, p=0.045; ANH: t=2.64, p=0.047), only the RES cluster remained significantly reduced compared to CTL at P12 (t=5.40, p<0.001). Moreover, ANX displayed elevated post-deprivation USVs at P2 compared to CTL (t=–3.18, p=0.017). Mean post-deprivation USVs were reduced in both RES and ANH compared to CTL (F_3,17.3_=7.5, p<0.001; RES: t=3.25, p=0.015; ANH: t=2.90, p=0.036). *Right:* In females, no significant cluster differences were detected according to post-natal day (F_6.5, 76.2_= 2.45, p = 0.028) or the mean number of USV emission (F_3,39_=0.51, p=0.68). Data are presented as mean ± SEM. Two-way ANOVA followed by Bonferroni’s multiple comparisons test was used, with Greenhouse–Geisser correction applied when sphericity was violated. Statistical significance is indicated as: *p < 0.05, **p < 0.01, ***p < 0.001. Colored asterisks indicate significant differences from the CTL group. Abbreviations: CTL, control; ELS, early-life stress; ANX, anxiety-like cluster; RES, resilient cluster; ANH, anhedonic-like cluster.

Cluster-specific USV patterns also emerged (**Figure 5B-C**). In males, RES and ANH showed reduced pre-deprivation USVs at P12 (F_6.8,88.4_=5.22, p<0.001; ANH: t=2.79, p=0.024; RES: t=8.48, p<0.001) with a stronger effect in RES (RES vs ANH: t=–4.42, p<0.001). Post-deprivation, this P12 reduction persisted only in RES (F_7.82,101.64_ =5.23, p<0.001; t=5.40, p<0.001) while at P7, reduced USV is observed in all groups compared to CTL (ANX: t=3.7, p=0.004; RES: t=2.75, p=0.045; ANH: t= 2.64, p=0.047). Consistently, mean pre-deprivation USV level was reduced only in RES (F_3,15.6_=4.44, p=0.019; t=3.31, p=0.012), while mean post-deprivation USVs were reduced in both RES and ANH compared to CTL (F_3,17.3_=12.15, p<0.001; RES: t=3.24, p=0.015; ANH: t=2.90, p=0.036). Conversely, ANX displayed elevated post-deprivation USVs at P2 compared to CTL (F_7.8,101.6_=5.23, p<0.001; *Post hoc*: t=–3.18, p=0.017). In females, the only effect was a transient pre-deprivation increase in USVs at P7 in RES (Pre: F_7.8,91.6_=3.8, p<0.001; P7, CTL vs RES: t=–3.76, p=0.003; Post: F_6.5, 76.2_=2.45, p=0.028, post hoc *ns*).

#### Late-onset markers

In males, despite lower baseline corticosterone in RES (F_3,39_=8.407, p<0.001; CTL *vs* RES: t=4.98, p<0.001; ANX vs RES: t=3.36, p=0.01; **Figure 6A**), a greater stress-induced increase was observed (F_3,15_=4.32, p=0.022; CTL *vs* RES: t=–4.62, p<0.001; RES *vs* ANX: t=–3.57, p=0.006; **Figure 6B**). No cluster differences compared to CTL emerged post-FST (F_3,15.6_=1.82, p=0.19), 1h30 post-FST (F_3,39_=3.71, p=0.019; **Supplementary Figure 7A-B**) or for percentage reduction (F_3.39_=0.78, p=0.51; **Figure 6C**). In females, corticosterone levels or percentage increase post-FST did not differ across clusters at any time point (Baseline: F_3,9.1_=3.44, p=0.13; Post-FST: F_3,35_=0.55, p=0.65; 1h30 post-FST: F_3,35_=2.06, p=0.12; %Increase: F_3,9.2_=1.48, p=0.28; **Figure 6A-B, Supplementary Figure 7A-B**). However, the percentage of decrease 1h30 post-FST was significantly increased in the ANX cluster compared to CTL (%Decrease: F_3,35_=3.4, p=0.028; t=-2.95, p=0.034; **Figure 6C**).

**Figure 6.**
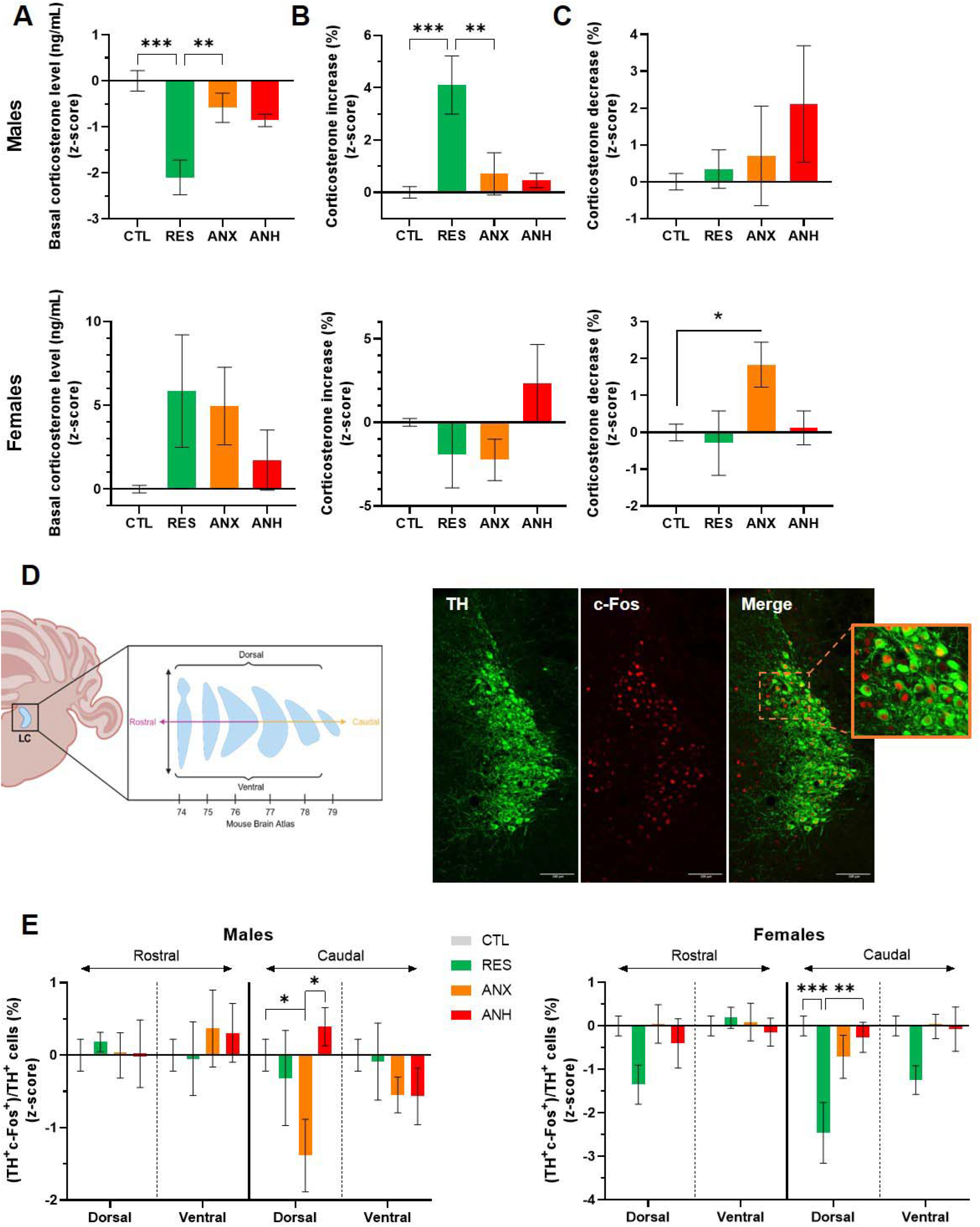
Physiological predictors and functional anatomy of LC-NE activity in susceptibility and resilience to the long-term effects of early-life stress identified by k-means clustering analysis. **A. Baseline corticosterone levels.** In males (*Top*), baseline corticosterone levels were significantly lower in the RES cluster compared to CTL and ANX (F_3, 39_ = 8.41, p < 0.001; CTL *vs* RES: t = 4.98, p < 0.001; ANX *vs* RES: t = 3.36, p = 0.01). In females *(Bottom*), no significant cluster effects were observed (F_3, 9.1_ = 3.44, p = 0.13). **B. Percentage increase in corticosterone following FST**. In males (*Top*), the percentage increase in corticosterone after FST was greater in the RES cluster compared to CTL and ANX (F_3, 15_ = 4.32, p = 0.022; CTL *vs* RES: t = –4.62, p < 0.001; CTL *vs* ANX: t = –3.57, p = 0.006). In females (*Bottom*), no significant cluster effect was detected (F_3, 9.2_ = 1.48, p = 0.28). **C. Percentage decrease in corticosterone levels 1h30 after FST.** No significant cluster effects were observed in males (F_3, 39_= 0.78, p = 0.51). In females, the percentage decrease in corticosterone 1h30 post-FST was greater in the ANX cluster compared to CTL (F_3, 35_ = 3.4, p = 0.0.028, t=-2.95, p=0.034). **D. Schematic representation of the locus coeruleus** (LC, *left*) and representative immunofluorescence image (Created in BioRender. Sla, D. (2025) https://BioRender.com/7ipzo68) (*Right*) showing TH-positive neurons (green), c-Fos expression (red), and colocalization of both markers (yellow), indicating activated noradrenergic (NE) cells. **E. Proportion of activated NE neurons (%TH^+^C-Fos^+^/TH^+^) along the rostro-caudal and dorso-ventral axes of the LC**. In males (*Left*), the proportion of activated NE neurons was significantly reduced in the ANX cluster compared to CTL and ANH, specifically in the caudal-dorsal LC (F_9, 117_ = 2.15, p = 0.031; CTL *vs* ANX: t = 2.81, p = 0.046; ANH *vs* ANX: t = –2.69, p = 0.05). In females (*Right*), a similar reduction was observed in the RES cluster relative to CTL and ANH (F_9, 105_ = 2.52, p = 0.012; CTL *vs* RES: t = 3.34, p < 0.001; ANH *vs* RES: t = –3.58, p = 0.005). Data are presented as mean ± SEM. One-way ANOVA (with Welch’s correction when Levene’s test indicated unequal variances) was followed by Bonferroni’s post hoc test. Statistical significance is indicated as *p < 0.05, **p < 0.01, ***p < 0.001. Abbreviations: CTL, control; ELS, early-life stress; ANX, anxiety-like cluster; RES, resilient cluster; ANH, anhedonic-like cluster; FST, forced swim test; LC, locus coeruleus; TH, tyrosine hydroxylase; NE, noradrenergic.

LC-NE analyses revealed sex-specific patterns (**Figure 6D-E**). In males, ANX showed reduced proportion of activated NE cells (%(TH^+^c-Fos^+^)/TH^+^) selectively in the caudal-dorsal LC (F_9,117_=2.15, p=0.031; CTL *vs* ANX: t=2.81, p=0.046; ANH *vs* ANX: t=-2.69, p=0.05), while in females, this reduction was observed in RES (F_9,105_=2.52, p=0.012; CTL *vs* RES: t=3.34, p<0.001; ANH *vs* RES: t=-3.58, p=0.005). No significant differences emerged across phenotypes for the density of activated NE or non-NE cells along LC sub-regions (**Supplementary figure 5C-D**).

## Discussion

This study investigated how ELS shapes anxio-depressive-like behaviors, with emphasis on resilience in the context of CA. Since CA is a major psychiatric risk factor, uncovering resilience mechanisms is critical. The LC-NE system has been implicated in resilience in several adult stress model^47,49,51,64–66^, however, it has not been studied in relation to resilience following ELS. Using a paradigm combining MD with limited bedding/nesting, we defined resilient versus susceptible phenotypes and identified early- and long-term markers. Findings revealed sex-specific behavioral trajectories associated with distinct physiological and neurobiological signatures (**Figure 7**). In males, reduced body weight with increased first post-deprivation USV was linked to susceptibility, especially anxiety-like phenotype, whereas prolonged USV reductions until P12 characterized resilience. In females, reduced body weight emerged in both resilient and depressive-like phenotypes but was greater in resilience. Increased USV emission was associated with resilience. Later trajectories also diverged by sex. Male resilience was associated with lower baseline corticosterone and enhanced corticosterone response to acute stress, while anxiety-like was associated with reduced LC-NE activity in the caudal-dorsal LC. Conversely, in females, this reduction marked resilience.

**Figure 7.**
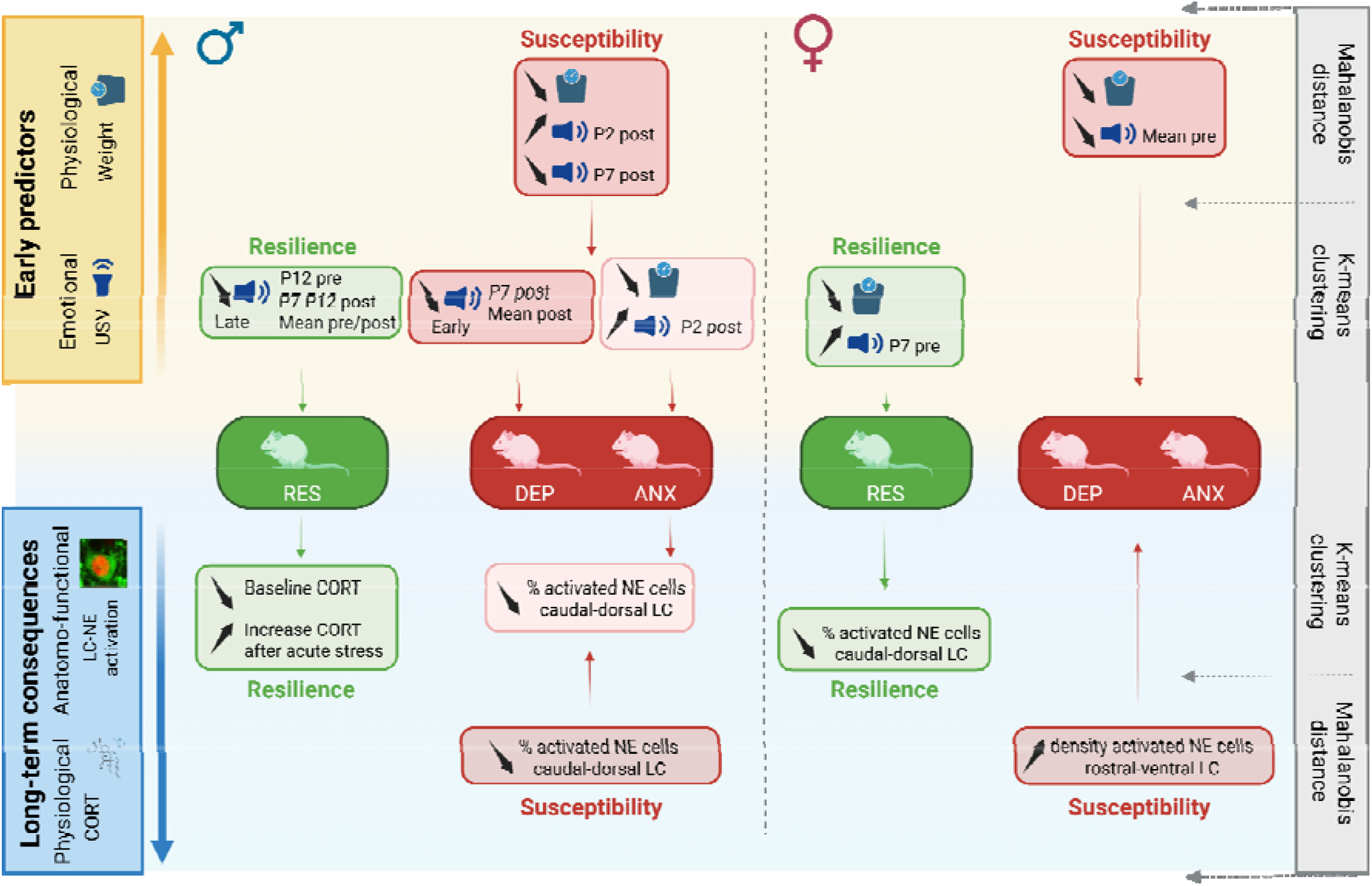
Summary of main findings in each phenotype in both males and females. Created in BioRender. Sla, D. (2025) https://BioRender.com/rv4ejoi.

In the presented MD model, dams’ behavior and stress response were evaluated as factors shaping pup development. This variant was designed to replicate CA by introducing maternal stress, counteracting the increased care often observed in deprivation alone. Here, maternal behavior and long-lasting stress response remained unaffected^60,67^. Additional real-time analyses could have refined these findings^68^. However, ELS reduced pup survival, especially in large litters, an effect rarely reported in deprivation paradigms^61,69–72^. Litter equalization was avoided to reduce disturbance^73^. Body weight was unaffected in females but reduced in ELS males from P7 onward, persisting into adulthood (**Figure 1**).Consistent with this sex-specific effect, maternal deprivation has been reported to reduce food intake more markedly in males than in females and to selectively increase hypothalamic leptin receptor expression and amygdalar serotonin levels in males ^74^. Together with evidence that leptin-mediated inhibition of serotonin is required for the postnatal suppression of appetite ^75^, these findings suggest that altered leptin–serotonin interactions may contribute to the male-specific reduction in body weight following ELS. ELS also reduced USVs, particularly in males. We did not replicate the pre/post-separation increase observed in females at P7 in the study of Yin et al. (2016) ^76^, possibly due to methodological differences^59^. Together, these findings highlight sex-specific early effects of ELS, with males showing greater vulnerability.

In the long-term, opposing sex-specific outcomes emerged (**Figure 2**). In males, ELS had no impact on anxio-depressive behaviors, while in females it increased anhedonia. These findings align with the resilience of C57BL/6 mice in ELS paradigms and variability across models^77,78^. For instance, MD associated with limited nesting showed no effect on anhedonia in females^79^ but produces social deficits in males^80^. To standardize comparisons, we assessed both sexes across tests covering anxiety-and depression-like dimensions^81,82^. To identify susceptible from resilient individuals, we employed Mahalanobis distance and k-means clustering analysis. Mahalanobis distance, typically used to detect outliers^83^, is a multivariate deviation metric which identified behaviors deviating most from controls, defined as susceptibility, while clustering^84^, classified mice into three clusters, including anxious-like (ANX), anhedonic-like (ANH), and resilient-like (RES) phenotypes^85^. This framework captured stress-induced phenotypes beyond task-specific effects. In males, susceptibility involved anxiety and anhedonia, while in females it was marked only by anhedonia. Clustering confirmed this, showing higher prevalence of anhedonic phenotypes in females and anxiety-like phenotypes in males. These results parallel clinical evidence, where women are twice as likely to develop depression^86,87^. Resilience prevalence was similar in both sexes (∼25%), consistent with adult chronic stress models^49,88–90^. In humans, resilience rates following CA varies widely (10–50%) depending on definitions, age, sex, and type of abuse^91–93^.

Resilience is increasingly recognized as a dynamic and adaptive process fluctuating across life rather than a fixed trait^18,94^. Human studies rarely clarify whether biobehavioral factors act as predictors or consequences of resilience. Animal models help address this gap by revealing phenotypic changes across multiple levels. In this study, we examined body weight and USVs during the stress period as potential early predictors (**Figures 5**). In males, both lower body weight and reduced USV emission linked to susceptibility in the Mahalanobis distance and confirmed by clustering, which tied them to an anxiety-like phenotype. While USV reductions appeared across clusters, they persisted only in resilient males until P12. Notably, anxiety-linked USV decreases followed an initial rise after the first deprivation, a pattern absent in resilient mice, suggesting early adaptive dampening of emotional responses. In females, weight loss occurred in both resilient and depressive-like clusters, but its association with increased USVs predicted resilience, whereas decreased USVs predicted susceptibility. These findings highlight sex-specific physiological and emotional markers as early predictors of long-term ELS outcomes.

ELS models often fail to capture human-like long-term effects, partly because resilience is overlooked. Our analysis highlighted that excluding the ∼25% resilient individuals revealed anxio-depressive outcomes. Beyond this, we identified physiological and anatomical signatures linked to both susceptible and resilient phenotypes (**Figure 6**). Corticosterone levels were not predictive in the Mahalanobis analysis, but clustering revealed phenotype-specific regulation, suggesting efficient adaptation in resilient males. Both patterns have been observed in humans, underscoring the complexity of resilience markers and making it challenging to establish clear conclusions^95^. Associations between behavioral phenotypes induced by ELS and LC-NE functional anatomy differed by sex. In males, Mahalanobis analysis linked susceptibility to reduced NE neuron activation in the caudal-dorsal LC, associated with anxiety-like traits in clustering. No adaptations were observed in resilient males. In female, resilient mice displayed reduced caudal-dorsal NE activation in the clustering analysis.

These sex differences likely arise from both developmental and signalling mechanisms. For instance, adult females have a larger LC than males due to a greater number of NE-containing neurons, possibly resulting from prolonged post-pubertal neurogenesis and hormone-driven developmental effects. Functionally, females show stronger LC responses to CRF than males. However, after stress, male LC neurons shift toward a fully cAMP-mediated pathway, a female-like profile, whereas female signalling remains stable^96^. Additionally, GABAergic input from peri-LC and dorso-medial regions known to inhibit NE activity^30,97,98^, may contribute to the sex-dependent LC-NE activation patterns observed in the present study, by buffering NE activity in resilient males while promoting adaptive reductions in resilient females. Another potential mechanism involves α2-adrenergic autoreceptor-mediated inhibition of LC neurons. Because these autoreceptors provide negative feedback on noradrenergic firing, changes in α2-autoreceptor function could contribute to the sustained LC hyperactivity induced by early-life stress. Consistent with this idea, maternal separation has been shown to decrease α2-autoreceptor expression in the locus coeruleus ^99^, while increasing LC firing rates ^46^. In our study, reduced LC-NE activation in resilient males may therefore reflect a compensatory restoration of inhibitory NE control. By contrast, the similar reduction observed in anxious females suggests that LC inhibition may have distinct functional consequences depending on sex, in line with the marked sexual dimorphism of the LC-NE system. Whether early-life stress differentially alters α2-autoreceptor regulation in males and females remains an important question for future studies. Importantly, the present study does not establish a causal contribution of LC-NE activity to resilience or susceptibility. LC-NE activation was measured after behavioral phenotyping and following an acute stress challenge, and therefore it should be interpreted as a neurobiological correlate of the identified phenotypes. Future studies using longitudinal analysis with circuit-specific recordings and manipulations will be required to determine whether these LC-NE adaptations actively contribute to resilient or susceptible outcomes. Identification of the developmental window during which early-life stress disrupts the LC-NE system could open new directions on how this system shapes brain maturation and drives the region-specific and behavioural alterations observed following early adversity.

This study highlights sex-specific mechanisms underlying susceptibility and resilience to ELS, identifying early emotional and physiological correlates and distinct LC-NE system adaptations. In males, reduced caudal-dorsal LC-NE activation was associated to anxiety-like behaviors, whereas in females a similar activation pattern was observed in resilient animals. Characterizing resilient and susceptible trajectories following ELS may improve our understanding of individual variability in stress outcomes and help identify biological processes associated with adaptive responses to adversity.

## Supporting information

Supplementary tables 1 to 6 - statistical analyses for Figures 1-6

Supplementary tables 7 to 13 - statistical analyses for supplementary Figures 1-6

## Acknowledgements

The authors express their gratitude to the financial support of the ANR – FRANCE (French National Research Agency, ANR-21-CE16-004), the FRM (Fondation pour la recherche médicale, EQU202203014667), the Sissley Fundation, the IDEX Emergence grant and the doctoral school MCTI (ED 563, Médicament, Toxicologie, Chimie, Imageries) from the University Paris Cité. The project was carried out with the support of the ERIE Foundation, a fund hosted by the King Baudouin Foundation. Image acquisitions were performed at the SCM Imaging facility and Corticosterone dosage at the Cyto2BM facility (BioMedTech Facilities, INSERM US36 | CNRS UAR2009 | Université Paris Cité, https://biomedicale.u-paris.fr/biomedtech-facilities/).

## Author contributions

EI and DS designed the experiments. DS and VG performed the behavioral and physiological experiments and analysis. DS and VG performed the anatomical experiments and VG the counting analysis. DS, VG and EI performed the statistical analysis. LG performed the matlab script for the mahalonobis distance analysis. DS and EI wrote the publication, which was edited by BG. EI supervised this research.

