## Supplementary Figures 1 to 7 for "Behavioral susceptibility and resilience following early-life stress are associated with sex-dependent locus coeruleus noradrenergic signatures in adulthood"

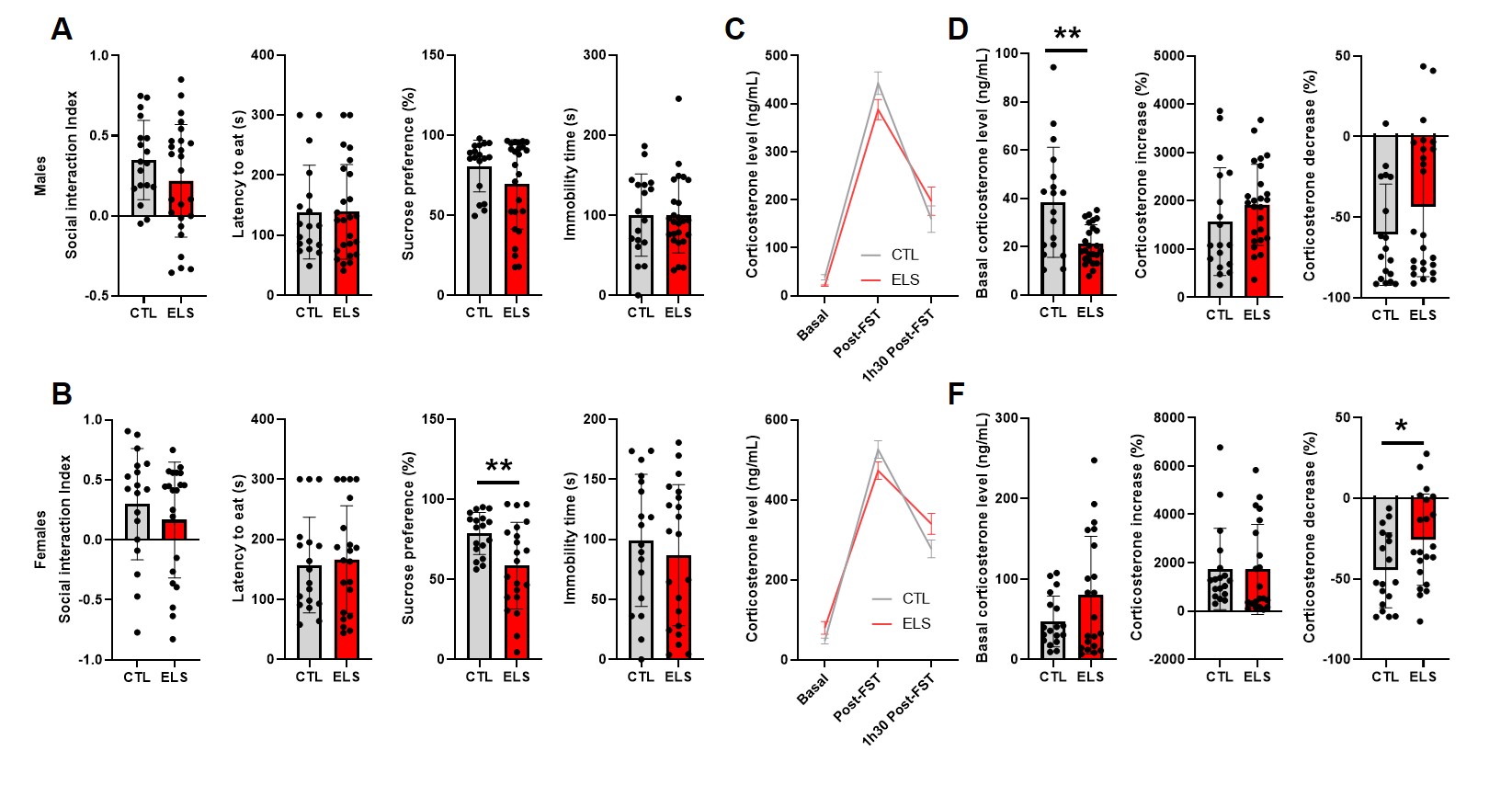
**Supplementary figure 1:** **ELS consequences on behavior and corticosterone level in males and females (raw data). (A)** In males, no significant differences between CTL and ELS groups were revealed for the social interaction index in the 3CH (t_40.91_=-1.42, p=0.162), the latency to eat in the NSF (U_41_=220.5, p=0.92), the percentage of sucrose preference (SPT, U_41_=198, p=0.52) and the immobility time in the FST (t_41_=-0.01, p=0.99). **B.** In females, no significant differences between CTL and ELS groups were revealed for the social interaction index in the 3CH (U_37_=160, p=0.44), the latency to eat in the NSF (U_37_=194, p=0.9) and for the time immobile in the FST (t_37_=-0.67, p=0.51). However, the percentage of sucrose preference was significantly reduced in ELS group compared to CTL (W_28.81_=-2.9, p=0.005).**C.** The evolution of corticosterone levels from baseline to post-FST to 1h30 post-FST did not differ between CTL and ELS groups in both male (F_1.58,64.66_=2.366, p=0.1; Up) and female (F_2,74_=4.55, p=0.014; *posthoc ns*; Bottom). **E.** In males, ELS induced a significant decrease in baseline corticosterone level compared to CTL (t_20.03_=-3.09, p=0.006), but had no effect on the percentage of increase from basal to immediately after the FST (t_41_=1.17, p=0.25) or the percentage of decrease from immediately post-FST to 1h30 later (% decrease: U_41_=287, p=0.13). **F.** In females, ELS had no significant effect in baseline corticosterone level compared to CTL (U37=210, p=0.57) and on the percentage of increase from basal to immediately after the FST (U_37_=155, p=0.35). However, ELS induced a significant decrease in the percentage of decrease from immediately post-FST to 1h30 later (% decrease: T_37_=2.27, p=0.0.029). Data are presented as mean ± SEM. Student’s t-test was used, with Welch’s correction applied when homogeneity of variance was violated. Mann–Whitney U test was used when normality assumptions were not met. Two-way ANOVA followed by Bonferroni's multiple-comparisons test was used, with Greenhouse–Geisser correction applied when the assumption of sphericity was violated. Statistical significance is indicated as *p < 0.05, **p < 0.01, and ***p < 0.001. Abbreviations: CTL, control; ELS, early-life stress; FST, forced swim test.

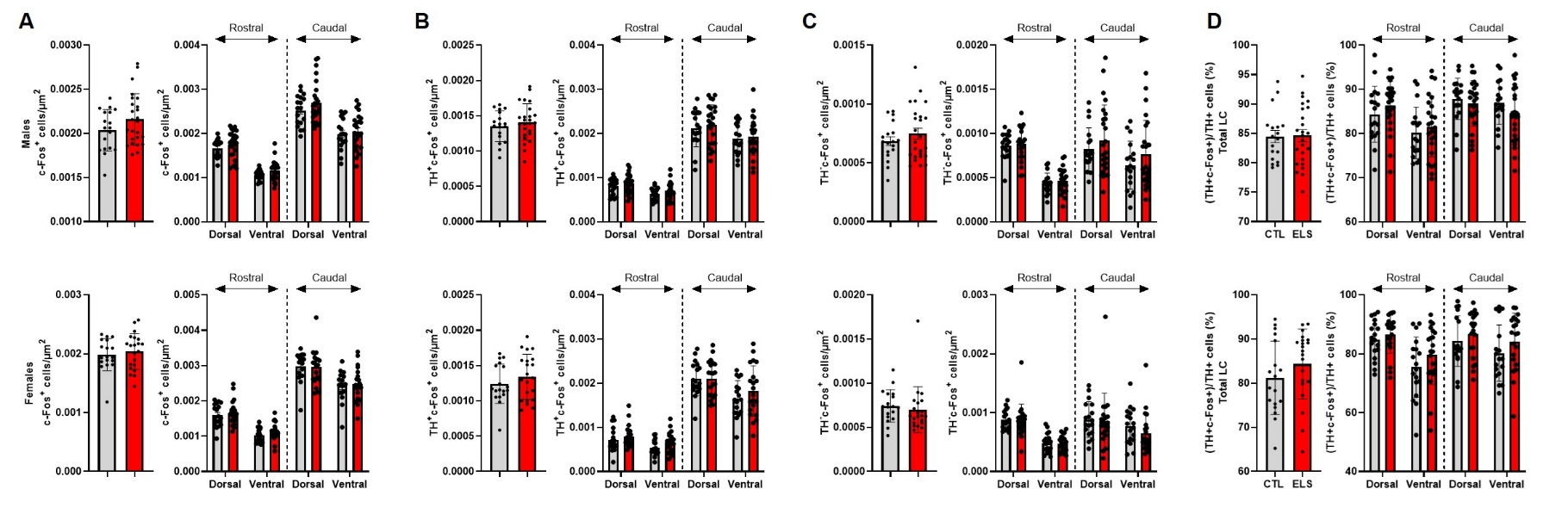

**Supplementary figure 2: ELS consequences on LC-NE functional anatomy in males and females (raw data).** No significant effect of ELS was detected on the total LC (left panels) or across the rostro-caudal and dorso-ventral axes of the LC (right panels) in either males (top panels) or females (bottom panels) for: (**A**) the density of activated neurons (c-Fos⁺/µm²; M: t_41_=1.47, p=0.15; F_1.72,70.56_=0.29, p=0.72; F: U_37_=204, p=0.69; F_2.24,82.85_=0.44, p=0.67), (**B**) the density of activated noradrenergic neurons (TH⁺c-Fos⁺/µm²; M: t_41_=0.78, p=0.44; F_1.83,75.17_=0.050, p=0.94; F: t_37_=1.015, p=0.32; F_1.89,70.02_=0.89, p=0.41), (**C**) the density of activated non-noradrenergic neurons (TH⁻c-Fos⁺/µm² M: U_41_=243, p=0.93; F_1.68,68.92_=0.77, p=0.45; F: U_37_=142, p=0.19; F_2.02,74.61_=0.64, p=0.53), and (**D**) the percentage of activated noradrenergic neurons (TH⁺c-Fos⁺/TH⁺; M: t_41_=0.092, p=0.92; F_3,123_=2.56, p=0.058; F: U_37_=230, p=0.26; F_2.71,100.19_=0.81, p=0.48). Data are presented as mean ± SEM. Student’s t-test was used, with Welch’s correction applied when homogeneity of variance was violated. Mann–Whitney U test was used when normality assumptions were not met. Two-way ANOVA followed by Bonferroni's multiple-comparisons test was used, with Greenhouse–Geisser correction applied when the assumption of sphericity was violated. Statistical significance is indicated as *p < 0.05, **p < 0.01, and ***p < 0.001. Abbreviations: LC, locus coeruleus; TH, tyrosine hydroxylase; NE, noradrenergic; CTL, control; ELS, early-life stress.

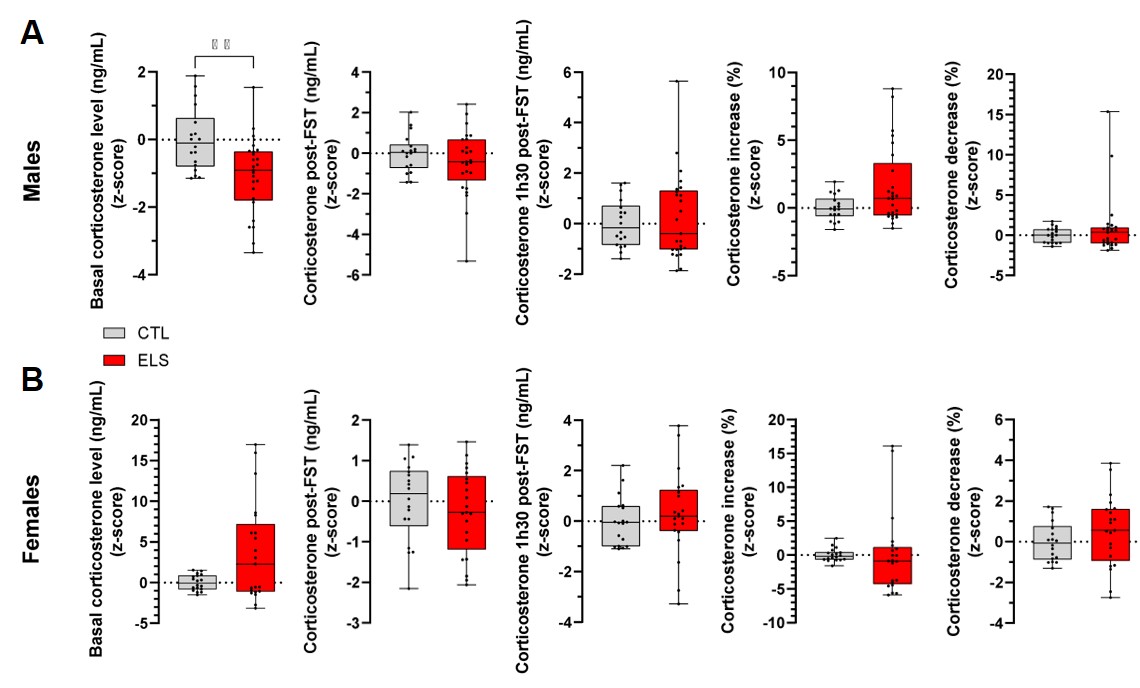

**Supplementary figure 3: ELS consequences on corticosterone level in males and females** **(Z-score).** (**A)** In males, ELS induced a significant decrease in baseline corticosterone level compared to CTL (t_41_=3.3, p=0.002), but had no effect immediately after the FST (post-FST: t_41_=1.06, p=0.30; % increase: U_41_=164, p=0.14) or 1h30 later (1h30 post-FST: U_41_=231, p=0.89; % decrease: U_41_=218, p=0.87). (**B)** In females, corticosterone levels did not differ between ELS and CTL animals at baseline (U_37_=150, p=0.28), post-FST (t_37_=0.8, p=0.43), or 1h30 post-FST (t_37_=–0.66, p=0.52). Similarly, the relative increase (U_37_=238, p=0.17) and subsequent decrease (t_32.13_=–1.16, p=0.25) were unaffected by ELS. Data are presented as mean ± SEM. Student’s t-test was used, with Welch’s correction applied when homogeneity of variance was violated. Mann–Whitney U test was used when normality assumptions were not met. Statistical significance: *p < 0.05, **p < 0.01, ***p < 0.001. Abbreviations: CTL, control; ELS, early-life stress; FST, forced swim test.

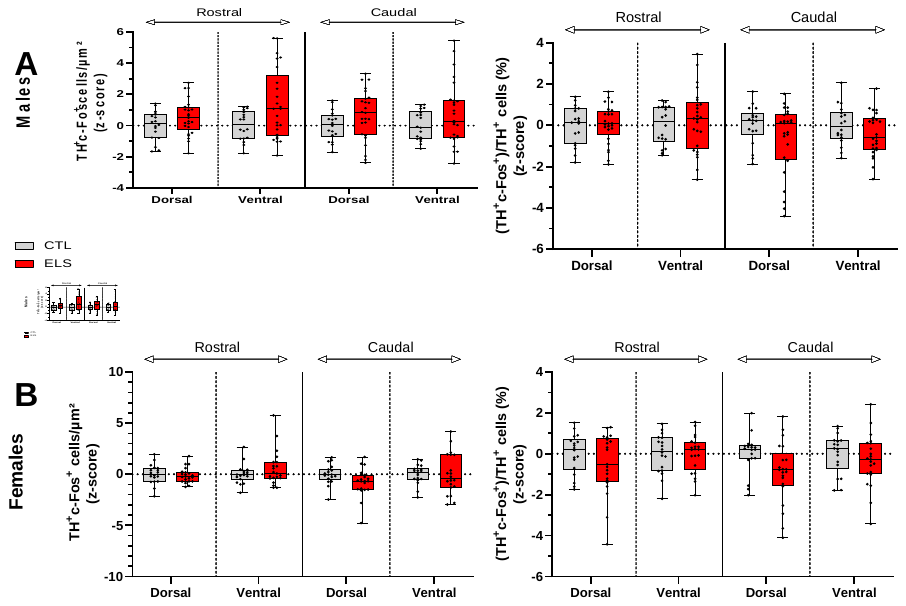

**Supplementary Figure 4. Long-term consequences of ELS on LC-NE functional anatomy in males and females (Z-score).** Along the rostro-caudal and dorso-ventral axis of the LC, neither the density of activated LC-NE neurons (TH^+^c-Fos^+^/µm²; M: F_2.5,103.1_=0.92, p=0.42; F: F_3,111_=1.83, p=0.15) nor the proportion of activated LC-NE neurons (TH^+^c-Fos^+^/TH^+^; M: F_3,123_=1.86, p=0.14; F: F_2.4,90.7_=2.49, p=0.076) were significantly altered by ELS compared to CTL, in either males (**A**) or females (**B**). Data are presented as mean ± SEM. Two-way ANOVA followed by Bonferroni’s multiple comparisons test was performed, with Greenhouse–Geisser correction applied when sphericity was violated. Statistical significance: *p < 0.05, **p < 0.01, ***p < 0.001. Abbreviations: LC, locus coeruleus; TH, tyrosine hydroxylase; NE, noradrenergic; CTL, control; ELS, early-life stress.

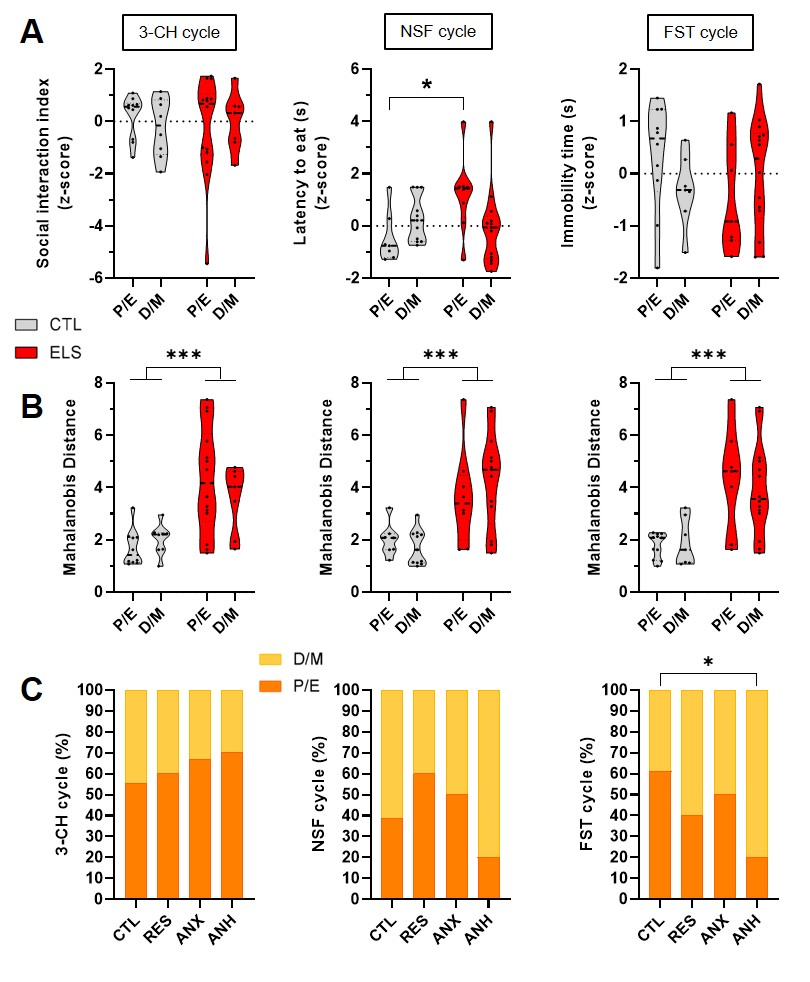

**Supplementary Figure 5. Impact of the oestrus cycle on behavior in females.** (**A**) The oestrus cycle stage (proestrus/estrus, P/E vs diestrus/metestrus, D/M), determined after the 3CH and FST, had no effect on the social interaction index or immobility time in either CTL or ELS groups (3CH: F_1,35_=0.51, p=0.48; FST: F_1,35_=2.05, p=0.16). However, when the cycle stage was determined after the NSF, females in the P/E stage displayed longer latency to eat in the ELS group compared to CTL (F_1,35_=5.72, p=0.022; P/E: t_35_=-2.51, p=0.017), whereas no difference was observed in the D/M stage (t_35_=0.69, p=0.5). (**B**) The Mahalanobis distance was significantly higher in ELS compared to CTL animals, independently of cycle stage (P/E or D/M) (CTL vs ELS; 3CH: F_1.35_=19.53, p<0.001; NSF: F_1,35_=20.1, p<0.001; FST: F_1,35_=22.06, p<0.001. (**C**) K-means clustering analysis revealed that the distribution of P/E and D/M stages was similar across all clusters when determined after the 3CH, NSF and FST (proportion tested against 0.5, P/E vs D/M). However, when comparing CTL and ANH clusters based on cycle stage determined after the FST, a significant difference was observed: 60% of CTL females were in P/E, whereas only 20% of ANH females were in P/E (X^2^=4.37, p=0.037). No such differences were detected when comparing CTL with RES or ANX clusters. Data are presented as mean ± SEM. Contingency tables with chi-square tests were used for comparisons with the CTL group, and binomial tests were applied for P/E vs D/M distributions. Statistical significance: *p < 0.05, **p < 0.01, ***p < 0.001. Abbreviations: CTL, control; ELS, early-life stress; P/E, proestrus/estrus; D/M, diestrus/metestrus, ANX, anxiety-like cluster; RES, resilient cluster; ANH, anhedonic-like cluster; 3CH, 3-chambers test; NSF, novelty-supressed feeding test; FST, forced swim test.

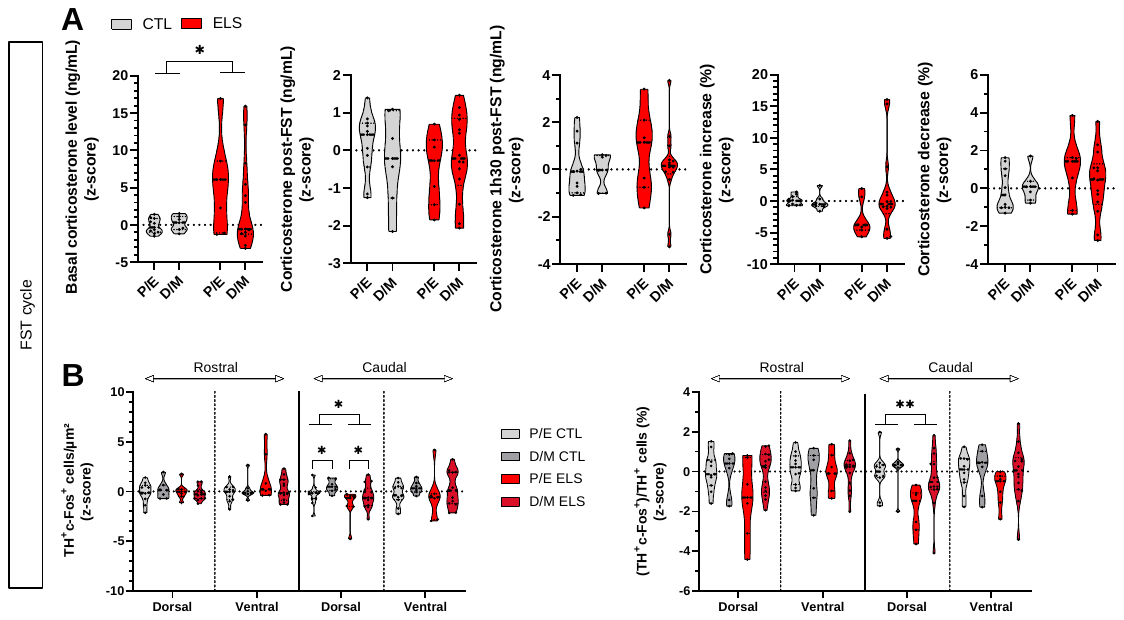

**Supplementary figure 4: Impact of the oestrus cycle in female on corticosterone levels and LC-NE functional anatomy.** (**A**) The oestrus cycle stage (proestrus/estrus, P/E vs diestrus/metestrus, D/M), determined after the FST, had no effect on corticosterone levels at baseline (F_1,35_=0.94, p=0.34), immediately post-FST (F_1,35_=1.09, p=0.30), or 1h30 post-FST (F_1,35_=0.40, p=0.53) in either CTL or ELS groups. However, baseline corticosterone levels were significantly higher in ELS compared to CTL animals, independently of cycle stage (F_1,35_=6.97, p=0.012). The relative corticosterone increase post-FST (F_1,35_=0.30, p=0.14) and decrease at 1h30 post-FST (F_1,35_=0.65, p=0.42) were unaffected by cycle stage in both groups. (**B**) The density of activated LC-NE neurons (TH+c-Fos+/µm², left) did not differ across cycle stages or conditions in the rostral LC, but was significantly affected in the caudal LC by both condition (structure × condition: F_2.46,86.14_=4.22, p=0.012) and cycle stage (structure × cycle: F_2.46,86.14_=3.12, p=0.039). Post-hoc tests revealed an increased density of activated neurons in D/M compared to P/E stages in the caudal-dorsal LC (t_35_=–2.313, p=0.027), independent of condition. In addition, ELS animals showed a reduced density compared to CTL, regardless of cycle stage (t_35_=2.63, p=0.013). The proportion of activated LC-NE neurons (% TH+c-Fos+/TH+, right) was unchanged across cycle stages or conditions in the rostral LC, but was significantly reduced in the caudal dorsal LC of ELS animals compared to CTL (structure × condition: F_3,105_=3.12, p=0.029; post-hoc: t_35_=2.99, p=0.005), independently of cycle stage. Data are presented as mean ± SEM. Three-way ANOVA followed by Bonferroni’s multiple comparisons test was used. Statistical significance: *p<0.05, **p<0.01, ***p<0.001. Abbreviations: LC, locus coeruleus; TH, tyrosine hydroxylase; NE, noradrenergic; CTL, control; ELS, early-life stress; P/E, proestrus/estrus; D/M, diestrus/metestrus.

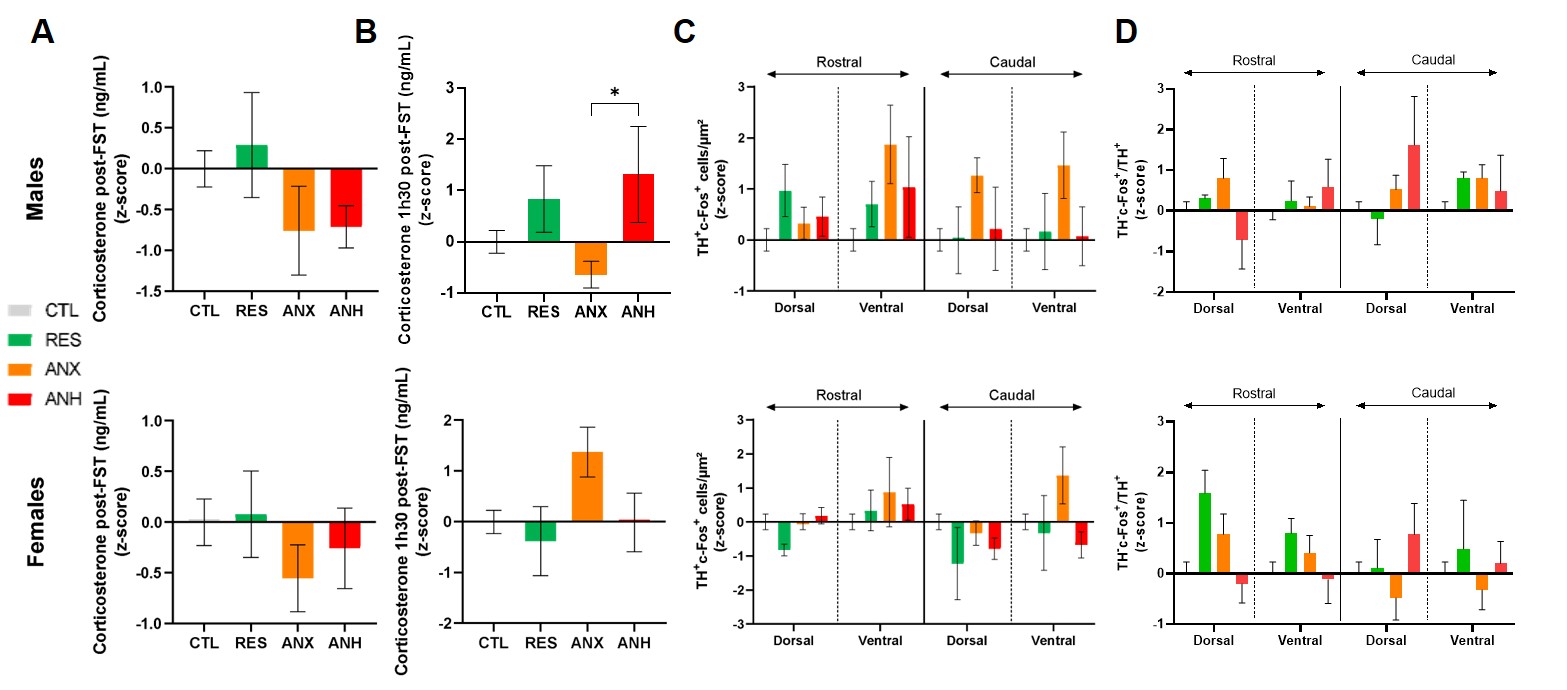

**Supplementary figure 5: Long-term consequences of ELS on corticosterone level and LC-NE functional anatomy in males and females.** (**A**) Corticosterone level post-FST was similar between clusters in male (*Top*, F_3,15.6_=1,82, p=0.19) and female (*Bottom*, F_3,35_=0.55, p=0.65). (**B**) Corticosterone level 1h30 post-FST, was similar between clusters in females (*Bottom*, F_3,35_=2.06, p=0.12), however, in males, an increase was observed in the ANH compared to the ANX cluster (*Top*, F_3,39_=3.71, p=0.019; t_39_=-2.97, p=0.025). (**C**)The density of activated LC-NE neurons (TH^+^c-Fos^+^/µm^2^) was similar between cluster across the rostro-caudal and dorso-ventral axis of the LC both in males (*Top*, F_7.2,97.3_=1.37, p=0.22) and females (*Bottom*, F_7.5,87.6_=1.96, p=0.065). (**D**) The density of activated non-NE neurons in the LC (TH^-^c-Fos^+^/µm^2^) was similar between cluster across the rostro-caudal and dorso-ventral axis of the LC both in males (*Top*, F_6.057, 78.74_=2.3, p=0.042) and females (*Bottom*, F_7.29,85_=1.72, p=0.11). Data are presented as mean ± SEM. Two-way ANOVA followed by Bonferroni’s multiple comparisons test was used, with Greenhouse–Geisser correction applied when sphericity was violated. Statistical significance: *p<0.05, **p<0.01, ***p<0.001. Abbreviations: LC, locus coeruleus; TH, tyrosine hydroxylase; NE, noradrenergic; CTL, control; ELS, early-life stress; ANX, anxiety-like cluster; RES, resilient cluster; ANH, anhedonic-like cluster P/E, proestrus/estrus; D/M, diestrus/metestrus.
