## Supplementary tables 1 to 6 - statistical analyses for Figures 1-6 for "Behavioral susceptibility and resilience following early-life stress are associated with sex-dependent locus coeruleus noradrenergic signatures in adulthood"

**Supplementary Table 1 | Detailed Statistics for Figure 1**

| Figure | N | | Test | Factor | Statistic | Statistic Value | *p* value | Notes |
|---|---|---|---|---|---|---|---|---|
| **1B.Maternal behaviour - Time on nest** | **CTL = 9, ELS = 11** | | Test of Sphericity | Day * Time of test | Mauchly's W $\chi^2(2)$ = 6.575(2) | 0.679 | 0.037* | Greenhouse-Geisser correction 0,876 |
| | | *Within -subjects effect* | repeated measures ANOVA | Day | F(2,36) | 0.434 | 0.651 | ns |
| | | | repeated measures ANOVA | Day * Condition | F(2,36) | 1.894 | 0.165 | ns |
| | | | repeated measures ANOVA | Time of test | F(1,18) | 7.713 | 0.012* | * |
| | | | repeated measures ANOVA | Time of test * Condition | F(1,18) | 2.042 | 0.17 | ns |
| | | | repeated measures ANOVA | Day * Time of test | F(1.514,27.257 | 1.007 | 0.358 | ns |
| | | | repeated measures ANOVA | Day * Time of test * Condition | F(1.514,27.257 | 2.988 | 0.08 | ns |
| | | *Between-subjects effect* | | Condition | F(1,18) | 0.384 | 0.543 | ns |
| **1C.Corticosterone level on P21 (left)** | **CTL =6, ELS =6** | | Leven's test of equality of variances | | F(1,10) | 1.967 | 0.191 | equality of variances |
| | | | Student t-test | | T(10) | 0.237 | 0.817 | ns |
| **1C.Splash test-Grooming time (s) on P21 (right)** | **CTL =6, ELS =6** | | Leven's test of equality of variances | | F(1,10) | 0.769 | 0.401 | Equality of variances |
| | | | Student t-test | | T(10) | -0.663 | 0.522 | ns |
| **1D. Number of pups on P2** | **CTL = 9, ELS = 11** | | Brown-Forsythe equality of variances | | F(1,18) | 0.238 | 0.632 | Equality of variances |
| | | | Mann-Whitney rank test | | U | 43 | 0.638 | ns |
| **1F. Weight males (top)** | **CTL = 18, ELS = 25** | | Test of Sphericity | Postnatal Day | Mauchly's W $\chi^2(77)$ =636,537(77) | 1.062x10^-9 | <0.001 | Greenhouse-Geisser correction 0,281 |
| | | *Within -subjects effect* | repeated measures ANOVA mixing model | Postnatal Day | F(3.376,118.17 | 3075.266 | <0.001 | *** |
| | | | repeated measures ANOVA mixing model | Postnatal Day*Condition | F(3.376,118.17 | 7.533 | >0.001 | *** |
| | | *Between-subjects effect* | | Condition | F(1,35) | 11.257 | 0.002** | ** |
| | | | Bonferroni post hoc test | CTL vs ELS — P2 | t | 0.5 | 0.62 | ns |
| | | | Bonferroni post hoc test | P5 | t | 1.969 | 0.057 | ns |
| | | | Bonferroni post hoc test | P7 | t | 2.209 | 0.034* | * |
| | | | Bonferroni post hoc test | P12 | t | 2.304 | 0.027* | * |
| | | | Bonferroni post hoc test | P14 | t | 2.438 | 0.02* | * |
| | | | Bonferroni post hoc test | P21 | t | 2.247 | 0.031* | * |
| | | | Bonferroni post hoc test | P28 | t | 3.201 | 0.003** | ** |
| | | | Bonferroni post hoc test | P35 | t | 4.937 | <0.001 | *** |
| | | | Bonferroni post hoc test | P42 | t | 2.912 | 0.006** | ** |
| | | | Bonferroni post hoc test | P49 | t | 2.229 | 0.032* | * |
| | | | Bonferroni post hoc test | P56 | t | 3.113 | 0.004** | ** |
| | | | Bonferroni post hoc test | P63 | t | 2.828 | 0.008** | ** |
| | | | Bonferroni post hoc test | P70 | t | 2.222 | 0.033* | * |
| **1F. Weight females (bottom)** | **CTL = 18, ELS = 21** | | Test of Sphericity | Postnatal Day | Mauchly's W $\chi^2(77)$ =638,751(77) | 1.055x10^-10 | <0.001 | Greenhouse-Geisser correction 0.175 |
| | | *Within -subjects effect* | repeated measures ANOVA mixing model | Postnatal Day | F(2.100,67.197 | 1305.421 | <0.001 | *** |
| | | | repeated measures ANOVA mixing model | Postnatal Day*Condition | F(2.100,67.197 | 0.67 | 0.522 | ns |
| | | *Between-subjects effect* | | Condition | F(1,32) | 0.001 | 0.97 | ns |
| **1G. Number of USV males (top)** | **CTL = 18, ELS = 25** | | Test of Sphericity | Postnatal Day | Mauchly's W $\chi^2(5)$ =8.946(5) | 0.869 | 0.111 | Sphericity not violated |
| | | *Within -subjects effect* | repeated measures ANOVA | Postnatal Day | F(3,195) | 4.889 | 0.003** | ** |
| | | | repeated measures ANOVA | PostnatalDay * Condition | F(3,195) | 3.863 | 0.001** | ** |
| | | *Between-subjects effect* | | Condition | F(2,65) | 6.923 | 0.002** | ** |
| | | | Bonferonni post-hoc test | CTL vs ELS pre — P2 | t | 0.009 | 0.993 | ns |
| | | | Bonferonni post-hoc test | P5 | t | 1.187 | 0.479 | ns |
| | | | Bonferonni post-hoc test | P7 | t | 2.891 | 0.01* | * |
| | | | Bonferonni post-hoc test | P12 | t | 3.605 | 0.002** | ** |
| | | | Bonferonni post-hoc test | P2 | t | -1.478 | 0.327 | ns |
| | | | Bonferonni post-hoc test | P5 | t | 2.177 | 0.099 | ns |
| | | | Bonferonni post-hoc test | P7 | t | 4.324 | <0.001 | *** |
| | | | Bonferonni post-hoc test | P12 | t | 1.743 | 0.092 | ns |
| **1G. Number of USV females (bottom)** | **CTL = 18, ELS = 21** | | Test of Sphericity | Postnatal Day | Mauchly's W $\chi^2(5)$ =15.013(5) | 0.764 | 0.01* | Greenhouse-Geisser correction 0.838 |
| | | *Within -subjects effect* | repeated measures ANOVA | Postnatal Day | F(2.515,143.34 | 3.4 | 0.026* | * |
| | | | repeated measures ANOVA | PostnatalDay * Condition | F(2.515,143.34 | 2.399 | 0.04* | * |
| | | *Between-subjects effect* | | Condition | F(2,57) | 1.575 | 0.216 | ns |
| | | | Bonferonni post-hoc test | CTL vs ELS pre — P2 | t | -0.5054 | 0.615 | ns |
| | | | Bonferonni post-hoc test | P5 | t | 1.998 | 0.151 | ns |
| | | | Bonferonni post-hoc test | P7 | t | -0.44 | 0.938 | ns |
| | | | Bonferonni post-hoc test | P12 | t | 2.565 | 0.039* | * |
| | | | Bonferonni post-hoc test | P2 | t | -2.065 | 0.13 | ns |
| | | | Bonferonni post-hoc test | P5 | t | 1.663 | 0.203 | ns |
| | | | Bonferonni post-hoc test | P7 | t | 0.729 | 0.938 | ns |
| | | | Bonferonni post-hoc test | P12 | t | 1.133 | 0.283 | ns |

**Supplementary Table 2 | Detailed Statistics for Figure 2**

| Figure | N | | Test | Factor | Statistic | Statistic Value | *p* value | Notes |
|---|---|---|---|---|---|---|---|---|
| **Fig 2B. Behavioural features in males (z-score data - top)** | **CTL = 18, ELS = 25** | *Three-chamber test (social interaction ratio)* | Shapiro Wilk normality test | | W | 0.954 / 0.955 | 0.484 / 0.318 | Normal distribution |
| | | | Levenes equality of variances | | F(1,41) | 3.642 | 0.063 | Equality of variances |
| | | | Welch t test | CTL vs ELS | T(41) | 1.255 | 0.217 | ns |
| | | *Novelty supressed feeding test (Latency to eat)* | Shapiro Wilk normality test | | W | 0.95 / 0.873 | ***0.421 / 0.005*** | Normal distribution violated |
| | | | Brown-Forsythe equality of variances | | F(1,41) | 7.597 | ***0.009*** | Equality of variances violated |
| | | | Mann-Whitney rank test | CTL vs ELS | U | 189 | 0.382 | ns |
| | | *% Sucrose preference test 1%* | Shapiro Wilk normality test | | W | 0.842 / 0.834 | ***0.006** / <0.001*** | Normal distribution violated |
| | | | Brown-Forsythe equality of variances | | F(1,41) | 3.86 | 0.056 | Equality of variances |
| | | | Mann-Whitney rank test | CTL vs ELS | U | 249 | 0.567 | ns |
| | | *Forced swim test (time immobile)* | Shapiro Wilk normality test | | W | 0.93 / 0.953 | 0.193 / 0.295 | Normal distribution |
| | | | Levenes equality of variances | | F(1,41) | 0.603 | 0.442 | Equality of variances |
| | | | T-test | CTL vs ELS | T(41) | 0.318 | 0.752 | ns |
| **Fig 1F. Behavioural features in females (z-score data - bottom)** | **CTL = 18, ELS = 21** | *Three-chamber test (social interaction ratio)* | Shapiro Wilk normality test | | W | 0.885 / 0.86 | ***0.032* / 0.006*** | Normal distribution violated, qqplot (several points out) |
| | | | Levenes equality of variances | | F(1,37) | 3.411 | 0.073 | Equality of variances |
| | | | Mann-Whitney rank test | CTL vs ELS | U | 187 | 0.966 | ns |
| | | *Novelty supressed feeding test (Latency to eat)* | Shapiro Wilk normality test | | W | 0.885 / 0.907 | ***0.032* / 0.047**** | Normal distribution violated, qqplot-several points out |
| | | | Brown-Forsythe equality of variances | | F(1,37) | 2.303 | 0.138 | Equality of variances |
| | | | Mann-Whitney rank test | CTL vs ELS | U | 175 | 0.703 | ns |
| | | *% Sucrose preference test 1%* | Shapiro Wilk normality test | | W | 0.912 / 0.955 | 0.093 / 0.421 | Normal distribution |
| | | | Levenes equality of variances | | F(1,37) | 11.655 | ***0.002*** | Equality of variances violated |
| | | | Welch t test | CTL vs ELS | T(30.887) | 3.04 | ***0.005*** | ** |
| | | *Forced swim test (time immobile)* | Shapiro Wilk normality test | | W | 0.964 / 0.923 | 0.688 / 0.099 | Normal distribution |
| | | | Levenes equality of variances | | F(1,37) | 0.702 | 0.401 | Equality of variances |
| | | | T-test | CTL vs ELS | T(37) | 0.465 | 0.644 | ns |

### Mahalanobis distance correlations — males

3CH, NSF, SPT, FST composite

| Variable | Pearson *r* | *p* value[1] |
|---|---|---|
| Weight_P14 | -0.523 | **0.001**** |
| Weight_P28 | 0.485 | **0.001**** |
| Latency to eat | 0.784 | **0.001**** |
| %TH+Cfos+/TH+ Caudal Dorsal | -0.522 | **0.001**** |
| Weight_P12 | -0.429 | **0.004**** |
| Weight_P5 | -0.410 | **0.008**** |
| Weight_P56 | -0.381 | **0.012*** |
| Weight_P63 | -0.381 | **0.012*** |
| Weight_Mean | -0.381 | **0.012*** |
| USVpost_P2 | 0.353 | **0.020*** |
| % Sucrose preference | -0.328 | **0.032*** |
| %TH+Cfos+/TH+ Total Dorsal | -0.296 | 0.054 |
| Weight_P7 | -0.294 | 0.055 |
| Weight_P2 | 0.289 | 0.061 |
| USVpost_P7 | -0.281 | 0.068 |
| Density TH+Cfos+ Total Ventral | 0.274 | 0.076 |
| %TH+Cfos+/TH+ Caudal | -0.273 | 0.077 |
| Density TH+ Total | 0.268 | 0.082 |
| Density TH+Cfos+ Rostral Ventral | 0.267 | 0.084 |
| USVpre_P5 | 0.259 | 0.093 |
| Density TH+ Total Dorsal | 0.251 | 0.105 |
| CORT-post | -0.246 | 0.112 |
| Density TH+ Rostral Ventral | 0.244 | 0.115 |
| Density TH+ Caudal | 0.244 | 0.116 |
| Density TH+Cfos+ Caudal Ventral | 0.239 | 0.122 |
| Weight_P35 | -0.239 | 0.123 |
| Social index | -0.235 | 0.129 |
| USVpost_P5 | 0.233 | 0.132 |
| Density Cfos+ Total Ventral | 0.229 | 0.140 |
| Density Cfos+ Rostral Dorsal | 0.229 | 0.140 |
| Density TH+ Caudal Dorsal | 0.225 | 0.147 |
| Density TH+ Total Ventral | 0.223 | 0.150 |
| Density Cfos+ Rostral | 0.215 | 0.167 |
| Density TH+Cfos+ Total | 0.213 | 0.170 |
| Density Cfos+ Caudal Ventral | 0.207 | 0.182 |
| USVpost_Mean | 0.203 | 0.191 |
| Weight_P42 | -0.198 | 0.204 |
| Weight_P70 | -0.191 | 0.219 |
| Density Cfos+ Total Dorsal | 0.184 | 0.237 |
| Density TH+Cfos+ Caudal | 0.183 | 0.240 |
| %TH+Cfos+/TH+ Rostral | 0.181 | 0.246 |
| Density Cfos+ Total | 0.180 | 0.249 |
| Density TH+ Caudal Ventral | 0.179 | 0.250 |
| Density TH+Cfos+ Rostral | 0.177 | 0.256 |
| Weight_P21 | -0.184 | 0.276 |
| Density TH+ Rostral | 0.170 | 0.277 |
| Density Cfos+ Caudal | 0.159 | 0.307 |
| Density Cfos+ Rostral Ventral | 0.158 | 0.312 |
| Density Cfos+ Caudal Ventral | 0.153 | 0.328 |
| %TH+Cfos+/TH+ Rostral Dorsal | 0.148 | 0.343 |
| %TH+Cfos+/TH+ Total | -0.141 | 0.367 |
| Density TH+Cfos+ Total Dorsal | 0.140 | 0.372 |
| Density TH+Cfos+ Caudal Ventral | 0.123 | 0.434 |
| Weight_P49 | -0.116 | 0.459 |
| Density TH+ Rostral Dorsal | 0.114 | 0.468 |
| CORT-1h30 post | -0.094 | 0.549 |
| Density TH+Cfos+ Rostral Dorsal | 0.092 | 0.558 |
| CORT-%increase | -0.087 | 0.577 |
| USVpre_P2 | -0.083 | 0.595 |
| USVpre_P7 | -0.070 | 0.657 |
| CORT-Pre | -0.061 | 0.697 |
| USVpre_P12 | -0.059 | 0.709 |
| Time immobility | 0.053 | 0.738 |
| USVpre_Mean | 0.048 | 0.760 |
| CORT-%decrease | -0.047 | 0.766 |
| USVpost_P12 | 0.044 | 0.778 |
| %TH+Cfos+/TH+ Rostral Ventral | 0.006 | 0.968 |
| %TH+Cfos+/TH+ Caudal Ventral | 0.003 | 0.982 |
| %TH+Cfos+/TH+ Total Ventral | 0.001 | 0.988 |

[1] * $p < 0.05$ ** $p < 0.01$ *** $p < 0.001$

### Mahalanobis distance correlations — females

3CH, NSF, SPT, FST composite

| Variable | Pearson *r* | *p* value[1] |
|---|---|---|
| % Sucrose preference | -0.679 | **0.001**** |
| Density TH+ Rostral Ventral | 0.436 | **0.005**** |
| Weight_P12 | -0.422 | **0.007**** |
| Weight_P7 | -0.417 | **0.008**** |
| Density TH+Cfos+ Rostral Ventral | 0.416 | **0.008**** |
| Weight_P14 | -0.413 | **0.009**** |
| Weight_P5 | -0.405 | **0.010*** |
| Weight_P49 | -0.383 | **0.016*** |
| Density Cfos+ Rostral Ventral | 0.385 | **0.016*** |
| Weight_Mean | -0.375 | **0.019*** |
| Weight_P42 | -0.318 | **0.048*** |
| USVpost_P2 | 0.318 | **0.049*** |
| Density TH+ Rostral | 0.313 | 0.052 |
| Density TH+Cfos+ Rostral | 0.302 | 0.062 |
| Weight_P21 | -0.314 | 0.071 |
| Weight_P35 | -0.289 | 0.075 |
| Weight_P28 | -0.277 | 0.087 |
| USVpre_Mean | -0.277 | 0.088 |
| Weight_P63 | -0.267 | 0.100 |
| Density Cfos+ Rostral | 0.266 | 0.102 |
| CORT-Pre | 0.235 | 0.150 |
| CORT-%increase | 0.228 | 0.163 |
| USVpre_P12 | -0.226 | 0.166 |
| USVpost_P12 | -0.223 | 0.173 |
| Density Cfos+ Total Ventral | 0.220 | 0.175 |
| Density TH+ Total | 0.218 | 0.182 |
| Density TH+Cfos+ Total Ventral | 0.218 | 0.182 |
| USVpost_P5 | -0.215 | 0.189 |
| Social index | -0.205 | 0.210 |
| USVpre_P5 | -0.205 | 0.211 |
| CORT-%decrease | 0.200 | 0.220 |
| Density TH+ Total Ventral | 0.199 | 0.224 |
| CORT-post | -0.192 | 0.242 |
| Density TH+ Rostral Dorsal | 0.181 | 0.271 |
| Weight_P56 | -0.179 | 0.277 |
| Density TH+Cfos+ Total | 0.175 | 0.286 |
| Weight_P70 | -0.164 | 0.318 |
| Density Cfos+ Total | 0.155 | 0.345 |
| Density TH+Cfos+ Rostral Dorsal | 0.152 | 0.355 |
| Latency to eat | 0.146 | 0.376 |
| USVpre_P7 | -0.144 | 0.380 |
| USVpre_P2 | -0.144 | 0.383 |
| Density TH+Cfos+ Caudal Dorsal | -0.142 | 0.389 |
| Density TH+ Caudal Dorsal | -0.132 | 0.423 |
| Time immobility | -0.118 | 0.475 |
| %TH+Cfos+/TH+ Caudal Dorsal | -0.106 | 0.520 |
| %TH+Cfos+/TH+ Total Dorsal | -0.105 | 0.523 |
| %TH+Cfos+/TH+ Rostral | -0.105 | 0.523 |
| Density Cfos+ Caudal Ventral | 0.102 | 0.536 |
| %TH+Cfos+/TH+ Total | -0.087 | 0.600 |
| Density TH+Cfos+ Caudal Ventral | 0.085 | 0.608 |
| %TH+Cfos+/TH+ Rostral Ventral | 0.083 | 0.617 |
| Density Cfos+ Caudal Dorsal | -0.080 | 0.630 |
| Density TH+ Caudal Ventral | 0.075 | 0.649 |
| CORT-1h30 post | 0.074 | 0.654 |
| %TH+Cfos+/TH+ Rostral Dorsal | -0.067 | 0.684 |
| USVpost_P7 | -0.058 | 0.725 |
| Density Cfos+ Caudal | 0.053 | 0.747 |
| %TH+Cfos+/TH+ Caudal | -0.045 | 0.784 |
| Density Cfos+ Rostral Dorsal | 0.039 | 0.813 |
| %TH+Cfos+/TH+ Total Ventral | 0.037 | 0.825 |
| Density TH+ Total Dorsal | 0.036 | 0.830 |
| Weight_P2 | 0.027 | 0.868 |
| %TH+Cfos+/TH+ Caudal Ventral | -0.024 | 0.885 |
| Density TH+ Caudal | 0.022 | 0.896 |
| Density Cfos+ Total Dorsal | -0.022 | 0.896 |
| USVpost_Mean | -0.013 | 0.939 |
| Density TH+Cfos+ Total Dorsal | 0.004 | 0.980 |
| Density TH+Cfos+ Caudal | 0.004 | 0.980 |

[1] * $p < 0.05$ ** $p < 0.01$ *** $p < 0.001$

### Supplementary Table 4 | Detailed Statistics for Figure 4

| Figure | N | | Test | Factor | Statistic | Statistic Value | *p* value | Notes |
|---|---|---|---|---|---|---|---|---|
| **Fig 4B. Behavioural features in males (z-score data - Left)** | **CTL = 18, RES = 7, ANX = 12, ANH = 6** | *Three-chamber test (social interaction ratio)* | Shapiro Wilk on residuals | | W | | | Normal distribution, qqplot ok |
| | | | Levenes equality of variances | | F(3,39) | 0.235 | 0.871 | Equality of variances |
| | | | One-way ANOVA | Cluster groups | F(3,39) | 7.871 | <0.001 | *** |
| | | | Dunett post-hoc test | CTL vs RES | t | -1.418 | 0.392 | ns |
| | | | Dunett post-hoc test | CTL vs ANX | t | 3.79 | ***0.002**** | ** |
| | | | Dunett post-hoc test | CTL vs ANH | t | -0.27 | 0.989 | ns |
| | | *Novelty supressed feeding test (Latency to eat)* | Shapiro Wilk on residuals | | W | | | Normal distribution, qqplot ok |
| | | | Levenes equality of variances | | F(3,39) | 6.265 | ***0.001**** | Equality of variances violated |
| | | | One-way ANOVA with Welch correction | Cluster groups | F(3,12.991) | 4.327 | ***0.025**** | * |
| | | | Dunett post-hoc test | CTL vs RES | t | 0.571 | 0.910 | ns |
| | | | Dunett post-hoc test | CTL vs ANX | t | -4.217 | <0.001 | *** |
| | | | Dunett post-hoc test | CTL vs ANH | t | 0.655 | 0.872 | ns |
| | | *% Sucrose preference test 1%* | Shapiro Wilk on residuals | | W | | | Normal distribution, qqplot ok |
| | | | Levenes equality of variances | | F(3,39) | 0.59 | 0.625 | Equality of variances |
| | | | One-way ANOVA | Cluster groups | F(3,39) | 19.327 | <0.001 | *** |
| | | | Dunett post-hoc test | CTL vs RES | t | -0.951 | 0.697 | ns |
| | | | Dunett post-hoc test | CTL vs ANX | t | 0.162 | 0.998 | ns |
| | | | Dunett post-hoc test | CTL vs ANH | t | 6.98 | <0.001 | *** |
| | | *Forced swim test (time immobile)* | Shapiro Wilk on residuals | | W | | | Normal distribution, qqplot ok |
| | | | Levenes equality of variances | | F(3,39) | 0.949 | 0.426 | Equality of variances |
| | | | One-way ANOVA | Cluster groups | F(3,39) | 0.843 | 0.479 | ns |
| **Fig 4B. Behavioural features in females (z-score data - Right)** | **CTL = 18, RES = 5, ANX = 6, ANH = 10** | *Three-chamber test (social interaction ratio)* | Shapiro Wilk on residuals | | W | | | Normal distribution, qqplot ok |
| | | | Levenes equality of variances | | F(3,35) | 1.541 | 0.221 | Equality of variances |
| | | | One-way ANOVA | Cluster groups | F(3,35) | 4.43 | ***0.010**** | * |
| | | | Dunett post-hoc test | CTL vs RES | t | -1.692 | 0.255 | ns |
| | | | Dunett post-hoc test | CTL vs ANX | t | 2.706 | ***0.03**** | * |
| | | | Dunett post-hoc test | CTL vs ANH | t | -0.181 | 0.997 | ns |
| | | *Novelty supressed feeding test (Latency to eat)* | Shapiro Wilk on residuals | | W | | | Normal distribution, qqplot ok |
| | | | Levenes equality of variances | | F(3,35) | 1.851 | 0.156 | Equality of variances |
| | | | One-way ANOVA | Cluster groups | F(3,35) | 6.989 | <0.001 | *** |
| | | | Dunett post-hoc test | CTL vs RES | t | 0.204 | 0.995 | ns |
| | | | Dunett post-hoc test | CTL vs ANX | t | -4.022 | <0.001 | *** |
| | | | Dunett post-hoc test | CTL vs ANH | t | 0.768 | 0.815 | ns |
| | | *% Sucrose preference test 1%* | Shapiro Wilk on residuals | | W | | | Normal distribution, qqplot ok |
| | | | Levenes equality of variances | | F(3,35) | 0.549 | 0.652 | Equality of variances |
| | | | One-way ANOVA | Cluster groups | F(3,35) | 31.517 | <0.001 | *** |
| | | | Dunett post-hoc test | CTL vs RES | t | -1.457 | 0.374 | ns |
| | | | Dunett post-hoc test | CTL vs ANX | t | 0.786 | 0.805 | ns |
| | | | Dunett post-hoc test | CTL vs ANH | t | 8.746 | <0.001 | *** |
| | | *Forced swim test (time immobile)* | Shapiro Wilk on residuals | | W | | | Normal distribution, qqplot ok |
| | | | Levenes equality of variances | | F(3,35) | 0.261 | 0.853 | Equality of variances |
| | | | One-way ANOVA | Cluster groups | F(3,35) | 0.495 | 0.688 | ns |

**Supplementary Table 5 | Detailed Statistics for Figure 5**

| Figure | N | | Test | Factor | Statistic | Statistic Value | p value | Notes |
|---|---|---|---|---|---|---|---|---|
| **Fig.5A. Weigth (g) z-score in males left** | CTL = 18, RES = 7, ANX = 12, ANH = 6 | | Shapiro Wilk on residuals | | | | | qqplot ok |
| | | | Test of Sphericity | Postnatal Day | Mauchly's W $\chi^2(9)$ =105.187(9) | 0.06 | <0.001 | Greenhouse-Geisser correction 0,473 |
| | | *Within -subjects effect* | repeated measures ANOVA | Postnatal Day | F(1.891,73.754 | 12.236 | <0.001 | *** |
| | | | repeated measures ANOVA | Postnatal Day * Condition | F(5.673,73.754 | 5.417 | <0.001 | *** |
| | | *Between-subjects effect* | repeated measures ANOVA | Condition | F(3,39) | 9.134 | <0.001 | *** |
| | | | Bonferroni post hoc test | CTL vs ANX | t | 0.254 | 1 | ns |
| | | | Bonferroni post hoc test | CTL vs RES | t | -0.516 | 1 | ns |
| | | | Bonferroni post hoc test | CTL vs ANH | t | -1.338 | 1 | ns |
| | | | Bonferroni post hoc test | ANX vs RES | t | -0.284 | 1 | ns |
| | | | Bonferroni post hoc test | ANX vs ANH | t | -1.072 | 1 | ns |
| | | | Bonferroni post hoc test | RES vs ANH | t | -0.72 | 1 | ns |
| | | | Bonferroni post hoc test | CTL vs ANX | t | 5.885 | <0.001 | *** |
| | | | Bonferroni post hoc test | CTL vs RES | t | 1.026 | 0.27 | ns |
| | | | Bonferroni post hoc test | CTL vs ANH | t | -1.527 | 0.27 | ns |
| | | | Bonferroni post hoc test | ANX vs RES | t | -3.431 | 0.006** | ** |
| | | | Bonferroni post hoc test | ANX vs ANH | t | -5.825 | <0.001 | *** |
| | | | Bonferroni post hoc test | RES vs ANH | t | -2.302 | 0.08 | ns |
| | | | Bonferroni post hoc test | CTL vs ANX | t | 4.897 | <0.001 | *** |
| | | | Bonferroni post hoc test | CTL vs RES | t | 2.122 | 0.121 | ns |
| | | | Bonferroni post hoc test | CTL vs ANH | t | -1.223 | 0.229 | ns |
| | | | Bonferroni post hoc test | ANX vs RES | t | -1.85 | 0.144 | ns |
| | | | Bonferroni post hoc test | ANX vs ANH | t | -4.803 | <0.001 | *** |
| | | | Bonferroni post hoc test | RES vs ANH | t | -2.735 | 0.037* | * |
| | | | Bonferroni post hoc test | CTL vs ANX | t | 4.63 | <0.001 | *** |
| | | | Bonferroni post hoc test | CTL vs RES | t | 1.438 | 0.475 | ns |
| | | | Bonferroni post hoc test | CTL vs ANH | t | -0.218 | 0.828 | ns |
| | | | Bonferroni post hoc test | ANX vs RES | t | -2.281 | 0.112 | ns |
| | | | Bonferroni post hoc test | ANX vs ANH | t | -3.657 | 0.004** | ** |
| | | | Bonferroni post hoc test | RES vs ANH | t | -1.337 | 0.475 | ns |
| | | | Bonferroni post hoc test | CTL vs ANX | t | 4.285 | <0.001 | *** |
| | | | Bonferroni post hoc test | CTL vs RES | t | 1.671 | 0.323 | ns |
| | | | Bonferroni post hoc test | CTL vs ANH | t | -0.143 | 0.887 | ns |
| | | | Bonferroni post hoc test | ANX vs RES | t | -1.793 | 0.323 | ns |
| | | | Bonferroni post hoc test | ANX vs ANH | t | -3.328 | 0.01* | * |
| | | | Bonferroni post hoc test | RES vs ANH | t | -1.459 | 0.323 | ns |
| **Fig.5A. Mean Weigth P2-P14 (g) z-score in males right** | CTL = 18, RES = 7, ANX = 12, ANH = 6 | | Shapiro Wilk on residuals | | | | | qqplot ok |
| | | | Levenes equality of variances | | F(3,39) | 7.979 | <0.001 | Equality of variances violated |
| | | | One-way ANOVA Welch correction | | F(3,15.2) | 10.69 | <0.001 | *** |
| | | | Bonferroni post hoc test | CTL vs ANX | t | 4.611 | <0.001 | *** |
| | | | Bonferroni post hoc test | CTL vs RES | t | 1.422 | 0.978 | ns |
| | | | Bonferroni post hoc test | CTL vs ANH | t | -0.874 | 1 | ns |
| | | | Bonferroni post hoc test | ANX vs RES | t | -2.281 | 0.168 | ns |
| | | | Bonferroni post hoc test | ANX vs ANH | t | -4.261 | <0.001 | *** |
| | | | Bonferroni post hoc test | RES vs ANH | t | -1.879 | 0.406 | ns |
| **Fig.5B.Number of USV pre separation (n) z-score in males left** | CTL = 18, RES = 7, ANX = 12, ANH = 6 | | Shapiro Wilk on residuals | | | | | qqplot ok |
| | | | Test of Sphericity | Postnatal Day | Mauchly's W $\chi^2(5)$ =16.846(5) | 0.64 | 0.005** | Greenhouse-Geisser correction 0,756 |
| | | | repeated measures ANOVA | Postnatal Day | F(2.269,88.481 | 14.029 | <0.001 | *** |
| | | | repeated measures ANOVA | Postnatal Day * Condition | F(6.809,88.481 | 5.221 | <0.001 | *** |
| | | | repeated measures ANOVA | Condition | F(3,39) | 4.647 | 0.007** | ** |
| | | | Bonferroni post hoc test | CTL vs ANX | t | -1.102 | 1 | ns |
| | | | Bonferroni post hoc test | CTL vs RES | t | -0.312 | 1 | ns |
| | | | Bonferroni post hoc test | CTL vs ANH | t | 0.495 | 1 | ns |
| | | | Bonferroni post hoc test | ANX vs RES | t | -0.572 | 1 | ns |
| | | | Bonferroni post hoc test | ANX vs ANH | t | 0.572 | 1 | ns |
| | | | Bonferroni post hoc test | RES vs ANH | t | 0.669 | 1 | ns |
| | | | Bonferroni post hoc test | CTL vs ANX | t | -0.638 | 1 | ns |
| | | | Bonferroni post hoc test | CTL vs RES | t | -0.889 | 1 | ns |
| | | | Bonferroni post hoc test | CTL vs ANH | t | 1.059 | 1 | ns |
| | | | Bonferroni post hoc test | ANX vs RES | t | -0.332 | 1 | ns |
| | | | Bonferroni post hoc test | ANX vs ANH | t | 1.474 | 0.891 | ns |
| | | | Bonferroni post hoc test | RES vs ANH | t | 1.609 | 0.695 | ns |
| | | | Bonferroni post hoc test | CTL vs ANX | t | 1.749 | 0.528 | ns |
| | | | Bonferroni post hoc test | CTL vs RES | t | 1.816 | 0.463 | ns |
| | | | Bonferroni post hoc test | CTL vs ANH | t | 1.056 | 1 | ns |
| | | | Bonferroni post hoc test | ANX vs RES | t | 0.33 | 1 | ns |
| | | | Bonferroni post hoc test | ANX vs ANH | t | -0.309 | 1 | ns |
| | | | Bonferroni post hoc test | RES vs ANH | t | -0.559 | 1 | ns |
| | | | Bonferroni post hoc test | CTL vs ANX | t | 2.282 | 0.056 | ns |
| | | | Bonferroni post hoc test | CTL vs RES | t | 8.48 | <0.001 | *** |
| | | | Bonferroni post hoc test | CTL vs ANH | t | 2.794 | 0.024* | * |
| | | | Bonferroni post hoc test | ANX vs RES | t | 6.154 | <0.001 | *** |
| | | | Bonferroni post hoc test | ANX vs ANH | t | 0.933 | 0.357 | ns |
| | | | Bonferroni post hoc test | RES vs ANH | t | -4.422 | <0.001 | *** |
| **Fig.5B.Mean number of USV pre separation (n) z-score in males right** | CTL = 18, RES = 7, ANX = 12, ANH = 6 | | Shapiro Wilk on residuals | | | | | qqplot ok |
| | | | Levenes equality of variances | | F(3,39) | 2.96 | 0.044* | Equality of variances violated |
| | | | One-way ANOVA Welch correction | | F(3,15.586) | 4.436 | 0.019* | * |
| | | | Bonferroni post hoc test | CTL vs ANX | t | 0.734 | 1 | ns |
| | | | Bonferroni post hoc test | CTL vs RES | t | 3.316 | 0.012* | * |
| | | | Bonferroni post hoc test | CTL vs ANH | t | 2.34 | 0.147 | ns |
| | | | Bonferroni post hoc test | ANX vs RES | t | 2.53 | 0.093 | ns |
| | | | Bonferroni post hoc test | ANX vs ANH | t | 1.659 | 0.631 | ns |
| | | | Bonferroni post hoc test | RES vs ANH | t | -0.672 | 1 | ns |
| **Fig.5C.Number of USV post separation (n) z-score in males left** | CTL = 18, RES = 7, ANX = 12, ANH = 6 | | Shapiro Wilk on residuals | | | | | qqplot ok |
| | | | Test of Sphericity | Postnatal Day | Mauchly's W $\chi^2(5)$ =11.265 | 0.742 | 0.046* | Greenhouse-Geisser correction 0,869 |
| | | | repeated measures ANOVA | Postnatal Day | F(2.606,101.64 | 12.556 | <0.001 | *** |
| | | | repeated measures ANOVA | Postnatal Day * Condition | F(7.82,101.642 | 5.23 | <0.001 | *** |
| | | | repeated measures ANOVA | Condition | F(3,39) | 7.501 | <0.001 | *** |
| | | | Bonferroni post hoc test | CTL vs ANX | t | -3.181 | 0.017* | * |
| | | | Bonferroni post hoc test | CTL vs RES | t | -0.827 | 1 | ns |
| | | | Bonferroni post hoc test | CTL vs ANH | t | -0.009 | 1 | ns |
| | | | Bonferroni post hoc test | ANX vs RES | t | 1.719 | 0.375 | ns |
| | | | Bonferroni post hoc test | ANX vs ANH | t | 2.362 | 0.116 | ns |
| | | | Bonferroni post hoc test | RES vs ANH | t | 0.654 | 1 | ns |
| | | | Bonferroni post hoc test | CTL vs ANX | t | -0.839 | 1 | ns |
| | | | Bonferroni post hoc test | CTL vs RES | t | 0.159 | 1 | ns |
| | | | Bonferroni post hoc test | CTL vs ANH | t | 2.301 | 0.134 | ns |
| | | | Bonferroni post hoc test | ANX vs RES | t | 0.806 | 1 | ns |
| | | | Bonferroni post hoc test | ANX vs ANH | t | 2.794 | 0.048* | * |
| | | | Bonferroni post hoc test | RES vs ANH | t | 1.822 | 0.304 | ns |
| | | | Bonferroni post hoc test | CTL vs ANX | t | 3.696 | 0.004** | ** |
| | | | Bonferroni post hoc test | CTL vs RES | t | 2.751 | 0.045* | * |
| | | | Bonferroni post hoc test | CTL vs ANH | t | 2.642 | 0.047* | * |
| | | | Bonferroni post hoc test | ANX vs RES | t | -0.319 | 1 | ns |
| | | | Bonferroni post hoc test | ANX vs ANH | t | -0.264 | 1 | ns |
| | | | Bonferroni post hoc test | RES vs ANH | t | 0.036 | 1 | ns |
| | | | Bonferroni post hoc test | CTL vs ANX | t | -0.081 | 0.936 | ns |
| | | | Bonferroni post hoc test | CTL vs RES | t | 5.397 | <0.001 | *** |
| | | | Bonferroni post hoc test | CTL vs ANH | t | 1.813 | 0.232 | ns |
| | | | Bonferroni post hoc test | ANX vs RES | t | 5.118 | <0.001 | *** |
| | | | Bonferroni post hoc test | ANX vs ANH | t | 1.77 | 0.232 | ns |
| | | | Bonferroni post hoc test | RES vs ANH | t | -2.785 | 0.033* | * |
| **Fig.5C.Mean number of USV post separation (n) z-score in males right** | CTL = 18, RES = 7, ANX = 12, ANH = 6 | | Shapiro Wilk on residuals | | | | | qqplot ok |
| | | | Levenes equality of variances | | F(3,35) | 2.006 | 0.129 | Equality of variances |
| | | | One-way ANOVA Welch correction | | F(3,17.34) | 12.148 | <0.001 | *** |
| | | | Bonferroni post hoc test | CTL vs ANX | t | -0.91 | 1 | ns |
| | | | Bonferroni post hoc test | CTL vs RES | t | 3.244 | 0.015* | * |
| | | | Bonferroni post hoc test | CTL vs ANH | t | 2.903 | 0.036* | * |
| | | | Bonferroni post hoc test | ANX vs RES | t | 3.751 | 0.003** | ** |
| | | | Bonferroni post hoc test | ANX vs ANH | t | 3.415 | 0.009** | ** |
| | | | Bonferroni post hoc test | RES vs ANH | t | -0.138 | 1 | ns |
| **Fig.5D. Weigth (g) z-score in females left** | CTL = 18, RES = 5, ANX = 6, ANH = 10 | | Shapiro Wilk on residuals | | | | | qqplot ok |
| | | | Test of Sphericity | Postnatal Day | Mauchly's W $\chi^2(9)$ =73.876(9) | 0.11 | <0.001 | Greenhouse-Geisser correction 0,527 |
| | | *Within -subjects effect* | repeated measures ANOVA | Postnatal Day | F(2.11,73.834) | 40.497 | <0.001 | *** |
| | | | repeated measures ANOVA | Postnatal Day * Condition | F(6,3,73.834) | 10.702 | <0.001 | *** |
| | | *Between-subjects effect* | repeated measures ANOVA | Condition | F(3,35) | 6.977 | <0.001 | *** |
| | | | Bonferroni post hoc test | CTL vs ANX | t | -3.06 | 0.025* | * |
| | | | Bonferroni post hoc test | CTL vs RES | t | 0.175 | 0.924 | ns |
| | | | Bonferroni post hoc test | CTL vs ANH | t | -2.684 | 0.055 | ns |
| | | | Bonferroni post hoc test | ANX vs RES | t | 2.094 | 0.065 | ns |
| | | | Bonferroni post hoc test | ANX vs ANH | t | 0.744 | 0.924 | ns |
| | | | Bonferroni post hoc test | RES vs ANH | t | -2.094 | 0.131 | ns |
| | | | Bonferroni post hoc test | CTL vs ANX | t | -1.785 | 0.166 | ns |
| | | | Bonferroni post hoc test | CTL vs RES | t | 2.78 | 0.035* | * |
| | | | Bonferroni post hoc test | CTL vs ANH | t | 2.523 | 0.049* | ns |
| | | | Bonferroni post hoc test | ANX vs RES | t | 3.71 | 0.004** | ** |
| | | | Bonferroni post hoc test | ANX vs ANH | t | 3.556 | 0.006** | ** |
| | | | Bonferroni post hoc test | RES vs ANH | t | -0.749 | 0.459 | ns |
| | | | Bonferroni post hoc test | CTL vs ANX | t | -1.561 | 0.255 | ns |
| | | | Bonferroni post hoc test | CTL vs RES | t | 3.779 | 0.003** | ** |
| | | | Bonferroni post hoc test | CTL vs ANH | t | 3.124 | 0.011* | * |
| | | | Bonferroni post hoc test | ANX vs RES | t | 4.371 | <0.001 | *** |
| | | | Bonferroni post hoc test | ANX vs ANH | t | 3.811 | 0.003** | ** |
| | | | Bonferroni post hoc test | RES vs ANH | t | -1.238 | 0.255 | ns |
| | | | Bonferroni post hoc test | CTL vs ANX | t | -0.649 | 0.521 | ns |
| | | | Bonferroni post hoc test | CTL vs RES | t | 4.211 | 0.001** | ** |
| | | | Bonferroni post hoc test | CTL vs ANH | t | 2.791 | 0.034* | * |
| | | | Bonferroni post hoc test | ANX vs RES | t | 4.02 | 0.001** | ** |
| | | | Bonferroni post hoc test | ANX vs ANH | t | 2.724 | 0.034* | * |
| | | | Bonferroni post hoc test | RES vs ANH | t | -7.876 | 0.138 | ns |
| | | | Bonferroni post hoc test | CTL vs ANX | t | 0.216 | 0.83 | ns |
| | | | Bonferroni post hoc test | CTL vs RES | t | 4.714 | <0.001 | *** |
| | | | Bonferroni post hoc test | CTL vs ANH | t | 2.726 | 0.04* | * |
| | | | Bonferroni post hoc test | ANX vs RES | t | 3.767 | 0.003** | ** |
| | | | Bonferroni post hoc test | ANX vs ANH | t | 1.885 | 0.136 | ns |
| | | | Bonferroni post hoc test | RES vs ANH | t | -2.388 | 0.067 | ns |
| **Fig.5D. Mean Weigth P2-P14 (g) z-score in females right** | CTL = 18, RES = 5, ANX = 6, ANH = 10 | | Shapiro Wilk on residuals | | | | | qqplot ok |
| | | | Levenes equality of variances | | F(3,35) | 16.887 | <0.001 | Equality of variances violated |
| | | | One-way ANOVA Welch correction | | F(3,11.165) | 10.668 | 0.001** | ** |
| | | | Bonferroni post hoc test | CTL vs ANX | t | -1.519 | 0.826 | ns |
| | | | Bonferroni post hoc test | CTL vs RES | t | 3.514 | 0.007** | ** |
| | | | Bonferroni post hoc test | CTL vs ANH | t | 1.957 | 0.35 | ns |
| | | | Bonferroni post hoc test | ANX vs RES | t | 4.116 | 0.001** | ** |
| | | | Bonferroni post hoc test | ANX vs ANH | t | 2.882 | 0.04* | * |
| | | | Bonferroni post hoc test | RES vs ANH | t | -1.834 | 0.451 | ns |
| **Fig.5E.Number of USV pre separation (n) z-score in females left** | CTL = 18, RES = 5, ANX = 6, ANH = 10 | | Shapiro Wilk on residuals | | | | | qqplot ok |
| | | | Test of Sphericity | Postnatal Day | Mauchly's W $\chi^2(5)$ =8.343(5) | 0.781 | 0.138 | Sphericity not violated |
| | | | repeated measures ANOVA | Postnatal Day | F(3,105) | 7.802 | <0.001 | *** |
| | | | repeated measures ANOVA | Postnatal Day * Condition | F(9,105) | 3.808 | <0.001 | *** |
| | | | repeated measures ANOVA | Condition | F(3,35) | | 0.07 | ns |
| | | | Bonferroni post hoc test | CTL vs ANX | t | -0.626 | 1 | ns |
| | | | Bonferroni post hoc test | CTL vs RES | t | -1.622 | 0.683 | ns |
| | | | Bonferroni post hoc test | CTL vs ANH | t | 0.151 | 1 | ns |
| | | | Bonferroni post hoc test | ANX vs RES | t | -0.867 | 1 | ns |
| | | | Bonferroni post hoc test | ANX vs ANH | t | 0.686 | 1 | ns |
| | | | Bonferroni post hoc test | RES vs ANH | t | 1.606 | 0.683 | ns |
| | | | Bonferroni post hoc test | CTL vs ANX | t | 1.058 | 0.352 | ns |
| | | | Bonferroni post hoc test | CTL vs RES | t | 1.955 | 1 | ns |
| | | | Bonferroni post hoc test | CTL vs ANH | t | 1.103 | 1 | ns |
| | | | Bonferroni post hoc test | ANX vs RES | t | 0.808 | 1 | ns |
| | | | Bonferroni post hoc test | ANX vs ANH | t | -0.123 | 1 | ns |
| | | | Bonferroni post hoc test | RES vs ANH | t | -1.01 | 1 | ns |
| | | | Bonferroni post hoc test | CTL vs ANX | t | -0.194 | 1 | ns |
| | | | Bonferroni post hoc test | CTL vs RES | t | -3.763 | 0.003** | ** |
| | | | Bonferroni post hoc test | CTL vs ANH | t | 0.67 | 1 | ns |
| | | | Bonferroni post hoc test | ANX vs RES | t | -2.99 | 0.2 | ns |
| | | | Bonferroni post hoc test | ANX vs ANH | t | 0.689 | 1 | ns |
| | | | Bonferroni post hoc test | RES vs ANH | t | 3.955 | 0.002** | ** |
| | | | Bonferroni post hoc test | CTL vs ANX | t | 2.307 | 0.163 | ns |
| | | | Bonferroni post hoc test | CTL vs RES | t | 1.071 | 1 | ns |
| | | | Bonferroni post hoc test | CTL vs ANH | t | 1.324 | 0.971 | ns |
| | | | Bonferroni post hoc test | ANX vs RES | t | -0.902 | 1 | ns |
| | | | Bonferroni post hoc test | ANX vs ANH | t | -1.095 | 1 | ns |
| | | | Bonferroni post hoc test | RES vs ANH | t | -0.035 | 1 | ns |
| **Fig.5E.Mean number of USV pre separation (n) z-score in females right** | CTL = 18, RES = 5, ANX = 6, ANH = 10 | | Shapiro Wilk on residuals | | | | | qqplot ok |
| | | | Levenes equality of variances | | F(3,35) | 1.394 | 0.261 | Equality of variances |
| | | | One-way ANOVA | | F(3,35) | 2.568 | 0.07 | ns |
| **Fig.5F.Number of USV post separation (n) z-score in females left** | CTL = 18, RES = 5, ANX = 6, ANH = 10 | | Shapiro Wilk on residuals | | | | | qqplot ok |
| | | | Test of Sphericity | Postnatal Day | Mauchly's W $\chi^2(5)$ =17.557(5) | 0.594 | 0.004** | Greenhouse-Geisser correction 0,726 |
| | | | repeated measures ANOVA | Postnatal Day | F(2.177,76.189 | 5.307 | 0.006** | ** |
| | | | repeated measures ANOVA | Postnatal Day * Condition | F(6.5,76.189) | 2.45 | 0.028* | * |
| | | | repeated measures ANOVA | Condition | F(3,35) | 0.506 | 0.681 | ns |
| | | | Bonferroni post hoc test | CTL vs ANX | t | -1.76 | 0.436 | ns |
| | | | Bonferroni post hoc test | CTL vs RES | t | -0.889 | 1 | ns |
| | | | Bonferroni post hoc test | CTL vs ANH | t | -2.392 | 0.134 | ns |
| | | | Bonferroni post hoc test | ANX vs RES | t | 0.628 | 1 | ns |
| | | | Bonferroni post hoc test | ANX vs ANH | t | -0.22 | 1 | ns |
| | | | Bonferroni post hoc test | RES vs ANH | t | -0.902 | 1 | ns |
| | | | Bonferroni post hoc test | CTL vs ANX | t | 1.517 | 0.829 | ns |
| | | | Bonferroni post hoc test | CTL vs RES | t | 0.771 | 1 | ns |
| | | | Bonferroni post hoc test | CTL vs ANH | t | 1.204 | 1 | ns |
| | | | Bonferroni post hoc test | ANX vs RES | t | -0.537 | 1 | ns |
| | | | Bonferroni post hoc test | ANX vs ANH | t | -0.466 | 1 | ns |
| | | | Bonferroni post hoc test | RES vs ANH | t | 0.155 | 1 | ns |
| | | | Bonferroni post hoc test | CTL vs ANX | t | -0.669 | 0.72 | ns |
| | | | Bonferroni post hoc test | CTL vs RES | t | -1.734 | 0.458 | ns |
| | | | Bonferroni post hoc test | CTL vs ANH | t | 1.456 | 0.463 | ns |
| | | | Bonferroni post hoc test | ANX vs RES | t | 0.928 | 0.72 | ns |
| | | | Bonferroni post hoc test | ANX vs ANH | t | 1.723 | 0.458 | ns |
| | | | Bonferroni post hoc test | RES vs ANH | t | 2.65 | 0.072 | ns |
| | | | Bonferroni post hoc test | CTL vs ANX | t | 0.89 | 1 | ns |
| | | | Bonferroni post hoc test | CTL vs RES | t | 0.483 | 1 | *** |
| | | | Bonferroni post hoc test | CTL vs ANH | t | 0.292 | 1 | ns |
| | | | Bonferroni post hoc test | ANX vs RES | t | 0.29 | 1 | *** |
| | | | Bonferroni post hoc test | ANX vs ANH | t | -0.589 | 1 | ns |
| | | | Bonferroni post hoc test | RES vs ANH | t | -0.235 | 1 | * |
| **Fig.5F.Mean number of USV post separation (n) z-score in females right** | CTL = 18, RES = 5, ANX = 6, ANH = 10 | | Shapiro Wilk on residuals | | | | | qqplot ok |
| | | | Levenes equality of variances | | F(3,35) | 0.924 | 0.439 | Equality of variances |
| | | | One-way ANOVA | | F(3,39) | 0.506 | 0.681 | ns |

### Supplementary Table 6 | Detailed Statistics for Figure 6

| Figure | N | | Test | Factor | Statistic | Statistic Value | *p* value | Notes |
|---|---|---|---|---|---|---|---|---|
| **Fig.6A. Basal corticosterone (ng/ml) z-score in males top** | CTL = 18, RES = 7, ANX = 12, ANH = 6 | | Shapiro Wilk on residuals | | | | | qqplot ok |
| | | | Levenes equality of variances | | F(3,39) | 1.049 | 0.382 | Equality of variances |
| | | | One-way ANOVA | | F(3,39) | 8.407 | <0.001 | *** |
| | | | Bonferroni post hoc test | CTL vs ANX | t | 1.658 | 0.632 | ns |
| | | | Bonferroni post hoc test | CTL vs RES | t | 4.98 | <0.001 | *** |
| | | | Bonferroni post hoc test | CTL vs ANH | t | 1.924 | 0.37 | ns |
| | | | Bonferroni post hoc test | ANX vs RES | t | 3.364 | 0.01* | * |
| | | | Bonferroni post hoc test | ANX vs ANH | t | 0.578 | 1 | ns |
| | | | Bonferroni post hoc test | RES vs ANH | t | -2.357 | 0.141 | ns |
| **Fig.6A. Basal corticosterone (ng/ml) z-score in females bottom** | CTL = 18, RES = 5, ANX = 6, ANH = 10 | | Shapiro Wilk on residuals | | | | | qqplot ok |
| | | | Levenes equality of variances | | F(3,35) | 6.165 | 0.002** | Equality of variances violated |
| | | | One-way ANOVA Welch correction | | F(3,9.064) | 3.437 | 0.131 | ns |
| **Fig.6B. Corticosterone increase (%) z-score in males top** | CTL = 18, RES = 7, ANX = 12, ANH = 6 | | Shapiro Wilk on residuals | | | | | qqplot ok |
| | | | Levenes equality of variances | | F(3,39) | 3.12 | 0.037* | Equality of variances violated |
| | | | One-way ANOVA Welch correction | | F(3,15.021) | 4.326 | 0.022* | * |
| | | | Bonferroni post hoc test | CTL vs ANX | t | -0.954 | 1 | ns |
| | | | Bonferroni post hoc test | CTL vs RES | t | -4.617 | <0.001 | *** |
| | | | Bonferroni post hoc test | CTL vs ANH | t | -0.485 | 1 | ns |
| | | | Bonferroni post hoc test | ANX vs RES | t | -3.577 | 0.006** | ** |
| | | | Bonferroni post hoc test | ANX vs ANH | t | 0.254 | 1 | ns |
| | | | Bonferroni post hoc test | RES vs ANH | t | 3.286 | 0.13 | ns |
| **Fig.6B. Corticosterone increase (%) z-score in females bottom** | CTL = 18, RES = 5, ANX = 6, ANH = 10 | | Shapiro Wilk on residuals | | | | | qqplot ok |
| | | | Levenes equality of variances | | F(3,35) | 6.817 | <0.001 | Equality of variances violated |
| | | | One-way ANOVA Welch correction | | F(3,9.064) | 1.477 | 0.284 | ns |
| **Fig.6C. Corticosterone decrease (%) z-score in males top** | CTL = 18, RES = 7, ANX = 12, ANH = 6 | | Shapiro Wilk on residuals | | | | | qqplot ok |
| | | | Levenes equality of variances | | F(3,39) | 1.904 | 0.145 | Equality of variances |
| | | | One-way ANOVA | | F(3,39) | 0.785 | 0.509 | ns |
| **Fig.6C. Corticosterone decrease (%) z-score in females bottom** | CTL = 18, RES = 5, ANX = 6, ANH = 10 | | Shapiro Wilk on residuals | | | | | qqplot ok |
| | | | Levenes equality of variances | | F(3,35) | 2.303 | 0.094 | Equality of variances |
| | | | One-way ANOVA | | F(3,35) | 3.401 | 0.028* | * |
| | | | Bonferroni post hoc test | CTL vs ANX | t | -2.947 | 0.034* | * |
| | | | Bonferroni post hoc test | CTL vs RES | t | 0.437 | 1 | ns |
| | | | Bonferroni post hoc test | CTL vs ANH | t | -0.235 | 1 | ns |
| | | | Bonferroni post hoc test | ANX vs RES | t | 2.659 | 0.07 | ns |
| | | | Bonferroni post hoc test | ANX vs ANH | t | 2.511 | 0.101 | ns |
| | | | Bonferroni post hoc test | RES vs ANH | t | -0.572 | 1 | ns |
| **Fig.6F. TH+cfos+/TH+ z-score in males Left** | CTL = 18, RES = 7, ANX = 12, ANH = 6 | | Shapiro Wilk on residuals | | | | | qqplot ok |
| | | | Test of Sphericity | LC level | Mauchly's W ?$^2$(5) =4.547(5) | 0.886 | 0.474 | Sphericity not violated |
| | | *Within -subjects effect* | repeated measures ANOVA | LC level | F(3,117) | 2.35 | 0.076 | ns |
| | | | repeated measures ANOVA | LC level * Condition | F(9,117) | 2.147 | 0.031* | * |
| | | *Between-subjects effect* | repeated measures ANOVA | Condition | F(3,39) | 0.655 | 0.585 | ** |
| | | | Bonferonni post-hoc test conditional on LC level | CD — CTL vs ANX | t | 2.808 | 0.046* | * |
| | | | Bonferonni post-hoc test conditional on LC level | CTL vs RES | t | 0.535 | 1 | ns |
| | | | Bonferonni post-hoc test conditional on LC level | CTL vs ANH | t | -0.631 | 1 | ns |
| | | | Bonferonni post-hoc test conditional on LC level | RES vs ANX | t | -1.7 | 0.389 | ns |
| | | | Bonferonni post-hoc test conditional on LC level | ANX vs ANH | t | -2.688 | 0.053 | ns |
| | | | Bonferonni post-hoc test conditional on LC level | RES vs ANH | t | -0.962 | 1 | ns |
| **Fig.6F. TH+cfos+/TH+ z-score in females Right** | CTL = 18, RES = 5, ANX = 6, ANH = 10 | | Shapiro Wilk on residuals | | | | | qqplot ok |
| | | | Test of Sphericity | LC level | Mauchly's W ?$^2$(5) =4.710(5) | 0.87 | 0.453 | Sphericity not violated |
| | | *Within -subjects effect* | repeated measures ANOVA | LC level | F(3,105) | 5.87 | <0.001 | *** |
| | | | repeated measures ANOVA | LC level * Condition | F(9,105) | 2.525 | 0.012* | * |
| | | *Between-subjects effect* | repeated measures ANOVA | Condition | F(3,35) | 2.706 | 0.06 | ns |
| | | | Bonferonni post-hoc test conditional on LC level | CD — CTL vs ANX | | 1.344 | 0.563 | ns |
| | | | Bonferonni post-hoc test conditional on LC level | CTL vs RES | | 3.342 | <0.001 | *** |
| | | | Bonferonni post-hoc test conditional on LC level | CTL vs ANH | | 0.597 | 0.891 | ns |
| | | | Bonferonni post-hoc test conditional on LC level | RES vs ANX | | -2.579 | 0.057 | ns |
| | | | Bonferonni post-hoc test conditional on LC level | ANX vs ANH | | -0.771 | 0.891 | ns |
| | | | Bonferonni post-hoc test conditional on LC level | RES vs ANH | | -3.578 | 0.005** | ** |
