## Supplementary tables 7 to 13 - statistical analyses for supplementary Figures 1-6 for "Behavioral susceptibility and resilience following early-life stress are associated with sex-dependent locus coeruleus noradrenergic signatures in adulthood"

**Supplementary Table 7 | Detailed Statistics for Supplementary Figure 1**

| Figure | N | | Test | Factor | Statistic | Statistic Value | *p* value | Notes |
|---|---|---|---|---|---|---|---|---|
| **Supplementary 1A. Behavioural features in males (raw data - up)** | **CTL = 18, ELS = 25** | *Three-chamber test (social interaction ratio)* | Shapiro Wilk normality test | | W | 0.958 / 0.955 | 0.565 / 0.327 | Normal distribution |
| | | | Levenes equality of variances | | F(1.41) | 4.678 | 0.036* | Equality of variances violated |
| | | | Welch t test | CTL vs ELS | T(40.911) | -1.425 | 0.162 | ns |
| | | *Novelty supressed feeding test (Latency to eat)* | Shapiro Wilk normality test | | W | 0.845 / 0.907 | 0.007** / 0.026* | Normal distribution violated |
| | | | Levenes equality of variances | | F(1.41) | 0.306 | 0.583 | Equality of variances |
| | | | Mann-Whitney rank test | CTL vs ELS | U | 220.5 | 0.922 | ns |
| | | *% Sucrose preference test 1%* | Shapiro Wilk normality test | | W | 0.81 / 0.843 | 0.002** / 0.001** | Normal distribution violated |
| | | | Levenes equality of variances | | F(1.41) | 10.991 | 0.002** | Equality of variances violated |
| | | | Mann-Whitney rank test | CTL vs ELS | U | 198 | 0.518 | ns |
| | | *Forced swim test (time immobile)* | Shapiro Wilk normality test | | W | 0.966 / 0.909 | 0.72 / 0.028* | Normal distribution violated, qqplot ok |
| | | | Levenes equality of variances | | F(1.41) | 0.951 | 0.335 | Equality of variances |
| | | | T-test | CTL vs ELS | T(41) | -0.01 | 0.992 | ns |
| **Supplementary 1B. Behavioural features in females (raw data - down)** | **CTL = 18, ELS = 21** | *Three-chamber test (social interaction ratio)* | Shapiro Wilk normality test | | W | 0.931 / 0.854 | 0.206 / 0.005** | Normal distribution violated, qqplot (several points out) |
| | | | Levenes equality of variances | | F(1.37) | 0.356 | 0.554 | Equality of variances |
| | | | Mann-Whitney rank test | CTL vs ELS | U | 160 | 0.442 | ns |
| | | *Novelty supressed feeding test (Latency to eat)* | Shapiro Wilk normality test | | W | 0.888 / 0.911 | 0.036* / 0.057 | Normal distribution violated, qqplot-several points out |
| | | | Levenes equality of variances | | F(1.37) | 0.306 | 0.583 | Equality of variances |
| | | | Mann-Whitney rank test | CTL vs ELS | U | 194 | 0.899 | ns |
| | | *% Sucrose preference test 1%* | Shapiro Wilk normality test | | W | 0.914 / 0.96 | 0.102 / 0.519 | Normal distribution |
| | | | Levenes equality of variances | | F(1.37) | 10.991 | 0.002** | Equality of variances violated |
| | | | Welch t test | CTL vs ELS | T(28.809) | -2.999 | 0.005** | ** |
| | | *Forced swim test (time immobile)* | Shapiro Wilk normality test | | W | 0.949 / 0.919 | 0.414 / 0.083 | Normal distribution |
| | | | Levenes equality of variances | | F(1.37) | 0.702 | 0.407 | Equality of variances |
| | | | T-test | CTL vs ELS | T(37) | -0.669 | 0.508 | ns |
| **Supplementary 1C.Corticosterone before/after and 1h30 after FST in males(raw data - up)** | **CTL = 18, ELS = 21** | | Test of Sphericity | | Mauchly's W $\chi^2(2)$ =12.483(2) | 0.732 | 0.002** | Greenhouse-Geisser correction 0.789 |
| | | *Within -subjects effect* | repeated measures ANOVA | Sample | F(1.577, 64.665) | 166.72 | <0.001 | *** |
| | | | repeated measures ANOVA | Sample — Sample * Condition | F(1.577, 64.665) | 2.366 | 0.1 | ns |
| | | *Between-subjects effect* | repeated measures ANOVA | Sample — Condition | F(1.41) | 0.404 | 0.528 | ns |
| | | | repeated measures ANOVA | Sample — Before vs After | t | -24.335 | <0.001 | *** |
| | | | repeated measures ANOVA | Sample — Before vs 1h30 after | t | -6.984 | <0.001 | *** |
| | | | repeated measures ANOVA | Sample — After vs 1h30 after | t | 9.242 | <0.001 | *** |
| **Supplementary 1C.Corticosterone before/after and 1h30 after FST in females(raw data - down)** | **CTL = 18, ELS = 21** | | Test of Sphericity | | Mauchly's W $\chi^2(2)$ =2.894(2) | 0.923 | 0.235 | Sphericity not violated |
| | | *Within -subjects effect* | repeated measures ANOVA | Sample | F(2.74) | 241.942 | <0.001 | *** |
| | | | repeated measures ANOVA | Sample — Sample * Condition | F(2.74) | 4.546 | 0.014* | * |
| | | *Between-subjects effect* | repeated measures ANOVA | Sample — Condition | F(1.37) | 0.595 | 0.445 | ns |
| | | | Bonferroni post-hoc test | Before vs After | t | -23.091 | <0.001 | *** |
| | | | Bonferroni post-hoc test | Before vs 1h30 after | t | -13.606 | <0.001 | *** |
| | | | Bonferroni post-hoc test | After vs 1h30 after | t | 8.501 | <0.001 | *** |
| | | | Bonferroni post-hoc test conditional on Sample | Before — CTL vs ELS | t | 1.758 | 0.087 | ns |
| | | | Bonferroni post-hoc test conditional on Sample | After — CTL vs ELS | t | -1.68 | 0.101 | ns |
| | | | Bonferroni post-hoc test conditional on Sample | 1h30 after — CTL vs ELS | t | 1.779 | 0.083 | ns |
| **Supplementary 1E.Corticosterone basal level in males(raw data - up, left)** | **CTL = 18, ELS = 25** | | Shapiro Wilk normality test | | W | 0.931 / 0.943 | 0.2 / 0.172 | Normal distribution |
| | | | Levenes test of equality of variances | | F(1.41) | 15.644 | <0.001 | Equality of variances violated |
| | | | Welch t test | | T(20.031) | -3.087 | 0.006** | ** |
| **Supplementary 1E.Corticosterone increase (%) males(raw data - up, middle)** | **CTL = 18, ELS = 25** | | Shapiro Wilk normality test | | W | 0.888 / 0.974 | 0.036* / 0.746 | normal qqplot |
| | | | Levenes test of equality of variances | | F(1.41) | 2.382 | 0.13 | Equality of variances |
| | | | Student t test | | T(41) | 1.171 | 0.248 | ns |
| **Supplementary 1E.Corticosterone decrease (%) males(raw data - up, right)** | **CTL = 18, ELS = 25** | | Shapiro Wilk normality test | | W | 0.861 / 0.859 | 0.013* / 0.003** | Normal distribution violated |
| | | | Levenes test of equality of variances | | F(1.41) | 7.034 | 0.011* | Equality of variances violated |
| | | | Mann-Whitney rank test | | U | 287 | 0.131 | ns |
| **Supplementary 1F.Corticosterone basal level in females(raw data - down, left)** | **CTL = 18, ELS = 21** | | Shapiro Wilk normality test | | W | 0.888 / 0.87 | 0.036* / 0.01* | Normal distribution violated |
| | | | Levenes test of equality of variances | | F(1.37) | 13.916 | <0.001 | Equality of variances violated |
| | | | Mann-Whitney rank test | | U | 210 | 0.568 | ns |
| **Supplementary 1F.Corticosterone increase (%) in females(raw data - down,middle)** | **CTL = 18, ELS = 21** | | Shapiro Wilk normality test | | W | 0.741 / 0.819 | <0.001 / 0.001** | Normal distribution violated |
| | | | Levenes test of equality of variances | | F(1.37) | 1.418 | 0.241 | Equality of variances |
| | | | Mann-Whitney rank test | | U | 155 | 0.349 | ns |
| **Supplementary 1F.Corticosterone decrease (%) in females(raw data - down,right)** | **CTL = 18, ELS = 21** | | Shapiro Wilk normality test | | W | 0.912 / 0.969 | 0.093 / 0.717 | Normal distribution |
| | | | Levenes test of equality of variances | | F(1.37) | 0.649 | 0.425 | Equality of variances |
| | | | Student t-test | | T(37) | 2.272 | 0.029* | * |

**Supplementary Table 8 | Detailed Statistics for Supplementary Figure 2**

| Figure | N | | Test | Factor | Statistic | Statistic Value | *p* value | Notes |
|---|---|---|---|---|---|---|---|---|
| **Supplementary 2A. C-fos+ cells/µm2 in males (raw data)Up left** | **CTL = 18, ELS = 25** | | Shapiro Wilk normality test | | W | 0.951 / 0.937 | 0.441 / 0.125 | Normal distribution |
| | | | Levenes test of equality of variances | | F(1.41) | 0.904 | 0.347 | Equality of variances |
| | | | Student t-test | | T(41) | 1.473 | 0.148 | ns |
| **Supplementary 2A. C-fos+ cells/µm2 Rostral/ Caudal in males (raw data) Up right** | **CTL = 18, ELS = 25** | | Shapiro Wilk on residuals | | | | | qqplot ok |
| | | | Test of Sphericity | LC level | Mauchly's W $\chi^2(5)$ =55.148(5) | 0.249 | <0.001 | Greenhouse-Geisser correction 0.574 |
| | | *Within -subjects effect* | repeated measures ANOVA | LC level | F(1.721, 70.569) | 348.538 | <0.001 | *** |
| | | | repeated measures ANOVA | LC level * Condition | F(1.721, 70.569) | 0.287 | 0.718 | ns |
| | | *Between-subjects effect* | repeated measures ANOVA | Condition | F(1.41) | 2.558 | 0.117 | ns |
| | | | Bonferonni post-hoc test | RD vs RV | t | 20.276 | <0.001 | *** |
| | | | Bonferonni post-hoc test | RD vs CD | t | -15.203 | <0.001 | *** |
| | | | Bonferonni post-hoc test | RD vs CV | t | -14.175 | <0.001 | *** |
| | | | Bonferonni post-hoc test | RV vs CD | t | -23.639 | <0.001 | *** |
| | | | Bonferonni post-hoc test | RV vs CV | t | -27.036 | <0.001 | *** |
| **Supplementary 2A. C-fos+ cells/µm2 in females (raw data) Down left** | **CTL = 18, ELS = 21** | | Shapiro Wilk normality test | | W | 0.869 / 0.971 | ***0.017* / 0.745*** | Normal distribution violated |
| | | | Levenes test of equality of variances | | F(1.37) | 0.893 | 0.351 | Equality of variances |
| | | | Mann-Whitney rank test | | U | 204 | 0.686 | ns |
| **Supplementary 2A. C-fos+ cells/µm2 Rostral/ Caudal in females (raw data) down right** | **CTL = 18, ELS = 21** | | Shapiro Wilk on residuals | | | | | qqplot ok |
| | | | Test of Sphericity | LC level | Mauchly's W $\chi^2(5)$ =21.039(5) | 0.555 | <0.001 | Greenhouse-Geisser correction 0.746 |
| | | *Within -subjects effect* | repeated measures ANOVA | LC level | F(2.239.82.858 | 385.564 | <0.001 | *** |
| | | | repeated measures ANOVA | LC level * Condition | F(2.239.82.858 | 0.442 | 0.667 | ns |
| | | *Between-subjects effect* | repeated measures ANOVA | Condition | F(1.37) | 0.279 | 0.6 | ns |
| | | | Bonferonni post-hoc test | RD vs RV | t | 11.759 | <0.001 | *** |
| | | | Bonferonni post-hoc test | RD vs CD | t | -28.15 | <0.001 | *** |
| | | | Bonferonni post-hoc test | RD vs CV | t | -12.819 | <0.001 | *** |
| | | | Bonferonni post-hoc test | RV vs CD | t | -27.486 | <0.001 | *** |
| | | | Bonferonni post-hoc test | RV vs CV | t | -25.021 | <0.001 | *** |
| | | | Bonferonni post-hoc test | CD vs CV | t | 6.864 | <0.001 | *** |
| | | | Bonferonni post-hoc test | CD vs CV | t | 6.856 | <0.001 | *** |
| **Supplementary 2B.TH+C-fos+ /µm2 in males (raw data) up left** | **CTL = 18, ELS = 25** | | Shapiro Wilk normality test | | W | 0.955 / 0.984 | 0.507 / 0.947 | Normal distribution |
| | | | Levenes test of equality of variances | | F(1.41) | 0.348 | 0.558 | Equality of variances |
| | | | Student t-test | | T(41) | 0.785 | 0.437 | ns |
| **Supplementary 2B.TH+C-fos+ /µm2 Rostral/ Caudal in males (raw data) up right** | **CTL = 18, ELS = 25** | | Shapiro Wilk on residuals | | | | | qqplot ok |
| | | | Test of Sphericity | LC level | Mauchly's W $\chi^2(5)$ =53.177(5) | 0.262 | <0.001 | Greenhouse-Geisser correction 0.611 |
| | | *Within -subjects effect* | repeated measures ANOVA | LC level | F(1.833.75.167 | 407.040 | <0.001 | *** |
| | | | repeated measures ANOVA | LC level * Condition | F(1.833.75.167 | 0.050 | 0.940 | ns |
| | | *Between-subjects effect* | repeated measures ANOVA | Condition | F(1.41) | 0.745 | 0.393 | ns |
| | | | Bonferonni post-hoc test | RD vs RV | t | 7.767 | <0.001 | *** |
| | | | Bonferonni post-hoc test | RD vs CD | t | -19.836 | <0.001 | *** |
| | | | Bonferonni post-hoc test | RD vs CV | t | -19.811 | <0.001 | *** |
| | | | Bonferonni post-hoc test | RV vs CD | t | -23.784 | <0.001 | *** |
| | | | Bonferonni post-hoc test | RV vs CV | t | -25.597 | <0.001 | *** |
| | | | Bonferonni post-hoc test | CD vs CV | t | 5.078 | <0.001 | *** |
| **Supplementary 2B.TH+C-fos+ /µm2 in females (raw data)down left** | **CTL = 18, ELS = 21** | | Shapiro Wilk normality test | | W | 0.915 / 0.945 | 0.106 / 0.27 | Normal distribution |
| | | | Levenes test of equality of variances | | F(1.37) | 1.175 | 0.285 | Equality of variances |
| | | | Student t-test | | T(37) | 1.015 | 0.317 | ns |
| **Supplementary 2B.TH+C-fos+ /µm2 Rostral/ Caudal in females (raw data) down right** | **CTL = 18, ELS = 21** | | Shapiro Wilk on residuals | | | | | qqplot ok |
| | | | Test of Sphericity | LC level | Mauchly's W $\chi^2(5)$ =39.672(5) | 0.329 | <0.001 | Greenhouse-Geisser correction 0.631 |
| | | *Within -subjects effect* | repeated measures ANOVA | LC level | F(1.892.70.022 | 378.438 | <0.001 | *** |
| | | | repeated measures ANOVA | LC level * Condition | F(1.892.70.022 | 0.891 | 0.410 | ns |
| | | *Between-subjects effect* | repeated measures ANOVA | Condition | F(1.37) | 0.935 | 0.340 | ns |
| | | | Bonferonni post-hoc test | RD vs RV | t | 5.616 | <0.001 | *** |
| | | | Bonferonni post-hoc test | RD vs CD | t | -27.608 | <0.001 | *** |
| | | | Bonferonni post-hoc test | RD vs CV | t | -13.721 | <0.001 | *** |
| | | | Bonferonni post-hoc test | RV vs CD | t | -31.534 | <0.001 | *** |
| | | | Bonferonni post-hoc test | RV vs CV | t | -18.625 | <0.001 | *** |
| | | | Bonferonni post-hoc test | CD vs CV | t | 6.818 | <0.001 | *** |
| **Supplementary 2C.TH-C-fos+ /µm2 in males (raw data) up left** | **CTL = 18, ELS = 25** | | Shapiro Wilk normality test | | W | 0.966 / 0.914 | ***0.72 / 0.038**** | qqplot ok 2 points out |
| | | | Levenes test of equality of variances | | F(1.41) | 5.265 | ***0.027**** | Equality of variances violated |
| | | | Mann-Whitney rank test | | U | 243 | 0.927 | ns |
| **Supplementary 2C.TH-C-fos+ /µm2 Rostral/ Caudal in males (raw data) up right** | **CTL = 18, ELS = 25** | | Shapiro Wilk on residuals | | | | | qqplot ok |
| | | | Test of Sphericity | LC level | Mauchly's W $\chi^2(5)$ =52.940(5) | 0.264 | <0.001 | Greenhouse-Geisser correction 0.560 |
| | | *Within -subjects effect* | repeated measures ANOVA | LC level | F(1.681, 68.929) | 44.311 | <0.001 | *** |
| | | | repeated measures ANOVA | LC level * Condition | F(1.681, 68.929) | 0.766 | 0.448 | ns |
| | | *Between-subjects effect* | repeated measures ANOVA | Condition | F(1.41) | 1.192 | 0.281 | ns |
| | | | Bonferonni post-hoc test | RD vs RV | t | 20.463 | <0.001 | *** |
| | | | Bonferonni post-hoc test | RD vs CD | t | 0.007 | 1 | ns |
| | | | Bonferonni post-hoc test | RD vs CV | t | 3.453 | ***0.008***** | ** |
| | | | Bonferonni post-hoc test | RV vs CD | t | -8.714 | <0.001 | *** |
| | | | Bonferonni post-hoc test | RV vs CV | t | -5.906 | <0.001 | *** |
| | | | Bonferonni post-hoc test | CD vs CV | t | 5.281 | <0.001 | *** |
| **Supplementary 2C.TH-C-fos+ /µm2 in females (raw data) down left** | **CTL = 18, ELS = 21** | | Shapiro Wilk normality test | | W | 0.984 / 0.671 | 0.983 / <0.001 | Normality violated |
| | | | Levenes test of equality of variances | | F(1.37) | 0.174 | 0.679 | Equality of variances |
| | | | Mann-Whitney rank test | | U | 142 | 0.192 | ns |
| **Supplementary 2C.TH-C-fos+ /µm2 Rostral/ Caudal in females (raw data) down right** | **CTL = 18, ELS = 21** | | Shapiro Wilk on residuals | | | | | qqplot ok |
| | | | Test of Sphericity | LC level | Mauchly's W $\chi^2(5)$ =34.261(5) | 0.383 | <0.001 | Greenhouse-Geisser correction 0.672 |
| | | *Within -subjects effect* | repeated measures ANOVA | LC level | F(2.017.74.614 | 30.471 | <0.001 | *** |
| | | | repeated measures ANOVA | LC level * Condition | F(2.017.74.614 | 0.643 | 0.530 | ns |
| | | *Between-subjects effect* | repeated measures ANOVA | Condition | F(1.37) | 0.319 | 0.575 | ns |
| | | | Bonferonni post-hoc test | RD vs RV | t | 11.780 | <0.001 | *** |
| | | | Bonferonni post-hoc test | RD vs CD | t | 0.030 | 1 | ns |
| | | | Bonferonni post-hoc test | RD vs CV | t | 3.468 | ***0.008***** | ** |
| | | | Bonferonni post-hoc test | RV vs CD | t | -6.176 | <0.001 | *** |
| | | | Bonferonni post-hoc test | RV vs CV | t | -5.152 | <0.001 | *** |
| | | | Bonferonni post-hoc test | CD vs CV | t | 3.793 | ***0.003***** | ** |
| **Supplementary 2D.% TH+C-fos+ /TH+ in males (raw data) up left** | **CTL = 18, ELS = 25** | | Shapiro Wilk normality test | | W | 0.918 / 0.981 | 0.121 / 0.899 | Normal distribution |
| | | | Levenes test of equality of variances | | F(1.41) | 0.784 | 0.381 | Equality of variances |
| | | | Student t-test | | T(41) | 0.092 | 0.927 | ns |
| **Supplementary 2D.% TH+C-fos+ /TH+ Rostral/ Caudal in males (raw data) up right** | **CTL = 18, ELS = 25** | | Shapiro Wilk on residuals | | | | | qqplot normal |
| | | | Test of Sphericity | LC level | Mauchly's W $\chi^2(5)$ =7.278(5) | 0.833 | 0.201 | Sphericity not violated |
| | | *Within -subjects effect* | repeated measures ANOVA | LC level | F(3.123) | 198.968 | <0.001 | *** |
| | | | repeated measures ANOVA | LC level * Condition | F(3.123) | 2.559 | 0.058 | ns |
| | | *Between-subjects effect* | repeated measures ANOVA | Condition | F(1.41) | 1.656x10^-5 | 0.997 | ns |
| | | | Bonferonni post-hoc test | RD vs RV | t | 4.645 | <0.001 | *** |
| | | | Bonferonni post-hoc test | RD vs CD | t | -1.979 | 0.327 | ns |
| | | | Bonferonni post-hoc test | RD vs CV | t | -0.401 | 1 | ns |
| | | | Bonferonni post-hoc test | RV vs CD | t | -8.527 | <0.001 | *** |
| | | | Bonferonni post-hoc test | RV vs CV | t | -6.519 | <0.001 | *** |
| | | | Bonferonni post-hoc test | CD vs CV | t | 1.847 | 0.432 | ns |
| **Supplementary 2D.% TH+C-fos+ /TH+ in females (raw data) down left** | **CTL = 18, ELS = 21** | | Shapiro Wilk normality test | | W | 0.954 / 0.885 | ***0.49 / 0.018**** | Normal distribution, several points out qqplot |
| | | | Levenes test of equality of variances | | F(1.37) | 0.089 | 0.767 | Equality of variances |
| | | | Mann-Whitney rank test | | U | 230 | 0.257 | ns |
| **Supplementary 2D.% TH+C-fos+ /TH+ Rostral/ Caudal in females (raw data) down right** | **CTL = 18, ELS = 21** | | Shapiro Wilk on residuals | | | | | qqplot ok, little deviation |
| | | | Test of Sphericity | LC level | Mauchly's W $\chi^2(5)$ =6.728(5) | 0.828 | 0.242 | Greenhouse-Geisser correction 0.903 |
| | | *Within -subjects effect* | repeated measures ANOVA | LC level | F(2.708.100.19 | 34.353 | <0.001 | *** |
| | | | repeated measures ANOVA | LC level * Condition | F(2.708.100.19 | 0.810 | 0.480 | ns |
| | | *Between-subjects effect* | repeated measures ANOVA | Condition | F(1.37) | 1.506 | 0.228 | ns |
| | | | Bonferonni post-hoc test | RD vs RV | t | 7.821 | <0.001 | *** |
| | | | Bonferonni post-hoc test | RD vs CD | t | 0.259 | 1 | ns |
| | | | Bonferonni post-hoc test | RD vs CV | t | 3.709 | ***0.004***** | ** |
| | | | Bonferonni post-hoc test | RV vs CD | t | -8.608 | <0.001 | *** |
| | | | Bonferonni post-hoc test | RV vs CV | t | -4.516 | <0.001 | *** |
| | | | Bonferonni post-hoc test | CD vs CV | t | 4.304 | <0.001 | *** |

**Supplementary Table 9 | Detailed Statistics for Supplementary Figure 3**

| Figure | N | Test | Factor | Statistic | Statistic Value | *p* value | Notes |
|---|---|---|---|---|---|---|---|
| **Supplementary 3A. Basal Corticosterone in males (z-score)** | **CTL = 18, ELS = 25** | Shapiro Wilk normality test | | W | 0.925 / 0.967 | 0.156 / 0.574 | Normal distribution |
| | | Levenes test of equality of variances | | F(1.41) | 0.386 | 0.538 | Equality of variances |
| | | Student t-test | | T(41) | 3.302 | ***0.002*** | ** |
| **Supplementary 3A. Corticosterone post FST in males (z-score)** | **CTL = 18, ELS = 25** | Shapiro Wilk normality test | | W | 0.958 / 0.947 | 0.557 / 0.213 | Normal distribution |
| | | Levenes test of equality of variances | | F(1.41) | 2.673 | 0.11 | Equality of variances |
| | | Student t-test | | T(41) | 1.059 | 0.296 | ns |
| **Supplementary 3A. Corticosterone 1h30 post FST in males (z-score)** | **CTL = 18, ELS = 25** | Shapiro Wilk normality test | | W | 0.944 / 0.332 | ***0.878 / 0.006*** | Normal distribution violated |
| | | Levenes test of equality of variances | | F(1.41) | 5.313 | ***0.026**** | Equality of variances violated |
| | | Mann-Whitney rank test | | U | 231 | 0.893 | ns |
| **Supplementary 3A. Corticosterone increase (%) in males (z-score)** | **CTL = 18, ELS = 25** | Shapiro Wilk normality test | | W | 0.974 / 0.832 | 0.866 / <0.001 | Normal distribution violated |
| | | Levenes test of equality of variances | | F(1.41) | 12.987 | <0.001 | Equality of variances violated |
| | | Mann-Whitney rank test | | U | 164 | 0.138 | ns |
| **Supplementary 3A. Corticosterone decrease (%) in males (z-score)** | **CTL = 18, ELS = 25** | Shapiro Wilk normality test | | W | 0.943 / 0.59 | 0.331 / <0.001 | Normal distribution violated |
| | | Levenes test of equality of variances | | F(1.41) | 2.887 | 0.097 | Equality of variances |
| | | Mann-Whitney rank test | | U | 218 | 0.874 | ns |
| **Supplementary 3B. Basal Corticosterone in females (z-score)** | **CTL = 18, ELS = 21** | Shapiro Wilk normality test | | W | 0.944 / 0.868 | ***0.339 / 0.009*** | Normal distribution violated |
| | | Levenes test of equality of variances | | F(1.37) | 25.808 | <0.001 | Equality of variances violated |
| | | Mann-Whitney rank test | | U | 150 | 0.282 | ns |
| **Supplementary 3B. Corticosterone post FST in females (z-score)** | **CTL = 18, ELS = 21** | Shapiro Wilk normality test | | W | 0.949 / 0.958 | 0.41 / 0.484 | Normal distribution |
| | | Levenes test of equality of variances | | F(1.37) | 0.133 | 0.718 | Equality of variances |
| | | Student t-test | | T(37) | 0.805 | 0.426 | ns |
| **Supplementary 3B. Corticosterone 1h30 post FST in females (z-score)** | **CTL = 18, ELS = 21** | Shapiro Wilk normality test | | W | 0.91 / 0.953 | 0.087 / 0.395 | Normal distribution |
| | | Levenes test of equality of variances | | F(1.37) | 0.893 | 0.177 | Equality of variances |
| | | Student t-test | | T(37) | -0.657 | 0.516 | ns |
| **Supplementary 3B. Corticosterone increase (%) in females (z-score)** | **CTL = 18, ELS = 21** | Shapiro Wilk normality test | | W | 0.933 / 0.783 | 0.216 / <0.001 | Normal distribution violated |
| | | Levenes test of equality of variances | | F(1.37) | 9.706 | ***0.004*** | Equality of variances violated |
| | | Mann-Whitney rank test | | U | 238 | 0.174 | ns |
| **Supplementary 3B. Corticosterone decrease (%) in females (z-score)** | **CTL = 18, ELS = 21** | Shapiro Wilk normality test | | W | 0.923 / 0.979 | 0.147 / 0.913 | Normal distribution |
| | | Levenes test of equality of variances | | F(1.37) | 4.293 | ***0.045**** | Equality of variances |
| | | Welch t test | | T(32.127) | -1.159 | 0.255 | ns |

**Supplementary Table 10 | Detailed Statistics for Supplementary Figure 4**

| Figure | N | | Test | Factor | Statistic | Statistic Value | *p* value | Notes |
|---|---|---|---|---|---|---|---|---|
| **Supplementary 4A. TH+cFos+/µm2 in males (z-score) left** | **CTL = 18, ELS = 25** | | Shapiro Wilk on residuals | | | | | qqplot ok |
| | | | Test of Sphericity | LC level | Mauchly's W $\chi^2(5)$ =13.657(5) | 0.709 | **_0.018*_** | Greenhouse-Geisser correction 0.838 |
| | | *Within -subjects effect* | repeated measures ANOVA | LC level | F(2.515.103.21 | 0.916 | 0.435 | ns |
| | | | repeated measures ANOVA | LC level * Condition | F(2.515.103.21 | 0.916 | 0.422 | ns |
| | | *Between-subjects effect* | repeated measures ANOVA | Condition | F(1.41) | 5.968 | **_0.019*_** | * |
| | | | Bonferonni post-hoc test conditional on LC level | CTL vs ELS — RD | t | -1.648 | 0.107 | ns |
| | | | Bonferonni post-hoc test conditional on LC level | CTL vs ELS — RV | t | -2.371 | **_0.023*_** | * |
| | | | Bonferonni post-hoc test conditional on LC level | CTL vs ELS — CD | t | -1.555 | 0.128 | ns |
| | | | Bonferonni post-hoc test conditional on LC level | CTL vs ELS — CV | t | -1.479 | 0.147 | ns |
| **Supplementary 4A. (TH+cFos+)/TH+ (%) in males (z-score) right** | **CTL = 18, ELS = 25** | | Shapiro Wilk on residuals | | | | | qqplot ok |
| | | | Test of Sphericity | LC level | Mauchly's W $\chi^2(5)$ =4.927(5) | 0.883 | 0.425 | Sphericity not violated |
| | | *Within -subjects effect* | repeated measures ANOVA | LC level | F(3.123) | 1.862 | 0.145 | ns |
| | | | repeated measures ANOVA | LC level * Condition | F(3.123) | 1.862 | 0.145 | ns |
| | | *Between-subjects effect* | repeated measures ANOVA | Condition | F(1.41) | 0.62 | 0.435 | ns |
| **Supplementary 4B. TH+cFos+/µm2 in females (z-score) left** | **CTL = 18, ELS = 21** | | Shapiro Wilk on residuals | | | | | qqplot ok |
| | | | Test of Sphericity | LC level | Mauchly's W $\chi^2(5)$ =3.061(5) | 0.918 | 0.691 | Sphericity not violated |
| | | *Within -subjects effect* | repeated measures ANOVA | LC level | F(3.111) | 1.827 | 0.146 | ns |
| | | | repeated measures ANOVA | LC level * Condition | F(3.111) | 1.827 | 0.146 | ns |
| | | *Between-subjects effect* | repeated measures ANOVA | Condition | F(1.37) | 2.34 | 0.135 | ns |
| **Supplementary 4B. (TH+cFos+)/TH+ (%) in females (z-score) right** | **CTL = 18, ELS = 21** | | Shapiro Wilk on residuals | | | | | qqplot ok |
| | | | Test of Sphericity | LC level | Mauchly's W $\chi^2(5)$ =16.622(5) | 0.628 | **_0.005**_** | Greenhouse-Geisser correction 0.817 |
| | | *Within -subjects effect* | repeated measures ANOVA | LC level | F(2.451.90.676 | 2.499 | 0.076 | ns |
| | | | repeated measures ANOVA | LC level * Condition | F(2.451.90.676 | 2.498 | 0.076 | ns |
| | | *Between-subjects effect* | repeated measures ANOVA | Condition | F(1.37) | 0.073 | 0.788 | ns |

### Supplementary Table 11 | Detailed Statistics for Supplementary Figure 5

| Figure | N | | Test | Factor | Statistic | Statistic Value | *p* value | Notes |
|---|---|---|---|---|---|---|---|---|
| **Supplementary 5A. Social interaction index depending on cycle (z-score) left** | CTL = 18, ELS = 21 | | Shapiro Wilk on residuals | | | | | qqplot ok |
| | | | Levenes equality of variances | | F(3.35) | 2.665 | 0.063 | Equality of variances |
| | | *Three chamber test* | ANOVA | Cycle Three chamber | F(1.35) | 0.044 | 0.836 | ns |
| | | | ANOVA | Cycle Three chamber * Condition | F(1.35) | 0.505 | 0.482 | ns |
| | | *Between-subjects effect* | ANOVA | Condition | F(1.35) | 0.0358 | 0.847 | ns |
| **Supplementary 5A. Latency to eat depending on cycle (z-score) middle** | CTL = 18, ELS = 21 | | Shapiro Wilk on residuals | | | | | qqplot ok |
| | | | Levenes equality of variances | | F(3.35) | 0.277 | 0.842 | Equality of variances |
| | | *Novelty supressed feeding test (NSFT)* | ANOVA | Cycle NSFT | F(1.35) | 0.405 | 0.529 | ns |
| | | | ANOVA | Cycle NSFT * Condition | F(1.35) | 5.725 | 0.022* | ns |
| | | *Between-subjects effect* | ANOVA | Condition | F(1.35) | 2.371 | 0.133 | ns |
| | | | Bonferonni post-hoc test conditional on NSFT cycle | P/E — CTL vs ELS | t | -2.508 | 0.017* | * |
| | | | Bonferonni post-hoc test conditional on NSFT cycle | D/M — CTL vs ELS | t | 0.687 | 0.496 | ns |
| **Supplementary 5A. Immobile time (s) depending on cycle (z-score) right** | CTL = 18, ELS = 21 | | Shapiro Wilk on residuals | | | | | qqplot ok |
| | | | Levenes equality of variances | | F(3.35) | 1.485 | 0.236 | Equality of variances |
| | | *Forced swim test (FST)* | ANOVA | Cycle FST | F(1.35) | 0.007 | 0.934 | ns |
| | | | ANOVA | Cycle FST * Condition | F(1.35) | 2.055 | 0.161 | ns |
| | | *Between-subjects effect* | ANOVA | Condition | F(1.35) | 0.254 | 0.618 | ns |
| **Supplementary 5B. Social interaction index depending on cycle (Mahalanobis distance) left** | CTL = 18, ELS = 21 | | Shapiro Wilk on residuals | | | | | qqplot ok |
| | | | Levenes equality of variances | | F(3.35) | 8.009 | < .001 | Equality of variances [illegible] |
| | | *Three chamber test* | ANOVA | Cycle Three chamber | F(1.35) | 0.179 | 0.675 | ns |
| | | | ANOVA | Cycle Three chamber * Condition | F(1.35) | 1.268 | 0.268 | ns |
| | | *Between-subjects effect* | ANOVA | Condition | F(1.35) | 19.526 | < .001 | *** |
| | | | Bonferonni Post hoc test | CTL vs ELS | t | -4.419 | <0.001 | *** |
| **Supplementary 5B. Latency to eat depending on cycle (Mahalanobis distance) middle** | CTL = 18, ELS = 21 | | Shapiro Wilk on residuals | | | | | qqplot ok |
| | | | Levenes equality of variances | | F(3.35) | 3.536 | 0.024* | Equality of variances [illegible] |
| | | *Novelty supressed feeding test (NSFT)* | ANOVA | Cycle NSFT | F(1.35) | 0.148 | 0.702 | ns |
| | | | ANOVA | Cycle NSFT * Condition | F(1.35) | 1.042 | 0.314 | ns |
| | | *Between-subjects effect* | ANOVA | Condition | F(1.35) | 20.101 | < .001 | *** |
| | | | Bonferonni Post hoc test | CTL vs ELS | t | 4.901 | <0.001 | *** |
| **Supplementary 5B. Immobile time (s) depending on cycle (Mahalanobis distance) right** | CTL = 18, ELS = 21 | | Shapiro Wilk on residuals | | | | | qqplot ok |
| | | | Levenes equality of variances | | F(3.35) | 4.311 | 0.011* | Equality of variances [illegible] |
| | | *Forced swim test (FST)* | ANOVA | Cycle FST | F(1.35) | 0.069 | 0.795 | ns |
| | | | ANOVA | Cycle FST * Condition | F(1.35) | 0.250 | 0.620 | ns |
| | | *Between-subjects effect* | ANOVA | Condition | F(1.35) | 22.059 | < .001 | *** |
| | | | Bonferonni Post hoc test | CTL vs ELS | t | 4.901 | <0.001 | *** |
| **Supplementary 5C. Binomial test against 0.5 for 3CH, NSF and FST cycle depending on group** | CTL = 18 | *FST cycle* | Binomial test | CTL P/E | Proportion | 11/18 (0.611) | 0.481 | ns |
| | | *FST cycle* | Binomial test | CTL D/M | Proportion | 7/18 (0.389) | 0.481 | ns |
| | | *3CH cycle* | Binomial test | CTL P/E | Proportion | 10/18 (0.556) | 0.815 | ns |
| | | *3CH cycle* | Binomial test | CTL D/M | Proportion | 8/18 (0.444) | 0.815 | ns |
| | | *NSF cycle* | Binomial test | CTL P/E | Proportion | 7/18 (0.389) | 0.481 | ns |
| | | *NSF cycle* | Binomial test | CTL D/M | Proportion | 11/18 (0.611) | 0.481 | ns |
| | | *FST cycle* | Binomial test | RES D/M | Proportion | 3/5 (0.600) | 1.000 | ns |
| | | *FST cycle* | Binomial test | RES P/E | Proportion | 2/5 (0.400) | 1.000 | ns |
| | | *3CH cycle* | Binomial test | RES P/E | Proportion | 3/5 (0.600) | 1.000 | ns |
| | | *3CH cycle* | Binomial test | RES D/M | Proportion | 2/5 (0.400) | 1.000 | ns |
| | | *NSF cycle* | Binomial test | RES D/M | Proportion | 2/5 (0.400) | 1.000 | ns |
| | | *NSF cycle* | Binomial test | RES P/E | Proportion | 3/5 (0.600) | 1.000 | ns |
| | | *FST cycle* | Binomial test | ANX D/M | Proportion | 3/6 (0.500) | 1.000 | ns |
| | | *FST cycle* | Binomial test | ANX P/E | Proportion | 3/6 (0.500) | 1.000 | ns |
| | | *3CH cycle* | Binomial test | ANX P/E | Proportion | 4/6 (0.667) | 0.687 | ns |
| | | *3CH cycle* | Binomial test | ANX D/M | Proportion | 2/6 (0.333) | 0.687 | ns |
| | | *NSF cycle* | Binomial test | ANX D/M | Proportion | 3/6 (0.500) | 1.000 | ns |
| | | *NSF cycle* | Binomial test | ANX P/E | Proportion | 3/6 (0.500) | 1.000 | ns |
| | | *NSF cycle* | Binomial test | ANH D/M | Proportion | 8/10 (0.800) | 0.109 | ns |
| | | *NSF cycle* | Binomial test | ANH P/E | Proportion | 2/10 (0.200) | 0.109 | ns |
| | | *3CH cycle* | Binomial test | ANH P/E | Proportion | 7/10 (0.700) | 0.344 | ns |
| | | *3CH cycle* | Binomial test | ANH D/M | Proportion | 3/10 (0.300) | 0.344 | ns |
| | | *NSF cycle* | Binomial test | ANH D/M | Proportion | 8/10 (0.800) | 0.109 | ns |
| | | *NSF cycle* | Binomial test | ANH P/E | Proportion | 2/10 (0.200) | 0.109 | ns |
| **Supplementary 5C. Contingency tables for 3CH, NSF and FST cycle depending on group** | CTL = 18, RES = 7, ANX = 12, ANH = 6 | *FST cycle* | Chi-squared test | cluster k-means | $\chi^2(3)$ | 4.785 | 0.214 | ns |
| | | *3CH cycle* | Chi-squared test | cluster k-means | $\chi^2(3)$ | 0.646 | 0.886 | ns |
| | | *NSF cycle* | Chi-squared test | cluster k-means | $\chi^2(3)$ | 2.759 | 0.430 | ns |

### Supplementary Table 12 | Detailed Statistics for Supplementary Figure 6

| Figure | N | | Test | Factor | Statistic | Statistic Value | *p* value | Notes |
|---|---|---|---|---|---|---|---|---|
| **Supplementary 6A. Basal corticosterone depending on cycle FST (z-score) left** | **CTL = 18, ELS = 21** | | Shapiro Wilk on residuals | | | | | qqplot ok |
| | | | Levenes equality of variances | | F(3,35) | 6.944 | < .001 | Equality of variances violated |
| | | *FST* | ANOVA | Cycle FST | F(1,35) | 0.554 | 0.462 | ns |
| | | | ANOVA | Cycle FST — Cycle FST * Condition | F(1,35) | 0.941 | 0.339 | ns |
| | | *Between-subjects effect* | ANOVA | Cycle FST — Condition | F(1,35) | 6.974 | ***0.012**** | * |
| | | | Bonferonni Post hoc test on Condition | Cycle FST — CTL vs ELS | t | -2.641 | ***0.012**** | * |
| **Supplementary 6A. Corticosterone post FST depending on cycle (z-score) left** | **CTL = 18, ELS = 21** | | Shapiro Wilk on residuals | | | | | qqplot ok |
| | | | Levenes equality of variances | | F(3,35) | 0.431 | 0.732 | Equality of variances |
| | | *FST* | ANOVA | Cycle FST | F(1,35) | 0.002 | 0.966 | ns |
| | | | ANOVA | Cycle FST — Cycle FST * Condition | F(1,35) | 1.092 | 0.303 | ns |
| | | *Between-subjects effect* | ANOVA | Cycle FST — Condition | F(1,35) | 0.655 | 0.424 | ns |
| **Supplementary 6A. Corticosterone 1h30 post FST depending on cycle (z-score) middle** | **CTL = 18, ELS = 21** | | Shapiro Wilk on residuals | | | | | qqplot ok |
| | | | Levenes equality of variances | | F(3,35) | 1.025 | 0.394 | Equality of variances |
| | | *FST* | ANOVA | Cycle FST | F(1,35) | 0.639 | 0.429 | ns |
| | | | ANOVA | Cycle FST — Cycle FST * Condition | F(1,35) | 0.396 | 0.533 | ns |
| | | *Between-subjects effect* | ANOVA | Cycle FST — Condition | F(1,35) | 0.773 | 0.385 | ns |
| **Supplementary 6A. Corticosterone increase (%) post FST depending on cycle (z-score) right** | **CTL = 18, ELS = 21** | | Shapiro Wilk on residuals | | | | | qqplot ok |
| | | | Levenes equality of variances | | F(3,35) | 4.582 | ***0.006***** | Equality of variances violated |
| | | *FST* | ANOVA | Cycle FST | F(1,35) | 1.755 | 0.194 | ns |
| | | | ANOVA | Cycle FST — Cycle FST * Condition | F(1,35) | 2.309 | 0.138 | ns |
| | | *Between-subjects effect* | ANOVA | Cycle FST — Condition | F(1,35) | 0.205 | 0.654 | ns |
| **Supplementary 6A. Corticosterone decrease (%) 1h30 post FST depending on cycle (z-score) right** | **CTL = 18, ELS = 21** | | Shapiro Wilk on residuals | | | | | qqplot ok |
| | | | Levenes equality of variances | | F(3,35) | 1.742 | 0.176 | Equality of variances |
| | | *FST* | ANOVA | Cycle FST | F(1,35) | 0.222 | 0.641 | ns |
| | | | ANOVA | Cycle FST — Cycle FST * Condition | F(1,35) | 0.654 | 0.424 | ns |
| | | *Between-subjects effect* | ANOVA | Cycle FST — Condition | F(1,35) | 1.506 | 0.228 | ns |
| **Supplementary 6B. TH+cfos+/µm2 depending on FST cycle (z-score) left** | **CTL = 18, ELS = 21** | | Shapiro Wilk on residuals | | | | | qqplot ok |
| | | | Test of Sphericity | LC level | Mauchly's W ?$^2$(5) =13.950(5) | 0.661 | ***0.016**** | Greenhouse-Geisser correction 0.820 |
| | | *Within -subjects effect* | repeated measures ANOVA | LC level | F(2.461,86.143 | 3.677 | ***0.022**** | * |
| | | | repeated measures ANOVA | LC level — LC level * Condition | F(2.461,86.143 | 4.223 | ***0.012**** | * |
| | | | repeated measures ANOVA | LC level — LC level * Cycle FST | F(2.461,86.143 | 3.117 | ***0.039**** | * |
| | | | repeated measures ANOVA | LC level — LC level * Condition*Cycle FST | F(2.461,86.143 | 1.247 | 0.297 | ns |
| | | *Between-subjects effect* | repeated measures ANOVA | LC level — Condition | F(1,35) | 0.231 | 0.634 | ns |
| | | | repeated measures ANOVA | LC level — Cycle FST | F(1,35) | 0.923 | 0.343 | ns |
| | | | repeated measures ANOVA | LC level — Condition*Cycle FST | F(1,35) | 0.654 | 0.424 | ns |
| | | | Bonferonni post-hoc test LC level* Condition conditional on LC level | CTL vs ELS — RD | t | 0.346 | 0.732 | ns |
| | | | Bonferonni post-hoc test LC level* Condition conditional on LC level | CTL vs ELS — RV | t | -1.603 | 0.118 | ns |
| | | | Bonferonni post-hoc test LC level* Condition conditional on LC level | CTL vs ELS — CD | t | 2.63 | ***0.013**** | * |
| | | | Bonferonni post-hoc test LC level* Condition conditional on LC level | CTL vs ELS — CV | t | 0.388 | 0.700 | ns |
| | | | LC level* Cycle FST conditional on LC level | P/E vs M/D — RD | t | -0.068 | 0.946 | ns |
| | | | LC level* Cycle FST conditional on LC level | P/E vs M/D — RV | t | 0.972 | 0.338 | ns |
| | | | LC level* Cycle FST conditional on LC level | P/E vs M/D — CD | t | -2.313 | ***0.027**** | * |
| | | | LC level* Cycle FST conditional on LC level | P/E vs M/D — CV | t | -1.309 | 0.199 | ns |
| | | | LC level* Cycle FST conditional on Cycle FST | P/E — RD vs CD | t | 3.123 | ***0.022**** | * |
| | | | LC level* Cycle FST conditional on Cycle FST | P/E — RV vs CD | t | 4.144 | ***0.001***** | ** |
| | | | Condition*LC level, conditional on Condition | ELS — RD vs RV | t | -2.951 | ***0.034**** | * |
| | | | Condition*LC level, conditional on Condition | ELS — RD vs CD | t | 3.344 | ***0.012**** | * |
| | | | Condition*LC level, conditional on Condition | ELS — RV vs CD | t | 4.93 | < .001 | *** |
| | | | Condition*LC level, conditional on Condition | ELS — RV vs CV | t | 2.424 | 0.124 | ns |
| | | | Condition*LC level, conditional on Condition | ELS — CD vs CV | t | -2.197 | 0.208 | ns |
| **Supplementary 6B. TH+cfos+/TH+ (%) depending on FST cycle (z-score) right** | **CTL = 18, ELS = 21** | | Shapiro Wilk on residuals | | | | | qqplot ok |
| | | | Test of Sphericity | LC level | Mauchly's W ?$^2$(5) =3.457(5) | 0.903 | 0.630 | Sphericity not violated |
| | | *Within -subjects effect* | repeated measures ANOVA | LC level | F(3,105) | 2.545 | 0.060 | ns |
| | | | repeated measures ANOVA | LC level — LC level * Condition | F(3,105) | 3.121 | ***0.029**** | * |
| | | | repeated measures ANOVA | LC level — LC level * Cycle FST | F(3,105) | 2.378 | 0.074 | ns |
| | | | repeated measures ANOVA | LC level — LC level * Condition*Cycle FST | F(3,105) | 0.634 | 0.595 | ns |
| | | *Between-subjects effect* | repeated measures ANOVA | LC level — Condition | F(1,35) | 4.212 | ***0.048**** | *, with precaution |
| | | | repeated measures ANOVA | LC level — Cycle FST | F(1,35) | 2.358 | 0.134 | ns |
| | | | repeated measures ANOVA | LC level — Condition*Cycle FST | F(1,35) | 2.768 | 0.105 | ns |
| | | | Bonferonni post-hoc test LC level* Condition conditional on LC level | CTL vs ELS — RD | t | 1.737 | 0.091 | ns |
| | | | Bonferonni post-hoc test LC level* Condition conditional on LC level | CTL vs ELS — RV | t | -0.132 | 0.895 | ns |
| | | | Bonferonni post-hoc test LC level* Condition conditional on LC level | CTL vs ELS — CD | t | 2.986 | ***0.005***** | ** |
| | | | Bonferonni post-hoc test LC level* Condition conditional on LC level | CTL vs ELS — CV | t | 1.25 | 0.220 | ns |

**Supplementary Table 13 | Detailed Statistics for Supplementary Figure 7**

| Figure | N | | Test | Factor | Statistic | Statistic Value | *p* value | Notes |
|---|---|---|---|---|---|---|---|---|
| **Supplementary 7A.Corticosterone post-FST in males (z-score) top** | **CTL = 18, RES = 7, ANX = 12, ANH = 6** | | Shapiro Wilk on residuals | | | | | qqplot ok |
| | | | Levenes equality of variances | | F(3.39) | 3.042 | 0.04* | Equality of variances violated |
| | | | One-way ANOVA Welch correction | kmeans clusters | F(3.15.601) | 1.815 | 0.186 | ns |
| **Supplementary 7A.Corticosterone post-FST in females (z-score) bottom** | **CTL = 18, RES = 5, ANX = 6, ANH = 10** | | Shapiro Wilk on residuals | | | | | qqplot ok |
| | | | Levenes equality of variances | | F(3.35) | 0.847 | 0.477 | Equality of variances |
| | | | One-way ANOVA | kmeans clusters | F(3.15.601) | 0.554 | 0.649 | ns |
| **Supplementary 7B.Corticosterone 1h30 post-FST in males (z-score) top** | **CTL = 18, RES = 7, ANX = 12, ANH = 6** | | Shapiro Wilk on residuals | | | | | qqplot ok |
| | | | Levenes equality of variances | | F(3.39) | 2.133 | 0.112 | Equality of variances |
| | | | One-way ANOVA | kmeans clusters | F(3.39) | 3.707 | 0.019* | * |
| | | | Tukey post-hoc test | CTL vs ANX | t | 1.307 | 0.564 | ns |
| | | | Tukey post-hoc test | CTL vs RES | t | -1.42 | 0.495 | ns |
| | | | Tukey post-hoc test | CTL vs AHN | t | -2.114 | 0.167 | ns |
| | | | Tukey post-hoc test | ANX vs RES | t | -2.354 | 0.103 | ns |
| | | | Tukey post-hoc test | ANX vs ANH | t | -2.967 | 0.025* | * |
| | | | Tukey post-hoc test | RES vs ANH | t | -0.654 | 0.913 | ns |
| **Supplementary 7B.Corticosterone 1h30 post-FST in females (z-score) bottom** | **CTL = 18, RES = 5, ANX = 6, ANH = 10** | | Shapiro Wilk on residuals | | | | | qqplot ok |
| | | | Levenes equality of variances | | F(3.35) | 0.506 | 0.68 | Equality of variances |
| | | | One-way ANOVA | kmeans clusters | F(3.35) | 2.063 | 0.123 | ns |
| **Supplementary 7C. TH+cfos+/µm2 z-score in males (z-score) top** | **CTL = 18, RES = 7, ANX = 12, ANH = 6** | | Test of Sphericity | LC level | Mauchly's W<br>$\chi^2(5)$<br>=15.403(5) | 0.665 | 0.009** | Greenhouse-Geisser correction 0.806 |
| | | *Within -subjects effect* | repeated measures ANOVA | LC level | F(2.417.94.270 | 1.54 | 0.216 | ns |
| | | | repeated measures ANOVA | LC level * Condition | F(7.252.94.270 | 1.37 | 0.225 | ns |
| | | *Between-subjects effect* | repeated measures ANOVA | Condition | F(3.39) | 3.143 | 0.036* | * |
| | | | Bonferonni post-hoc test | CTL vs ANX | t | -3.068 | 0.023* | * |
| | | | Bonferonni post-hoc test | CTL vs RES | t | -0.953 | 1 | ns |
| | | | Bonferonni post-hoc test | CTL vs ANH | t | -0.876 | 1 | ns |
| | | | Bonferonni post-hoc test | ANX vs RES | t | 1.512 | 0.693 | ns |
| | | | Bonferonni post-hoc test | ANX vs ANH | t | 1.461 | 0.693 | ns |
| | | | Bonferonni post-hoc test | RES vs ANH | t | 0.021 | 1 | ns |
| **Fig.7C. TH+cfos+/µm2 z-score in females (z-score) bottom** | **CTL = 18, RES = 5, ANX = 6, ANH = 10** | | Test of Sphericity | LC level | Mauchly's W<br>$\chi^2(5)$<br>=13.816(5) | 0.664 | 0.017* | Sphericity violated |
| | | *Within -subjects effect* | repeated measures ANOVA | LC level | F(2.503.87.597 | 5.004 | 0.005** | ** |
| | | | repeated measures ANOVA | LC level * Condition | F(7.508.87.597 | 1.961 | 0.065 | ns |
| | | *Between-subjects effect* | repeated measures ANOVA | Condition | F(3.35) | 1.312 | 0.286 | ns |
| **Supplementary 7D.TH-cfos+/µm2 in males (z-score) top** | **CTL = 18, RES = 7, ANX = 12, ANH = 6** | | Shapiro Wilk on residuals | | | | | qqplot ok |
| | | | Test of Sphericity | LC level | Mauchly's W<br>$\chi^2(5)$<br>=30.835(5) | 0.442 | <0.001 | Greenhouse-Geisser correction 0.673 |
| | | *Within -subjects effect* | repeated measures ANOVA | LC level | F(2.019.78.742 | 1.433 | 0.245 | ns |
| | | | repeated measures ANOVA | LC level * kmeans cluster | F(6.057.78.742 | 2.303 | 0.042* | * |
| | | *Between-subjects effect* | repeated measures ANOVA | kmeans cluster | F(3.39) | 1.042 | 0.385 | ns |
| | | | Bonferonni post-hoc test LC level* kmeans cluster, conditional on kmeans | ANH — RD vs RV | t | -2.779 | 0.05 | ns |
| | | | Bonferonni post-hoc test LC level* kmeans cluster, conditional on kmeans | ANH — RD vs CD | t | -3.163 | 0.018* | * |
| | | | Bonferonni post-hoc test LC level* kmeans cluster, conditional on kmeans | ANH — RD vs CV | t | -1.975 | 0.332 | ns |
| **Supplementary 7D.TH-cfos+/µm2 in females (z-score) bottom** | **CTL = 18, RES = 5, ANX = 6, ANH = 10** | | Shapiro Wilk on residuals | | | | | qqplot ok |
| | | | Test of Sphericity | LC level | Mauchly's W<br>$\chi^2(5)$<br>=16.530(5) | 0.613 | 0.006** | Greenhouse-Geisser correction 0.810 |
| | | *Within -subjects effect* | repeated measures ANOVA | LC level | F(2.429, 85.007) | 1.734 | 0.176 | ns |
| | | | repeated measures ANOVA | LC level * kmeans cluster | F(7.286, 85.007) | 1.717 | 0.113 | ns |
| | | *Between-subjects effect* | repeated measures ANOVA | kmeans cluster | F(3.35) | 0.504 | 0.682 | ns |
